# Perisynaptic astroglial response to *in vivo* long-term potentiation and concurrent long-term depression in the hippocampal dentate gyrus

**DOI:** 10.1101/2025.05.13.653827

**Authors:** Andrea J. Nam, Masaaki Kuwajima, Patrick H. Parker, Jared B. Bowden, Wickliffe C. Abraham, Kristen M. Harris

## Abstract

Perisynaptic astroglia provide critical molecular and structural support to regulate synaptic transmission and plasticity in the nanodomain of the axon-spine interface. Three-dimensional reconstruction from serial section electron microscopy (3DEM) was used to investigate relationships between perisynaptic astroglia and dendritic spine synapses undergoing plasticity in the hippocampus of awake adult male rats. Delta-burst stimulation (DBS) of the medial perforant pathway induced long-term potentiation (LTP) in the middle molecular layer and concurrent long-term depression (cLTD) in the outer molecular layer of the dentate gyrus. The contralateral hippocampus received baseline stimulation as a within-animal control. Brains were obtained 30 minutes or 2 hours after DBS onset. An automated 3DEM pipeline was developed to enable unbiased quantification of astroglial coverage at the perimeter of the axon-spine interface. Under all conditions, >85% of synapses had perisynaptic astroglia processes within 120 nm of some portion of the perimeter. LTP broadened the distribution of spine sizes while reducing the presence and proximity of perisynaptic astroglia near the axon-spine interface of large spines. In contrast, cLTD transiently reduced the length of the axon-spine interface perimeter without substantially altering astroglial apposition. The postsynaptic density was discovered to be displaced from the center of the axon-spine interface, with this offset increasing during LTP and decreasing during cLTD. Astroglial access to the postsynaptic density was diminished during LTP and enhanced during cLTD, in parallel with changes in spine size. Thus, access of perisynaptic astroglia to synapses is dynamically modulated during LTP and cLTD alongside synaptic remodeling.

**Significance Statement:** Perisynaptic astroglia provide critical molecular and structural regulation of synaptic plasticity underlying learning and memory. The hippocampal dentate gyrus, a brain region crucial for learning and memory, was found to have perisynaptic astroglia at the axon-spine interface of >85% of excitatory synapses measured. Long-term potentiation triggered the retraction of perisynaptic astroglia processes selectively from large synapses. This retraction decreased access of perisynaptic astroglia to the postsynaptic density, which was discovered to be located off-center in the axon-spine interface. Concurrent long-term depression temporarily (< 2 h) decreased spine perimeter and thus increased access of synapses to perisynaptic astroglia. These findings provide new insights into how the structural dynamics of spines and synapses shape access to perisynaptic astroglia.

## Introduction

Astroglia are complex cells that play essential roles in synaptic information processing (Semyanov and Verkhratsky, 2021). During development, they shape neuronal connectivity by participating in synapse formation and pruning (Risher et al., 2014; Chung et al., 2015; Allen and Eroglu, 2017; Lee et al., 2021; Tan et al., 2021; Saint-Martin and Goda, 2023). In the adult brain, astroglia support synaptic communication by maintaining ion homeostasis, removing excess glutamate, and supplying glutamine to neurons (Allen and Eroglu, 2017; Saint-Martin and Goda, 2023). Astroglial N-methyl-D-aspartate (NMDA) receptors regulate presynaptic strength (Letellier et al., 2016; Chipman et al., 2021), while calcium elevations (Shigetomi et al., 2013; Bindocci et al., 2017; Arizono et al., 2020) trigger the release of gliotransmitters, (including glutamate, ATP, and D-serine, that modulate synaptic activity) (Sahlender et al., 2014; Bazargani and Attwell, 2016; Fiacco and McCarthy, 2018; Lim et al., 2021; Letellier and Goda, 2023). These mechanisms are critical for synaptic function and position astroglia as key contributors in long-term potentiation (LTP) (Henneberger et al., 2010; Liu et al., 2022) and long-term depression (LTD) (Durkee et al., 2021)—cellular processes widely considered to underlie learning and memory.

Astroglial influence on synaptic activity is shaped by their structural relationship with pre- and postsynaptic components (Saint-Martin and Goda, 2023). Their highly branched morphology includes perisynaptic astroglial processes (PAPs), which comprise over 60% of the astroglial volume (Aboufares El Alaoui et al., 2021; Salmon et al., 2023). PAPs predominantly lie beyond the diffraction limit of conventional light microscopy (Rusakov, 2015) and display alternating constrictions and expansions (Salmon et al., 2023) that compartmentalize intracellular calcium transients (Bindocci et al., 2017; Arizono et al., 2020; Denizot et al., 2022). Computational modeling indicates that PAP proximity impacts ionic homeostasis and rates of extracellular neurotransmitter diffusion (Kinney et al., 2013; Toman et al., 2023). Hence, a comprehensive analysis of astroglia-neuron structural relationships is essential for understanding the functional roles of astroglia during LTP and LTD.

The structural relationship between astroglia and neurons is dynamic over time and heterogeneous across brain regions. PAPs exhibit spontaneous motility and can undergo morphological changes in response to circuit-wide perturbations in neuronal activity (Hirrlinger et al., 2004; Bernardinelli et al., 2014; Perez-Alvarez et al., 2014). For example, in the hippocampal CA1 region, PAPs retract from excitatory synapses during LTP (Henneberger et al., 2020), whereas whisker stimulation increases astroglial coverage at synapses in the mouse somatosensory cortex (Genoud et al., 2006).

Connectomic analyses suggest that astroglial coverage decreases at small, same-axon-same-dendrite synapse pairs with low size variance—features indicative of LTD (Yener et al., 2025). In parallel, single-cell RNA sequencing studies have identified 5-7 subtypes of protoplasmic astroglia across the hippocampus, striatum, and cortex (Batiuk et al., 2020; Endo et al., 2022). Marked morphological and molecular differences distinguish astroglia across cortical layers (Lanjakornsiripan et al., 2018). Thus, clarifying astroglial contributions to learning and memory also requires examining their structural relationships with neurons during synaptic plasticity in different brain areas.

In this study, we investigated astroglial apposition at the axon-spine interface (ASI) during LTP and LTD in the dentate gyrus—a hippocampal region critical for pattern separation and episodic memory, and one of the few areas where adult neurogenesis persists (Aimone et al., 2011; Hainmueller and Bartos, 2020; Denoth-Lippuner and Jessberger, 2021). Previous work using randomly selected photomicrographs demonstrated that astroglial coverage of synapses increases during LTP in the dentate gyrus molecular layer (Wenzel et al., 1991). However, more recent studies have revealed that key features of astroglial nanostructure are lost with even moderate sectioning intervals in electron microscopy (Salmon et al., 2023). To overcome this limitation, we employed three-dimensional reconstruction from serial section electron microscopy (3DEM) coupled with a novel, automated method to measure PAP apposition at the ASI perimeter. Our results reveal that in the dentate gyrus, astroglial processes selectively withdraw from the ASI of large synapses during LTP, while maintaining close apposition to most synapses during LTD, regardless of synapse size.

## Materials and Methods

### Surgery

Data were collected from six adult male Long-Evans rats aged 121-185 days. In the experimental hemisphere of each animal, wire stimulating electrodes were surgically implanted into the medial and lateral perforant pathways of the angular bundle. In the contralateral control hemisphere, a single stimulating electrode was implanted into the medial perforant pathway only. Bilateral wire electrodes were also implanted into the dentate gyrus hilus to record field excitatory postsynaptic potentials (fEPSP). Full details of the surgical procedures can be found in Bowden et al. (2012).

### Electrophysiology

Two weeks following surgery, 30-minute-long baseline recording sessions were commenced and carried out every two days until a stable baseline was achieved. Baseline recording sessions (which were conducted during the animals’ dark cycle and while the animals were in a quiet, alert state) consisted of constant-current biphasic square-wave test pulses (150 µs half-wave duration) delivered at a rate of 1 per 30 seconds. Test pulses were administered alternating between the three stimulating electrodes. Test pulse intensity was set to evoke medial path waveforms with fEPSP slopes ≥3.5 mV/ms in association with population spike amplitudes between 2 and 4 mV, at a stimulation current ≤500 µA. Once baseline recordings stabilized and following 30 minutes of test pulses, delta-burst stimulation (DBS) was delivered to the ipsilateral medial perforant path of the experimental hemisphere to induce LTP in the middle and cLTD in the outer molecular layer of the dentate gyrus (Bowden et al., 2012). The DBS protocol consisted of five trains of 10 pulses (250 µs half-wave duration) delivered at 400 Hz at a 1 Hz inter-train frequency, repeated 10 times at 1-minute intervals. The contralateral hemisphere received test pulses only to serve as within-subject control recordings. Following DBS, test pulse stimulation was resumed until the animal was sacrificed at either 30 min or 2 h following DBS onset depending on the experimental group (3 animals per group). The initial slopes of the medial and lateral path fEPSPs were measured for each waveform and expressed as a percentage of the average response during the last 15 min of recording before DBS (Fig. 1A-C).

**Figure 1.**
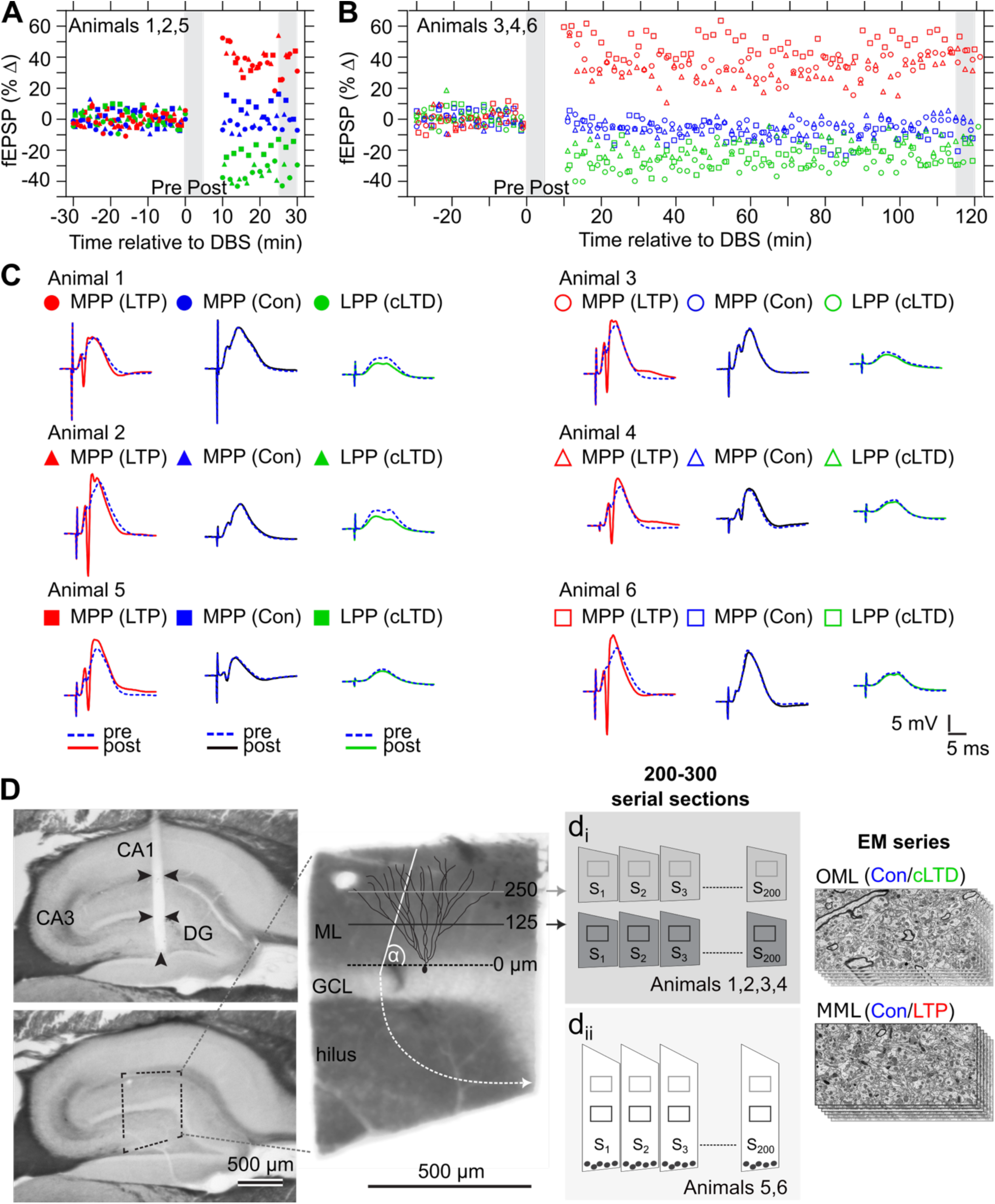
LTP and cLTD outcomes and preparation of dentate gyrus MML and OML tissue for 3DEM. **A-D**: Experimental hemisphere medial perforant pathway (MPP) responses (red, LTP), control hemisphere MPP responses (blue, control), and experimental hemisphere lateral perforant pathway (LPP) responses (green, cLTD). **(A)** Graphs of percent change in average fEPSP response relative to baseline for (A) animals 1, 2, and 5 sacrificed 30 minutes (min) after DBS stimulation (delivered over t = 0-10 mins), and **(B)** animals 3, 4, and 6 sacrificed 2 hours (h) after DBS stimulation (delivered over t = 0-10 mins). **(C)** Representative smoothed waveforms obtained during the times shown as gray vertical bars in the graphs of A and B for animals 1-6. Pre-DBS baseline responses (blue, dotted) are superimposed by post-DBS responses (solid). Scale bars: 5 mV (vertical), 5 ms (horizontal). **(D)** (Left) Example parasagittal hippocampal tissue section with visible tract from recording electrode positioned in dentate gyrus hilus (top) and adjacent tissue section used for EM series preparation (bottom). (Middle) Region of dentate gyrus molecular layer isolated for ultra-thin serial sectioning. Horizontal lines indicate sectioning planes: OML (grey) and MML (black) at 250 µm and 125 µm from top of granule cell layer, respectively. OML and MML were sectioned separately for animals 1-4 (d_i_), and on the same tissue section for animals 5 and 6 (white sectioning plane, ⍺ = 26.6°, d_ii_). (Right) Stacks of 200-300 serial section EM images were obtained in the OML (top) and MML (bottom) using transmission electron microscopy or scanning electron microscopy operating in the transmission mode. Scale bars: 500 µm.

### Perfusion and Fixation

Animals were perfusion-fixed under halothane anesthesia and a tracheal supply of oxygen (Kuwajima et al., 2013a). The perfusion protocol consisted of a brief (∼20 s) wash with oxygenated Krebs-Ringer Carbicarb buffer (concentration (in mM): 2.0 CaCl_2_, 11.0 D-glucose, 4.7 KCl, 4.0 MgSO_4_, 118 NaCl, 12.5 Na_2_CO_3_, 12.5 NaHCO_3_; pH 7.4; Osmolality: 300-330 mmol/kg), followed by 2% formaldehyde and 2.5% glutaraldehyde in 0.1 M cacodylate buffer (pH 7.4) containing 2 mM CaCl_2_ and 4 mM MgSO_4_ for approximately 1 hour (∼2 L of fixative per animal). Brains were removed from the skull around 1-hour post-perfusion, wrapped in layers of cotton gauze, and then shipped by overnight delivery (TNT Holdings B.V.) in the same fixative from the Abraham Laboratory in Dunedin, New Zealand to the Harris Laboratory in Austin, Texas.

### Tissue Processing and Serial Sectioning

The fixed tissue was sliced using a vibrating blade microtome (Leica Microsystems) along the parasagittal plane into 70 µm thick slices (Fig. 1D). The slice containing the recording electrode and its two neighboring slices were processed for electron microscopy as described previously (Harris et al., 2006; Kuwajima et al., 2013a; Bromer et al., 2018). In summary, the tissue was treated with reduced osmium (1% osmium tetroxide and 1.5% potassium ferrocyanide in 0.1 M cacodylate buffer with 2 mM Ca^2+^ and 4 mM Mg^2+^) followed by microwave-assisted incubation in 1% osmium tetroxide under vacuum. Next the tissue was subjected to microwave-assisted dehydration and en bloc staining with 1% uranyl acetate in ascending concentrations of ethanol. The dehydrated tissue was embedded into LX-112 epoxy resin (Ladd Research) at 60°C for 48 h. Then the tissue-containing resin blocks were cut into ultra-thin sections at the nominal thickness of 45 nm with a 35° diamond knife (DiATOME) on an ultramicrotome (Leica Microsystems). For 4 of the 6 animals, the MML and OML regions were sectioned ∼125 µm and ∼250 µm from the top of the granule cell layer in the dorsal blade of the hippocampal dentate gyrus (Fig. 1D, d_i_). For the remaining 2 animals, the tissue was sectioned at a 26.6° angle relative to the granule cell layer, allowing many MML and OML dendrites to be cut in cross-section and to be captured within the same set of ultra-thin serial sections (Fig. 1D, d_ii_). The ultra-thin tissue sections were collected onto Synaptek Be-Cu slot grids (Electron Microscopy Sciences or Ted Pella), coated with Pioloform (Ted Pella), and finally stained with a saturated aqueous solution of uranyl acetate followed by lead citrate (Reynolds, 1963).

### Imaging

The ultra-thin serial tissue sections were imaged, blinded as to experimental condition (Fig. 1D). Tissue from 4 of the 6 animals was imaged with a JEOL JEM-1230 TEM to produce 16 EM image series (2 series per condition). Tissue from the remaining 2 animals was imaged with a transmission-mode scanning EM (tSEM) (Zeiss SUPRA 40 field-emission SEM with a retractable multimode transmitted electron detector and ATLAS package for large-field image acquisition), to produce 8 EM image series (1 series per condition) (Kuwajima et al., 2013a, 2013b). Sections imaged with TEM were captured in two field mosaics at 5,000x magnification with a Gatan UltraScan 4000 CCD camera (4,080 pixels × 4,080 pixels) controlled by Digital Micrograph software (Gatan). These mosaics were then stitched together post-hoc using the Adobe Photoshop Photomerge function. On the tSEM, each section was imaged with the transmitted electron detector from a single field encompassing up to 32.768 µm × 32.768 μm (16,384 pixels × 16,384 pixels at 2 nm/pixel resolution). The scan beam dwell time was set to 1.3-1.4 ms and the accelerating voltage was set to 28 kV in high-current mode.

### Alignment

Serial TEM images were first manually aligned in legacy Reconstruct (Fiala, 2005). Then the initial round of automatic alignment for all image volumes was completed using the TrakEM2 Fiji plugin (Cardona et al., 2012; Saalfeld et al., 2012; Schindelin et al., 2012). Images underwent rigid alignment, followed by affine alignment, and then elastic alignment. These new alignments were applied permanently to the images. Next the TrakEM2 aligned image volumes were automatically aligned again using AlignEM SWiFT, a software that aligns serial section images using Signal Whitening Fourier Transform Image Registration (SWiFT-IR) (Wetzel et al., 2016). Finally, image volumes were subjected to further regional, by-dendrite alignments to overcome local artifacts (stretches, tears, or folds) using Reconstruct’s modern replacement PyReconstruct, an open-source Python-based software for serial EM analysis (https://github.com/SynapseWeb/PyReconstruct). All image series in the dataset were given a five-letter code to blind investigators as to the experimental condition. The grating replica (0.463 μm per square; Ernest Fullam, Inc., Latham, New York) image was acquired along with serial sections and used to calibrate pixel size for each series. In addition, the section thickness was estimated using the cylindrical diameters method and found to be 42-55 nm, close to the nominal setting (45 nm) on the ultramicrotome (Fiala and Harris, 2001a).

### Unbiased Annotation of Tripartite Synapses

Previous work has demonstrated that the average number of dendrite microtubules scales linearly with dendrite diameter and the number of spines per micron length of dendrite (Fiala and Harris, 2001b; Harris et al., 2022). Therefore, in each image volume from every layer of the dentate gyrus in each animal, three dendrites of comparable caliber were chosen based on the average microtubule count for that layer (average microtubule count per dendrite: OML – 20-25; MML – 30-35). A total of 72 dendrites and 2,083 excitatory synapses were analyzed for the 24 series in this dataset.

Object annotations were completed while masked to condition using legacy Reconstruct (Fiala, 2005) and PyReconstruct. Annotation files (.ser) created using legacy Reconstruct were updated to the PyReconstruct file format (.jser). Dendritic spine contours were manually traced from each selected dendrite. Postsynaptic densities (PSD) of excitatory synapses were identified based on their asymmetric, high-contrast electron densities and the presence of clear, round presynaptic vesicles in the presynaptic axonal bouton (Harris and Weinberg, 2012). The PSD area was measured in PyReconstruct according to the specific sectioning orientation, as previously described in Harris et al. (Harris et al., 2015). PAPs, characterized by their tortuous morphology, relatively clear cytoplasm (Fig. 2A), and glycogen granules on serial sections (Witcher et al., 2007), were traced if they fell within 1 µm of a synaptic profile center. Given that synapses can interact with all astroglial compartments (Aten et al., 2022), the annotated PAPs consisted of both thick astroglia branches and terminal leaflets (Semyanov and Verkhratsky, 2021; Salmon et al., 2023). Along the z-axis, both PAPs and presynaptic axons were traced throughout the range of sections where the ASI was visible, plus an additional 1-2 sections beyond the ASI in both directions. In the x-y plane, axons were traced until the bouton tapered into a vesicle-free region.

**Figure 2.**
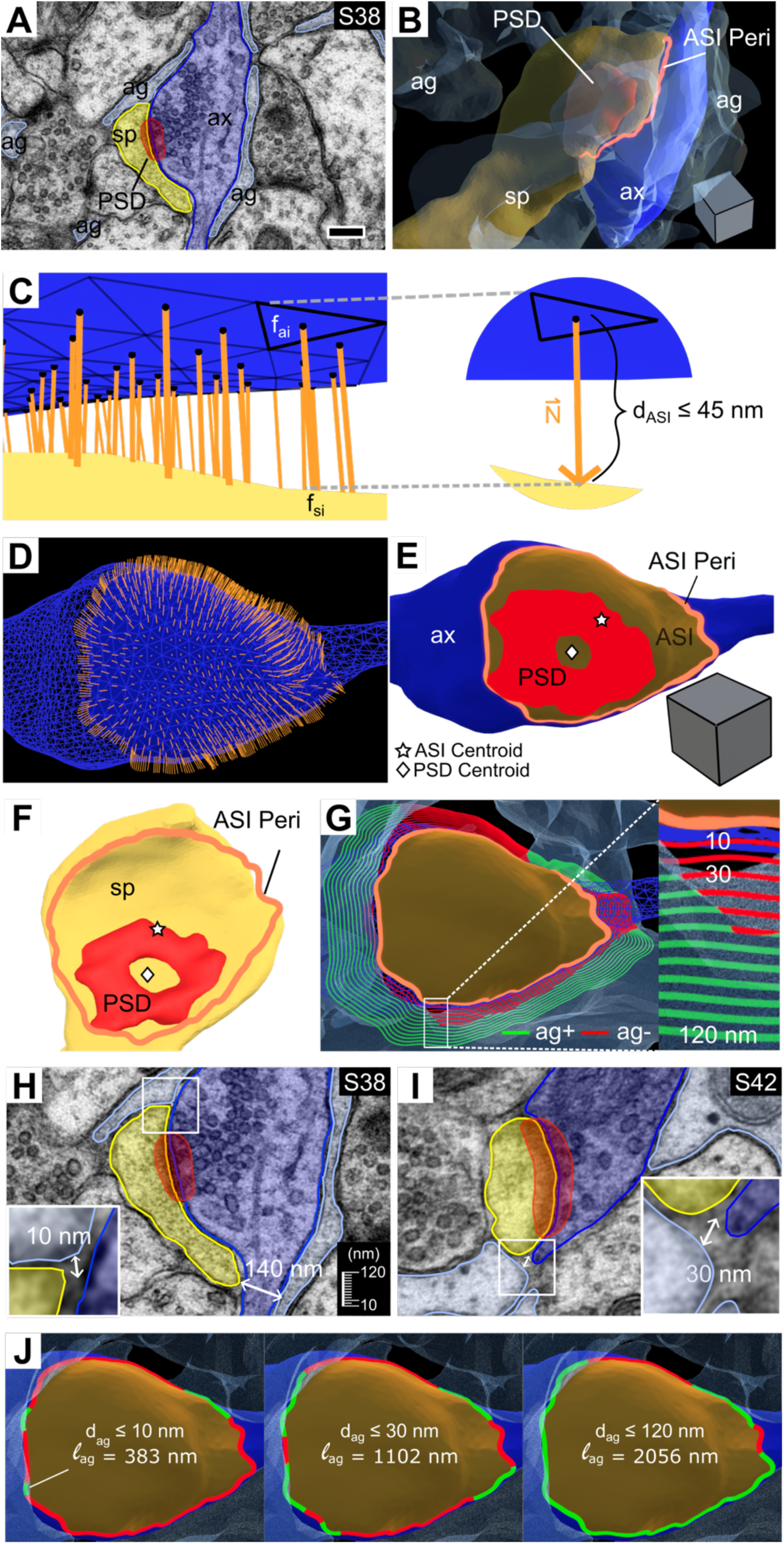
Automated ASI detection and measurement of astroglia apposition at the ASI perimeter. **(A)** EM image of serial section (S) 38 through a tripartite synapse. **(B)** 3D reconstruction showing the postsynaptic dendritic spine (sp, yellow), presynaptic axon (ax, dark blue), perisynaptic astroglia (ag, light blue), and postsynaptic density (PSD, red), with the ASI perimeter (peri) outlined (orange). **(C)** Detection of ASI facets (black triangles, f_ai_) on the axon mesh surface with normal vector projections (orange lines/arrow) from the center of each triangular mesh facet (black dot) to the intersecting spine mesh facet (f_si_). (Right) Zoomed-in view of an example ASI face highlighted with bold black edges and maximum d_ASI_. **(D)** ASI axon facets with corresponding normal vectors (orange). **(E)** Relative location of PSD within the ASI (brown) projected onto the axon surface and **(F)** the spine surface. The computed ASI perimeter, ASI centroid (white star), and PSD centroid (white diamond) are shown. For this synapse the centroid ends up in the middle of the perforation. **(G)** Concentric rings around the ASI perimeter indicate the Euclidean distances between the astroglia and ASI perimeter (d_ag_). Rings are color-coded by astroglia presence (ag+, green) or absence (ag-, red) within a particular d_ag_. The axon and surrounding astroglia are also shown. (Inset) Zoomed-in view of distance thresholds (d_ag_ ≤ 10-120 nm) tested. **(H)** Zoomed-in view of EM image from A, showing astroglia positioned 10 nm from the ASI perimeter at one edge (d_ag_ = 10 nm) but 140 nm away at the opposite edge (d_ag_>120 nm, exceeding the maximum distance threshold tested), with an intervening axon. **(I)** EM image of a different synapse on the same dendrite as in A, observed on S42, depicting d_ag_ = 30 nm. In H and I, insets show an enlarged view of the regions indicated by white boxes. **(J)** Length of the ASI perimeter apposed (ag+, green) and not apposed (ag-, red) by astroglia based on d_ag_ ≤ 10 nm (left), 30 nm (middle), and 120 nm (right). The axon and surrounding astroglia are also shown. The same synapse is shown in A-H and J, with consistent color coding applied across all panels. Scale bars, scale cube edge lengths: 250 nm.

Rare dually innervated spines (with both excitatory and inhibitory synapses) were excluded from analyses (Villa et al., 2016; Kleinjan et al., 2023). Likewise, synapses were omitted if local section flaws or proximity to image volume boundaries prevented accurate and complete PAP segmentations.

### 3D Reconstruction: Mesh Generation and Processing

3D watertight mesh analysis was completed using Blender, a free, open-source computer graphics tool with a Python interface. Spine and axon objects were reconstructed from serial section traces using Neuropil Tools (https://github.com/mcellteam/neuropil_tools), a Blender add-on, in the MCell/CellBlender v4.0.6 bundle for Blender 2.93. Any incorrectly meshed objects were manually fixed using native Blender mesh editing tools before being exported in the Polygon File Format (PLY) for 3D objects. Exported meshes were re-meshed using the isotropic re-meshing function within the Computational Geometry Algorithms Library (CGAL) 5.0.2 Polygon Mesh Processing package (https://www.cgal.org). This tool converts faulty meshes into manifold and watertight meshes with outward-facing normal vectors. For the re-meshing routine, the number of iterations was set to 3 and the target edge length parameter was set to 0.04. All object meshes were then smoothed using GAMer2, a 3D mesh processing software that conditions surface meshes to correct for artifacts such as jagged boundaries and high aspect ratio faces (Lee et al., 2020). All axon meshes met a face density threshold of at least 10,000 faces per volume (Fig. 2B). The spine volumes were estimated in Blender using the Blender add-on, 3D Print Toolbox (https://extensions.blender.org/add-ons/print3d-toolbox).

PAPs and synapses were reconstructed using PyReconstruct, which uses trimesh to generate triangulated meshes with watertight surfaces (https://trimesh.org). Synapses were reconstructed from contact traces, drawn around each PSD area trace, and extended just beyond the spine membrane contour to enable visualization (Fig. 2A). All astroglial and synapse meshes were imported into Blender in the correct by-dendrite alignment as PLY files.

Linear tissue shrinkage can occur with chemical fixation for electron microscopy, which can be accounted for by the loss of extracellular space (Kalimo, 1976; Kirov et al., 1999; Kinney et al., 2013; Korogod et al., 2015; Tønnesen et al., 2018). However, all tissue samples were prepared in the same way, and it is not possible to accurately account for potential variations from the *in vivo* state. Therefore, the 3D reconstructions were evaluated without arbitrary correction.

### Automated 3D Analysis of Astroglial Apposition at the ASI Perimeter

An automated analysis pipeline was developed to measure astroglial apposition at the ASI perimeter. The algorithm was developed in Blender 3.6.5, using Blender’s Python API.

### Stage 1: ASI Identification

Since the axon mesh was more uniformly shaped than the spine mesh, the ASI was identified from the axon mesh faces based on two criteria (Fig. 2C, D). First, the projected normal vector from a given axon face (f_ai_) had to intersect a face (f_si_) on the apposing spine mesh. Second, the Euclidean distance between the f_ai_ center and apposing f_si_ center had to be less than a pre-determined maximum distance threshold (d_ASI_). For this study, d_ASI_ ≤ 45 nm was used (Fig. 2C).

### Stage 2: Measurement of the ASI area and Perimeter

The set of edges surrounding the triangulated ASI faces was extracted (Fig. 2E, F). Then the Relax function from the Loop Tools Blender add-on (https://extensions.blender.org/add-ons/looptools/) was used to smooth the boundary loop of edges over 10 iterations. The ASI perimeter was calculated by summing the lengths of each edge comprising the final ASI boundary (Fig. 2D-F). Similarly, the ASI area was determined by summing the areas of each face enclosed by the final ASI boundary.

### Stage 3: Measurement of Astroglial apposition at the ASI perimeter

The Euclidean distance from each ASI boundary edge midpoint to the nearest point on the PAP mesh (d_ag_) was measured. Then the lengths of all ASI edges with d_ag_ less than or equal to a pre-determined distance threshold were summed to give the length of the ASI perimeter surrounded by astroglia (*l*ag) (Fig. 2G). Astroglial proximity to the ASI perimeter was quantified by averaging d_ag_ across all ASI perimeter edges with d_ag_ ≤ 120 nm. The upper limit of d_ag_ ≤ 120 nm was chosen to ensure that the PAPs had an unobstructed extracellular diffusion path to the ASI perimeter (Fig. 2H, I).

For this study, *l*ag measurements were made based on distance thresholds ranging from d_ag_ ≤ 10 to 120 nm, in 10 nm increments (Fig. 2G, J), balancing spatial resolution limitations and functional relevance. With an x-y resolution of ∼2 nm per pixel and an axial resolution of ∼45 nm, a lower limit of d_ag_ ≤ 10 nm minimized anisotropy-related errors.

### Categorizing the Location of Astroglial Apposition

Synapses with astroglia within the 120 nm (upper limit discussed above) of the ASI perimeter were assigned to the ASI^ag+^ category. All remaining synapses (ASI^ag-^) were categorized based on astroglial apposition present at both pre- and postsynaptic elements but not at the ASI (Pre-Post), only at the presynaptic bouton (Pre), only at the postsynaptic spine (Post), or absent from all these locations (None). This subsequent classification was determined by examining the serial EM images to identify astroglial contact with the postsynaptic spine, the presynaptic bouton, or both.

### Measurement of PSD Offset

For each synapse, the ASI faces were duplicated, separated from the axon object, and converted into a new ASI surface mesh. Each PSD object was also duplicated and conformed to the axon surface using Blender’s native shrink-wrap mesh modifier (ax-PSD object). Then the geometric centroids of the ASI surface and ax-PSD meshes were calculated by averaging the coordinates of their respective vertices. PSD offset was defined as the Euclidean distance between these centroids.

For ASI^ag+^ synapses only, the average distance from astroglial apposition at the ASI to the nearest PSD edge (d_ag-PSD_) was estimated by averaging the Euclidean distances from each ASI perimeter edge midpoint with d_ag_ ≤ 120 nm to the nearest point on the ax-PSD object. Meanwhile, the overall ASI perimeter-to-PSD distance (d_ASI-PSD_) was calculated by averaging the Euclidean distances from all ASI perimeter edge midpoints to the nearest point on the ax-PSD object. Synapses were classified as ag-proximal (d_ag-PSD_ < d_ASI-PSD_) or ag-distal (d_ag-PSD_ ≥ d_ASI-PSD_). Additionally, the ratio of d_ag-PSD_ to d_ASI-PSD_ served as an alternative measure of PSD offset, incorporating directional information relative to astroglial apposition at the ASI.

### Unit Conversion

For each EM image series, a reference circle with a known, calibrated radius of 1 µm was stamped in PyReconstruct and imported into Blender as a two-dimensional object. The reference circle’s radius was measured in Blender (r_b_), and all subsequent Blender measurements were multiplied by a factor of 1 µm/r_b_ to ensure accurate scaling to microns.

### Experimental Design

As described above for Fig. 1, 3 animals were prepared for serial EM analysis at each of the two time points after the induction of LTP and cLTD. For each time point and condition, 3 dendrites of comparable caliber were selected for 3D reconstruction and analysis. To avoid introducing variance related to the correlation between spine density and dendrite caliber, we used the unbiased dendritic length (length of the dendritic segment extending from the first spine origin to the beginning of the last spine origin) to sample dendrites instead of randomly sampling the neuropil (Fiala and Harris, 2001b). To assess differences between LTP and control or cLTD and control, a mixed model was used with random effects for animal and dendrite included to account for inter-animal and inter-dendrite variability.

### Statistical Analyses

All statistical analyses and graphical plots were completed in RStudio using R version 4.4.1. All kernel density estimate (KDE) plots showing the probability density function (PDF) for parameters of interest were generated using geom_density() (ggplot2 package in R) with the default bandwidth calculated by stats::density()(adjust = 1).

Data clustering was conducted based on ASI perimeter and spine volume with conditions and time points pooled, but separately for MML and OML data. The silhouette method determines the optimal number of clusters (k) by measuring object similarity to its cluster compared to other clusters, with values ranging from −1 to 1 indicating bad to good data clustering. Iterating over k = 2 to 20, we determined that the optimal number of clusters was k = 2 for both layers using the silhouette function from the cluster package (version 2.1.6) in R. Then, we performed k-means clustering to separate the data into two clusters: cluster 1 (c1) and cluster 2 (c2). For the clustering algorithm, the maximum number of iterations allowed was set to 10 and the number of random sets to be chosen was set to 25.

Relative synapse percentages in each astroglia-ASI apposition category (ASI^ag+^/ASI^ag-^) or cluster category (c1/c2) were compared using Pearson’s Chi-squared (χ^2^) test with Yates’ continuity correction applied. Variable sample distributions were compared using a Kolmogorov-Smirnov (KS) test with a two-sided alternative hypothesis. Comparisons of means and linear regressions were conducted using the lmerTest package for R (Kuznetsova et al., 2017).

Linear mixed models (LMM) were fitted to the data with additive random intercepts for animal and dendrite included to account for variability at these group levels. To meet LMM assumptions of normality and homoscedasticity, the following data transformations were applied during analysis: log-transformations (applied to spine volume, average d_ag_, PSD area, ASI area, PSD offset, PSD-to-ASI area ratio, average d_ag-PSD_, and average d_ASI-PSD_ data) and square-root transformations (applied to ASI perimeter and *l*ag data).

Transformations of ASI perimeter data were applied only when ASI perimeter served as the response variable, not as a covariate predictor in the model. To limit data-based multicollinearity, numerical covariates were mean-centered. For comparisons across astroglia-ASI apposition categories, condition, clusters, PSD offset direction, and layer, the reference categories were defined as ASI^ag-^, control, cluster 1, ag-distal, and MML synapses, respectively. Model goodness-of-fit was assessed using the marginal coefficient of determination (R²), which represents the ratio of explained variance (Sum of Squares Regression, SSR) over total variance (Sum of Squares Total, SSTO). R^2^ was calculated using the r.squaredGLMM function from the MuMIn package in R. Effect sizes of individual predictors were quantified using partial eta² (η_p_^2^). All p-values from multiple comparisons were adjusted using the Benjamini-Hochberg (BH) procedure to control the False Discovery Rate (FDR).

The significance level was set at α = 0.05. Potential outliers were identified as values falling more than 1.5 times the interquartile range below the first quartile or above the third quartile. However, outliers were only excluded from analyses if manual inspection confirmed they were due to measurement errors.

## Code Accessibility

All code for the automated ASI detection and assessment of astroglial apposition is hosted on GitHub (code available at: https://github.com/ajnam03/asi_blender.git).

## Results

### Apposition of perisynaptic astroglial processes (PAPs) at MML synapses

MML synapses were reconstructed from serial EM images (Fig. 3A, B), and an automated 3D analysis method was developed to evaluate astroglial coverage within 10 to 120 nm of the ASI perimeter (Methods). Synapses were categorized based on the presence (ASI^ag+^) or absence (ASI^ag-^) of astroglia within 120 nm of the ASI perimeter.

**Figure 3.**
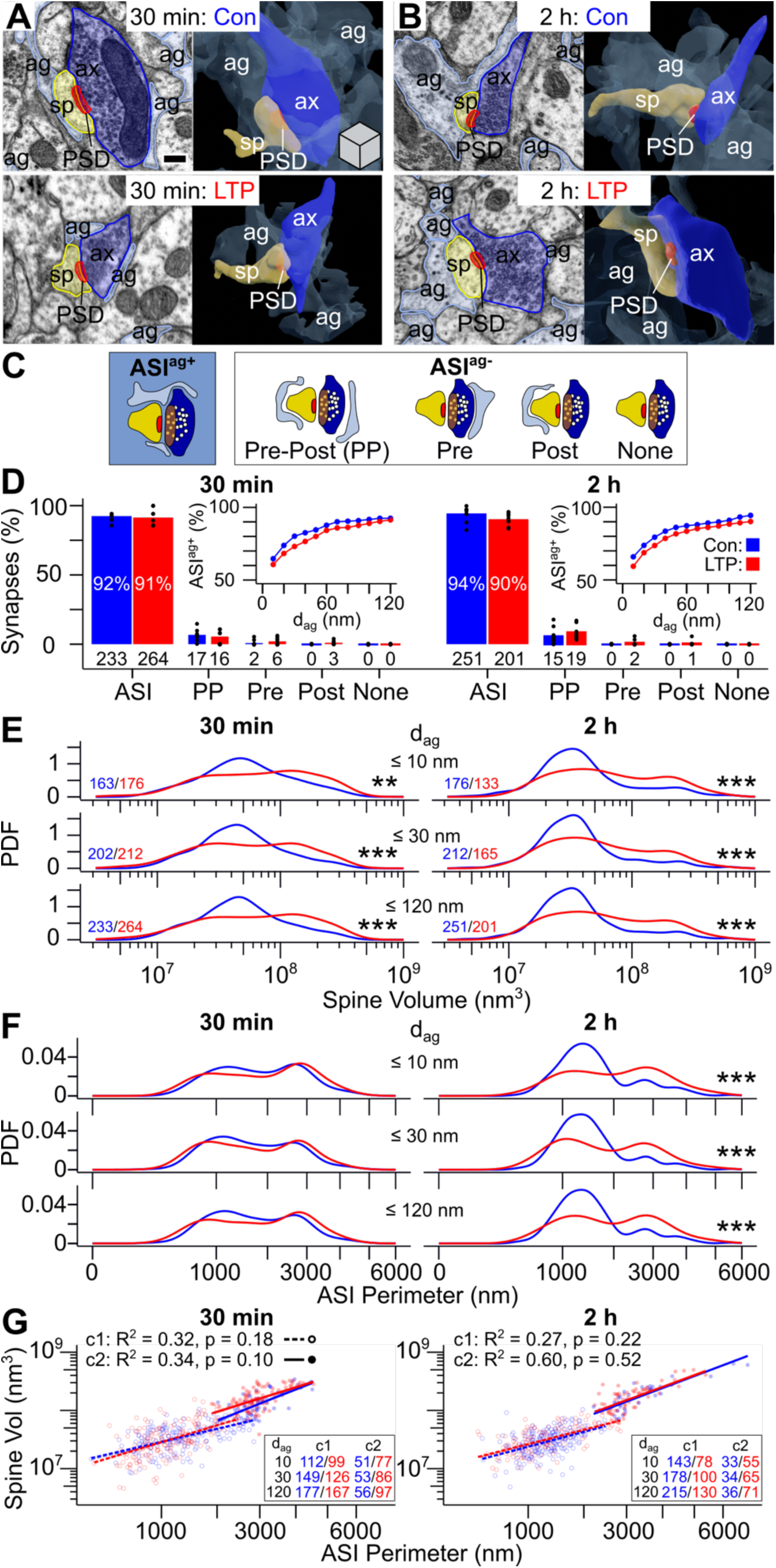
Nearly all MML spines have some perisynaptic astroglia and undergo a sustained increase in spine volume and ASI length during LTP relative to control stimulation. **(A)** EM image (left) and 3D reconstruction (right) of the spine (sp, yellow), postsynaptic density (PSD, red), presynaptic axon (ax, dark blue), and perisynaptic astroglia (ag, light blue) for an MML synapse during control (top) and LTP (bottom) at 30 min and **(B)** 2 h after LTP induction. **(C)** Synapse categories based on astroglial presence (ASI^ag+^) or absence (ASI^ag-^) within 120 nm of the ASI perimeter and specific astroglia location. The spine, axon, PSD, perisynaptic astroglia, presynaptic vesicles (small white circles), and ASI (brown oval) are represented using the same color scheme as A-B. **(D)** Bar graph of overall synapse percentage in each synapse category. Numbers below bars indicate synapse count. Dots represent dendrite-specific synapse percentages. (Insets) Line plots show ASI^ag+^ synapse percentages based on each d_ag_ threshold tested. **(E)** Kernel density estimate plots show the probability density functions (PDFs) for spine volume (log-scale axis) and **(F)** ASI perimeter (square-root axis) for synapse subsets based on d_ag_. Synapse counts color-coded by condition are shown in bottom left corner in E and apply to E-F (relatively few synapses in cluster 2 had d_ag_ ≤ 10 nm). **(G)** Regression plot of spine volume (log-scale y-axis) and ASI perimeter (square-root x-axis) for synapses in cluster 1 (c1) or cluster 2 (c2) with d_ag_ ≤ 10 (lowest opacity), 30 nm (medium opacity), and 120 nm (highest opacity). Raw data, R² values, and Benjamini-Hochberg (BH)-adjusted p-values (condition effect) for d_ag_ ≤ 120 nm are displayed. Synapse counts color-coded by condition are shown in bottom right corner. In D-G, data are from control (blue) and LTP (red) MML synapses at 30 min (left) and 2 h (right) during LTP. Asterisks indicate significant condition effects for synapse subsets defined by d_ag_ (BH-adjusted **p < 0.01, ***p < 0.001; see Table S1 for statistical details). Scale bar and cube (edge length = 250 nm) in A apply to A-B.

Then ASI^ag-^ synapses were further subdivided by the astroglial location beyond the ASI perimeter (Fig. 3C). The percentage of ASI^ag+^ synapses varied depending on the specific distance threshold used (Fig. 3D, insets). However, approximately 60% of MML synapses had astroglial apposition within 10 nm of the ASI, 70-80% had astroglial apposition within 30 nm, and more than 90% had astroglial apposition within 120 nm at both time-points in both control and LTP hemispheres (Fig. 3D). Thus, synapses with astroglial apposition within 10, 30, or 120 nm were assessed in all subsequent analyses. No synapses were entirely devoid of astroglial contact at all locations. These findings indicate that most MML synapses were tripartite at some portion of their ASI perimeter under control and LTP conditions.

### Enlargement of spines and expansion of ASI during LTP

At 30 minutes and 2 hours during LTP, a shift toward larger spine volumes was observed, regardless of whether astroglia occurred within 10, 30, or 120 nm of some portion of the ASI (Fig. 3E). Similarly, the ASI perimeter distribution consistently shifted rightward at 2 hours (Fig. 3F), and both the mean spine volume and ASI perimeter were significantly elevated at 2 hours compared to control (Fig. S1A, B). Interestingly, synapses with astroglial apposition had larger spine volumes and ASI perimeters than those without astroglial contact at 30 minutes in control and at both time points during LTP (Fig. S1A, B). Together, these results suggest that both spine volume and ASI perimeter—two measures of spine size—increased during LTP at MML tripartite synapses.

### K-means clustering based on spine volume and ASI perimeter in MML

It is notable that the spine volume and ASI perimeter distributions showed distinct bimodal shapes during LTP, suggesting that LTP could preferentially affect sub-populations of spines (Fig. 3E, F). Using the silhouette method and k-means clustering, two clusters of MML ASI^ag+^ synapses were identified based on spine volume and ASI perimeter (Fig. 3G). Cluster 2, which accounted for 27% of MML ASI^ag+^ synapses, had larger spine volumes and ASI perimeters than cluster 1 synapses (Table S1). The proportion of cluster 2 synapses significantly increased at 30 minutes and 2 hours during LTP. Spine volume and ASI perimeter were positively correlated across synapse clusters under both experimental conditions (Fig. 3G). Hence, in all subsequent work, regression analyses were done independently for these distinct clusters of small and large spines.

### Sustained spine enlargement increases PSD displacement from the ASI center during LTP

The PSD—an electron-dense region enriched with proteins critical for synaptic transmission and plasticity—was displaced from the ASI center by 47-57 nm under control conditions (Fig. 4A, B). During LTP, the distribution of PSD areas broadened and remained elevated for at least 2 hours (Fig. 4C), and the PSD’s offset increased significantly at 2 hours (Fig. 4D). Both PSD area and offset were positively correlated with ASI area, and after controlling for spine size, no direct effect of LTP on either measure was detected (Fig. 4E, F). In contrast, PSD area and offset showed only weak and inconsistent relationships with the extent of astroglial coverage (Fig. S3C, E), astroglial distance from the ASI perimeter (Fig. S3D, F), and with each other (Fig. S3G). These findings suggest that LTP increased the prevalence of both small and large synapses, and that the associated spine enlargement contributed to greater PSD offset from the ASI center.

**Figure 4.**
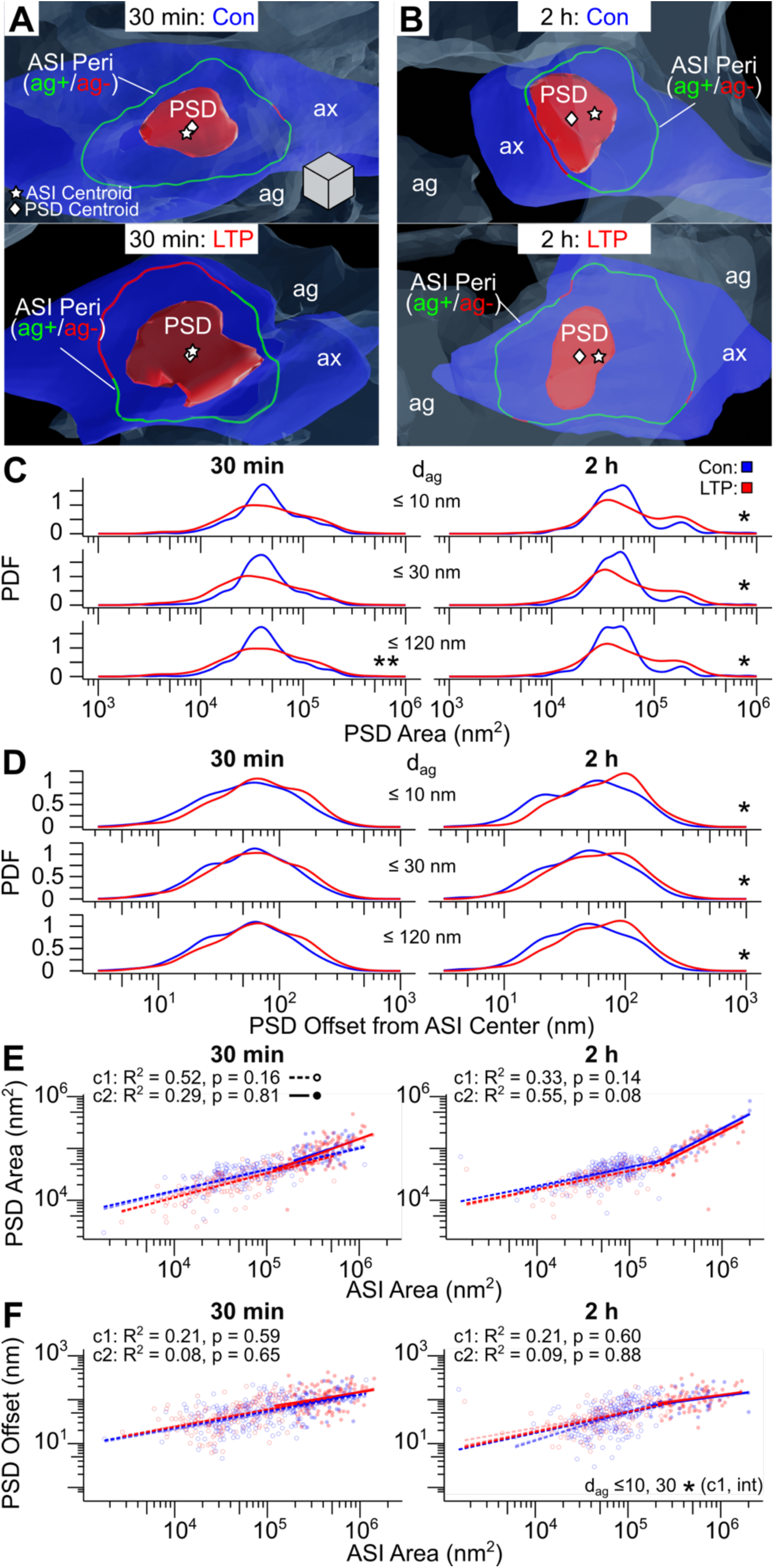
The PSD area range expands, and the PSD offset from the ASI center increases during LTP. **(A)** 3D reconstruction of the postsynaptic density (PSD, red) projected onto the axon mesh (ax, dark blue), perisynaptic astroglia (ag, light blue), ASI perimeter color-coded by presence (ag+, green) or absence (ag-, red) of astroglia within 120 nm, ASI centroid (white star), and PSD centroid (white diamond) for an MML synapse during control (top) and LTP (bottom) at 30 min and **(B)** 2 h following LTP induction. **(C)** Kernel density estimate (KDE) plots show the probability density function (PDF) for PSD area (log-scale axis) and **(D)** PSD offset from the ASI center (log-scale axis) for synapses subset based on d_ag_. **(E)** Regression plot of PSD area (log-scale y-axis) and **(F)** PSD offset from the ASI center (log-scale y-axis) versus ASI area (log-scale x-axis) for c1 or c2 synapses with d_ag_ ≤ 10 nm (lowest opacity lines), 30 nm (medium opacity lines), and 120 nm (highest opacity lines). Raw data, R² values, and Benjamini-Hochberg (BH)-adjusted p-values (condition effect) for d_ag_ ≤ 120 nm are displayed. In C-F, data are from control (blue) and LTP (red) MML synapses at 30 min (left) and 2 h (right) during LTP (see Figure 3G for synapse counts). Asterisks indicate significant condition and condition-by-ASI perimeter interaction (int) effects for synapse subsets defined by d_ag_ (BH-adjusted *p < 0.05, **p < 0.01; see Table S2 for statistical details). Scale cube (edge length = 100 nm) in A applies to A-B. Cluster legend in E applies to E-F.

### Decrease in astroglial surround at ASI perimeter of large synapses during LTP

To assess the impact of LTP on astroglial apposition, the length of the ASI perimeter surrounded by astroglia was measured as depicted in Figures 2G and 5A. On average, astroglia surrounded only 50% of the ASI perimeter and did not fully envelop any MML synapses (Fig. S1C). The mean length of astroglial surround was positively correlated with total ASI perimeter for both small and large spine clusters (Fig. 5B, C). By 30 minutes and 2 hours after LTP induction, the extent of astroglia surrounding the ASI perimeter was reduced for cluster 2 synapses across all distance thresholds from 10-120 nm, but not for cluster 1 synapses (Fig. 5B, C). Therefore, these results indicate that as spines and their ASI perimeters enlarged during LTP, a smaller proportion of their interface remained in contact with surrounding astroglial processes.

**Figure 5.**
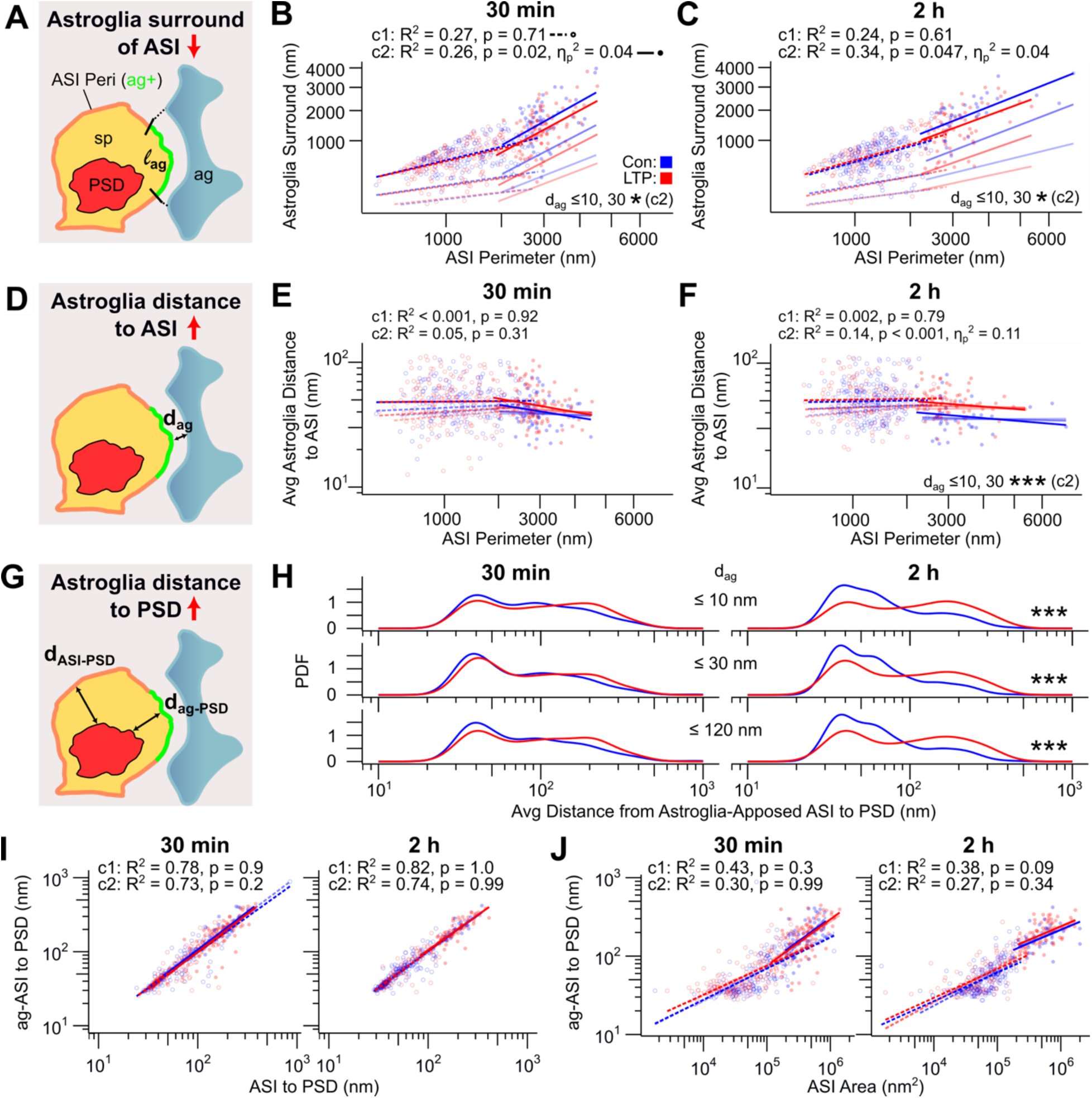
By 2 hours during LTP, PAP coverage length is reduced, and the minimum PAP distance to the ASI perimeter is increased selectively at large synapses. Meanwhile, astroglial access to the PSD decreases across synapse sizes. **(A)** Schematic representation of the ASI perimeter (orange) and the segment of its length (l_ag_, green) surrounded by perisynaptic astroglia (ag, light blue) within a given distance threshold (d_ag_, dashed black lines), with key synaptic structures (spine, yellow; PSD, red) also indicated. **(B)** Regression plot of l_ag_ (square-root y-axis) versus ASI perimeter (square-root x-axis) for c1 and c2 synapses with d_ag_ ≤ 120 nm at 30 min and **(C)** 2 h during LTP. In B-C, l_ag_ was measured using d_ag_ thresholds of ≤ 10 nm (lowest opacity lines), 30 nm (medium opacity), and 120 nm (highest opacity). Raw data, R² values, and Benjamini-Hochberg (BH)-adjusted p-values (condition effect) for d_ag_ ≤ 120 nm are displayed. All subsequent regression plots in (E-F, I-J) were performed separately for c1 and c2 synapses with d_ag_ ≤ 10, 30, and 120 nm, using the same marker and opacity scheme as in B-C, with statistical results displayed similarly. **(D)** Illustration of the average distance (black d_ag_ arrow) from the ASI perimeter to the nearest astroglial process, using the same schematic elements as in A. **(E)** Regression plot of average d_ag_ (log-scale y-axis) versus ASI perimeter (square-root x-axis) at 30 min and (**F**) 2 h. **(G)** Illustration of the average distance from astroglia-apposed ASI perimeter (within 120 nm) to the nearest PSD edge (black d_ag-PSD_ arrow), using the same schematic elements as in A. **(H)** Kernel density plots of average d_ag-PSD_ (log-scale y-axis) for synapses grouped by d_ag_ at 30 min (left) and 2 h (right). **(I)** Regression plot of average d_ag-PSD_ (log-scale y-axis) versus average distance from all ASI perimeter vertices to the nearest PSD edge (d_ASI-PSD_, log-scale x-axis), and (**J**) ASI area (log-scale x-axis), at 30 min (left) and 2 h (right). In all plots (B-C, E-F, H-J), data are shown for control (blue) and LTP (red) synapses (see Figure 3G for synapse counts). Asterisks indicate significant condition effects for synapse subsets defined by d_ag_ (BH-adjusted *p < 0.05, **p < 0.01, ***p < 0.001; see Table S3 for statistical details). Cluster legend in B applies to B-C, E-F, I-J.

### Increase in minimum astroglial distance to the ASI perimeter at large synapses during LTP

To evaluate changes in astroglial proximity during LTP, the minimum distance from the ASI perimeter to the nearest astroglial process was measured for all vertices with d_ag_ ≤ 120 nm (see Figs. 2J, 5D; Methods). Astroglia processes were 46-48 nm from the ASI perimeter, and this minimum distance did not differ between control and LTP when all MML synapses were analyzed together (Fig. S1D). For cluster 2 synapses, however, the mean minimum astroglial distance remained unchanged at 30 minutes (Fig. 5E) but was significantly increased at 2 hours during LTP across all distance thresholds (10-120 nm) (Fig. 5F). This suggests that in addition to reducing their length of surround, astroglial processes gradually withdrew from the ASI perimeter of large synapses during LTP.

### Decrease in astroglial access to the PSD during LTP

To examine whether LTP affects astroglial access to the PSD, we next measured the average distance from the PSD to the region of the ASI perimeter contacted by perisynaptic astroglia within 120 nm (Fig. 5G; Methods). This PSD-to-astroglia distance was significantly increased by 2 hours during LTP (Fig. 5H) and showed positive correlations with both the PSD-to-overall ASI perimeter distance (Fig. 5I) and ASI area (Fig. 5J). Notably, the PSD-to-astroglia distance scaled nearly one-to-one with the PSD-to-ASI perimeter distance under both control and LTP conditions, suggesting no preferential displacement toward or away from astroglial apposition (Fig. 5I). After controlling for ASI area, the effect of LTP on the PSD’s distance to astroglial apposition was no longer significant (Fig. 5J). Together, these results indicate that, in addition to increasing PSD offset, spine enlargement during LTP also reduced astroglial access to the PSD.

### PAP apposition at OML synapses

By applying the same strategies in the OML as used to analyze MML synapses, we found that more than 85% of synapses had astroglia within 120 nm of some region of the ASI perimeter, with around 60% having astroglia within 10 nm (Fig. 6D). Hence, like in the MML, the majority of OML synapses showed close astroglial apposition under control and cLTD conditions, and synapses with astroglial apposition within 10, 30, or 120 nm were included in all subsequent analyses.

**Figure 6.**
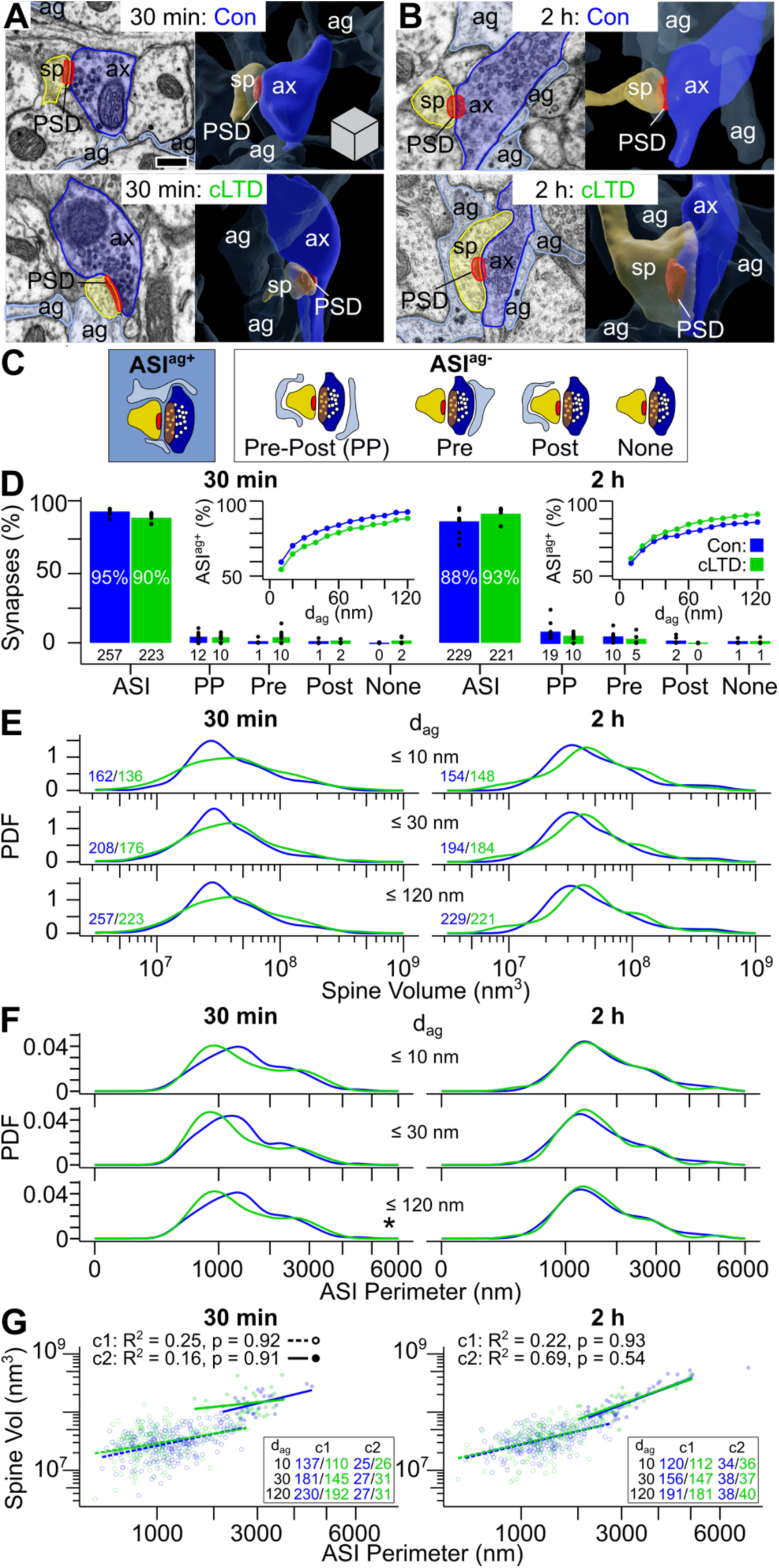
Nearly all OML spines have some perisynaptic astroglia and undergo a transient decrease in ASI perimeter length during cLTD relative to control stimulation. **(A)** EM image (left) and 3D reconstruction (right) of the spine (sp, yellow), postsynaptic density (PSD, red), presynaptic axon (ax, dark blue), and perisynaptic astroglia (ag, light blue) for an OML synapse during control (top) and cLTD (bottom) at 30 min and **(B)** 2 h after cLTD induction. **(C)** Synapse categories based on astroglial presence (ASI^ag+^) or absence (ASI^ag-^) within 120 nm of the ASI perimeter and specific astroglia location. The spine, axon, PSD, perisynaptic astroglia, presynaptic vesicles (small white circles), and ASI (brown oval) are represented using same color scheme as A-B. **(D)** Bar graph of overall synapse percentage in each synapse category. Numbers below bars indicate synapse count. Dots represent dendrite-specific synapse percentages. (Insets) Line plots show ASI^ag+^ synapse percentages based on each d_ag_ threshold tested. **(E)** Kernel density estimate plots show the probability density functions for spine volume (log-scale axis) and **(F)** ASI perimeter (square-root axis) for synapse subsets based on d_ag_ (*p < 0.01 that control and cLTD distributions differ). Synapse counts color-coded by condition are shown in bottom left corner in E and apply to E-F. **(G)** Regression plot of spine volume (log-scale y-axis) and ASI perimeter (square-root x-axis) for c1 or c2 synapses with d_ag_ ≤ 10 nm (lowest opacity), 30 nm (medium opacity), and 120 nm (highest opacity). Raw data, R² values, and Benjamini-Hochberg (BH)-adjusted p-values (condition effect) for d_ag_ ≤ 120 nm are displayed. Synapse counts color-coded by condition are shown in bottom right corner (relatively few synapses in cluster 2 had d_ag_ ≤ 10 nm). In D-G, data are from control (blue) and cLTD (green) OML synapses at 30 min (left) and 2 h (right) during cLTD. Asterisks indicate significant condition effects for synapse subsets defined by d_ag_ (BH-adjusted *p < 0.05; see Table S4 for statistical details). Scale bar and cube (edge length = 250 nm) in A apply to A-B.

### Transient increase in relative frequency of synapses with shorter ASI perimeters during cLTD

Both spine volume and the length of the ASI perimeter were greater at synapses with astroglia present along some portions of the ASI perimeter under control and cLTD conditions (Fig. S2A, B). Although cLTD did not significantly affect spine volume (Fig. 6E; S2A), the distribution of ASI perimeter lengths shifted leftward at 30 minutes—but not at 2 hours—reaching statistical significance when astroglia located within 120 nm were included (Fig. 6F). These results suggest that while spine volume remained stable during cLTD, there was a transient increase in the relative prevalence of synapses with shorter ASI perimeters.

### K-means clustering based on spine volume and ASI perimeter in the OML

Similar to the MML, OML ASI^ag+^ synapses were grouped into two clusters based on spine volume and ASI perimeter (see Methods). Cluster 2 synapses, comprising 15% of OML ASI^ag+^ synapses, had larger spine volumes and ASI perimeters compared to cluster 1 synapses (Fig. 6G). The relative proportion of cluster 2 synapses remained stable between control and cLTD conditions (Table S4). Spine volume and ASI perimeter were also positively correlated across synapse clusters, with no observed effect of cLTD on this relationship (Fig. 6G). Therefore, once again, in all subsequent regression analyses of OML synapses, these unique clusters of small and large spines were analyzed separately.

### Transient ASI perimeter shrinkage temporarily decreases PSD offset from the ASI center during cLTD

Although cLTD had minimal effect on PSD area (Fig. 7C, S5A), PSD offset from the ASI center—47-54 nm under control conditions—was significantly reduced at 30 minutes during cLTD (Fig. 7D). As in the MML, both PSD area and offset were positively correlated with ASI area (Fig. 7E, F). Moreover, after controlling for spine size, the effect of cLTD on PSD displacement was no longer evident (Fig. 7F). Neither PSD area nor PSD offset showed consistent relationships with the length of astroglial surround (Fig. S5C, E), the astroglial distance to the ASI perimeter (Fig. S5D, F), or with each other (Fig. S5G). Taken together, these findings suggest that during cLTD, the transient decrease in ASI perimeter coincided with a higher prevalence of synapses with more centrally located PSDs within the ASI, independent of the extent of astroglial apposition.

**Figure 7.**
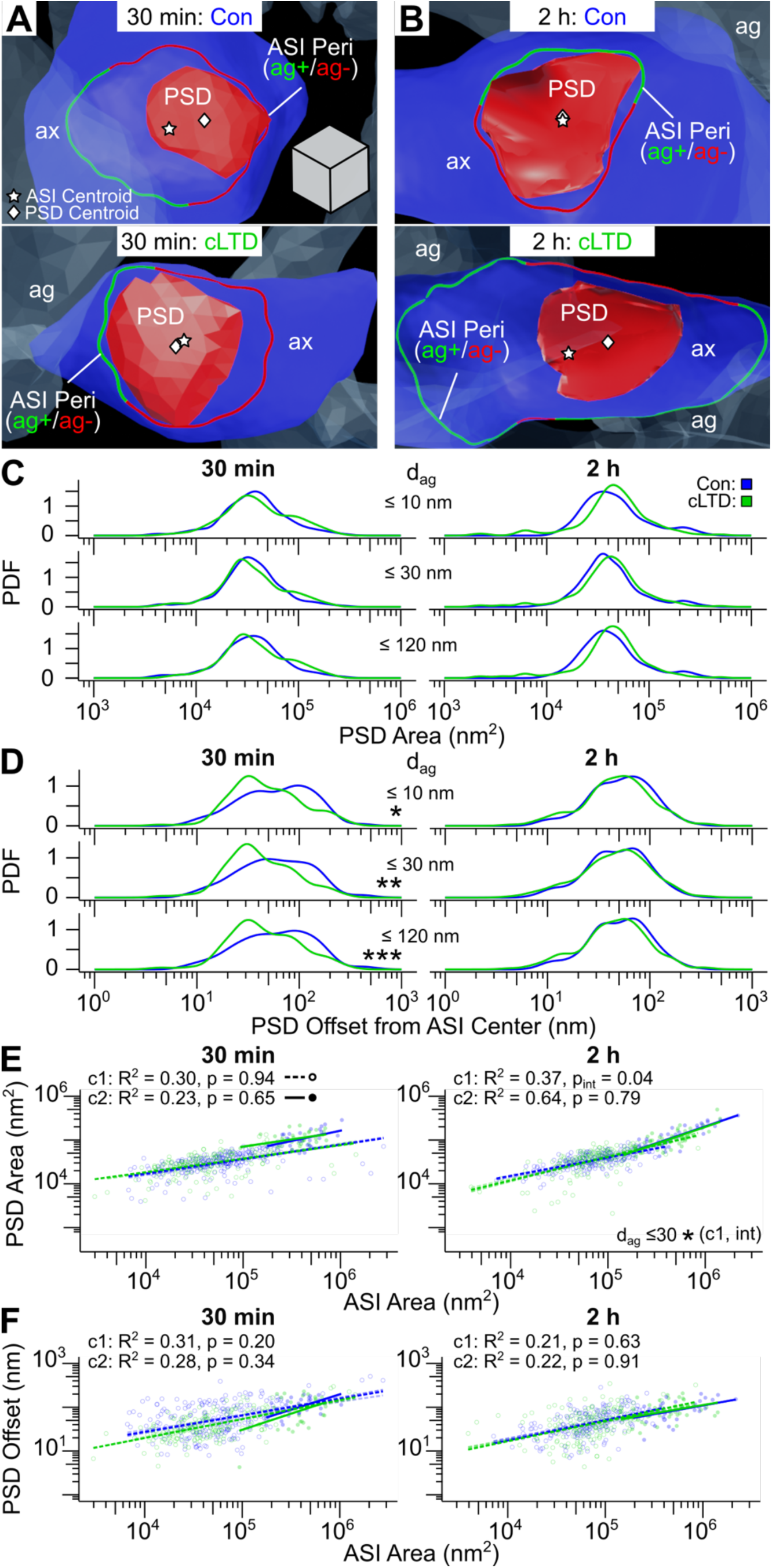
The PSD area does not change, but the PSD offset from the ASI center transiently decreases during cLTD. **(A)** 3D reconstruction of the postsynaptic density (PSD, red) projected onto the axon mesh (ax, dark blue), perisynaptic astroglia (ag, light blue), ASI perimeter color coded by presence (ag+, green) or absence (ag-, red) of astroglia within 120 nm, ASI centroid (white star), and PSD centroid (white diamond) for an OML synapse during control (top) and cLTD (bottom) at 30 min and **(B)** 2 h following cLTD induction. **(C)** Kernel density estimate (KDE) plots show the probability density function (PDF) for PSD area (log-scale axis) and **(D)** PSD offset from the ASI center (log-scale axis) for synapses subset based on d_ag_. **(E)** Regression plot of PSD area (log-scale y-axis) and **(F)** PSD offset from the ASI center (log-scale y-axis) versus ASI area (log-scale x-axis) for c1 or c2 synapses with d_ag_ ≤ 10 nm (lowest opacity lines), 30 nm (medium opacity lines), and 120 nm (highest opacity lines). Raw data, R² values, and Benjamini-Hochberg (BH)-adjusted p-values (condition effect) for d_ag_ ≤ 120 nm are displayed. In C-F, data are from control (blue) and cLTD (green) OML synapses at 30 min (left) and 2 h (right) during cLTD (see Figure 6G for synapse counts). Asterisks indicate significant condition and condition-by-ASI perimeter interaction (int) effects for synapse subsets defined by d_ag_ (BH-adjusted *p < 0.05, **p < 0.01, ***p < 0.001; see Table S5 for statistical details). Scale cube (edge length = 100 nm) in A applies to A-B. Cluster legend in E applies to E-F.

### Stable amount of PAP surrounding the ASI perimeter during cLTD

Evaluation of OML ASI^ag+^ synapses revealed that most had about 50% of the length of their ASI perimeters surrounded by astroglial processes (Fig. S2C). Additionally, the length of the apposition correlated positively with ASI perimeter across synapse clusters (Fig. 8B, C). At 2 hours during cLTD, significant interactions between condition and ASI perimeter influenced the mean length of astroglial surround for cluster 2 synapses across all distance thresholds from 10-120 nm (Fig. 8C). However, no cLTD-related effects were found at 30 minutes, nor for cluster 1 synapses at either time point. These results indicate that the extent of astroglial apposition at the ASI perimeter remained largely unchanged during cLTD.

**Figure 8.**
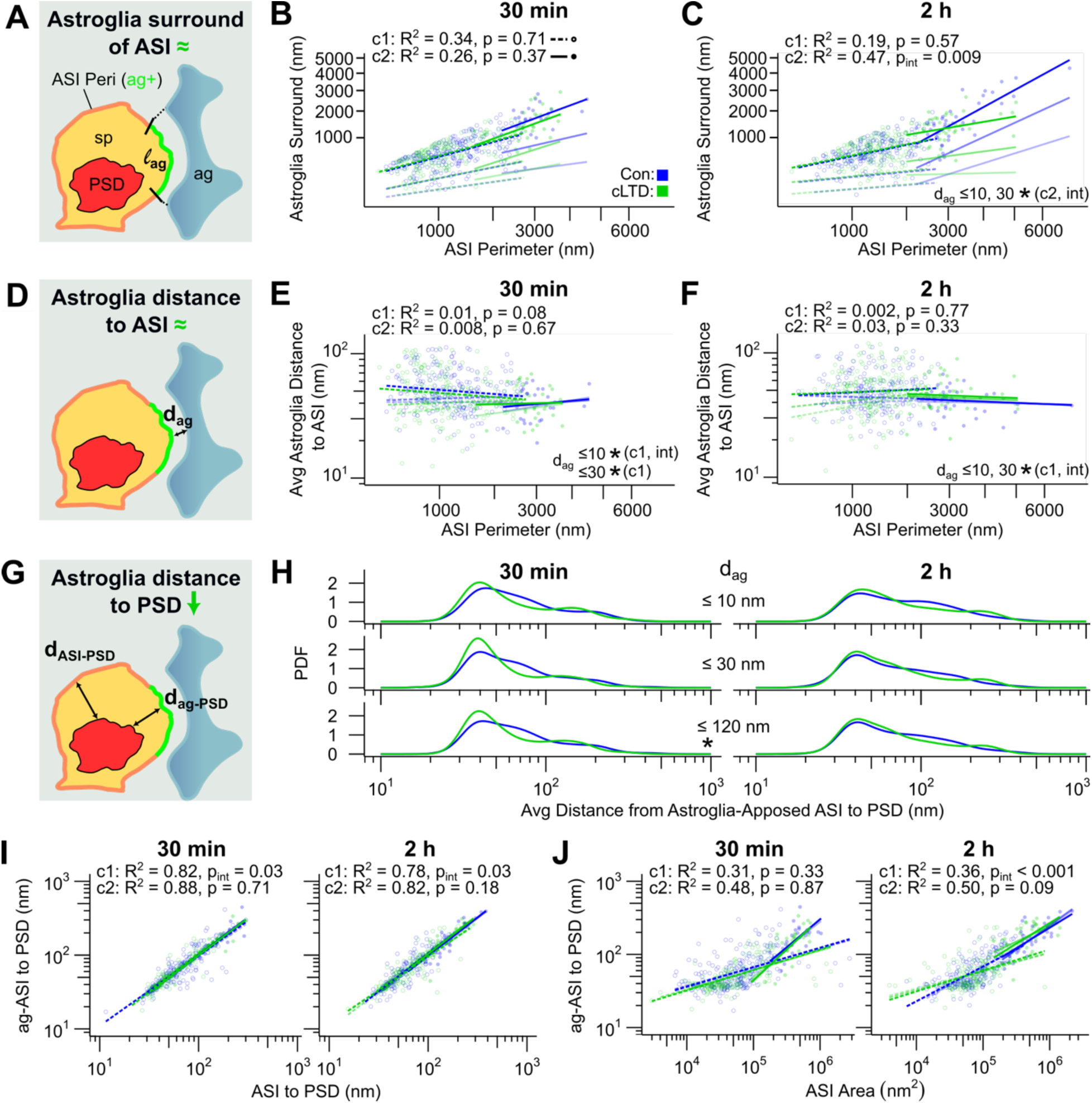
During cLTD, there is a minimal change in PAP coverage length and minimum distance to the ASI perimeter, but a transient increase in astroglial access to the PSD. **(A)** Schematic representation of the ASI perimeter (orange) and the segment of its length (l_ag_, green) surrounded by perisynaptic astroglia (light blue) within a given distance threshold (d_ag_, dashed black lines), with key synaptic structures (spine, yellow; PSD, red) also indicated. **(B)** Regression plot of l_ag_ (square-root y-axis) versus ASI perimeter (square-root x-axis) for c1 and c2 synapses with d_ag_ ≤ 120 nm at 30 min and **(C)** 2 h during cLTD. In B-C, l_ag_ was measured using d_ag_ thresholds of ≤ 10 nm (lowest opacity lines), 30 nm (medium opacity), and 120 nm (highest opacity). Raw data, R² values, and Benjamini-Hochberg (BH)-adjusted p-values (condition effect) for d_ag_ ≤ 120 nm are displayed. All subsequent regression plots in (E-F, I-J) were performed separately for c1 and c2 synapses with d_ag_ ≤ 10, 30, and 120 nm, using the same marker and opacity scheme as in B-C, with statistical results displayed similarly. **(D)** Illustration of the average distance (black d_ag_ arrow) from the ASI perimeter to the nearest astroglial process, using the same schematic elements as in A. **(E)** Regression plot of average d_ag_ (log-scale y-axis) versus ASI perimeter (square-root x-axis) at 30 min and **(F)** 2 h. **(G)** Illustration of the average distance from astroglia-apposed ASI perimeter (within 120 nm) to the nearest PSD edge (black d_ag-PSD_ arrow), using the same schematic elements as in A. **(H)** Kernel density plots of average d_ag-PSD_ (log-scale y-axis) for synapses grouped by d_ag_ at 30 min (left) and 2 h (right). **(I)** Regression plot of average d_ag-PSD_ (log-scale y-axis) versus average distance from all ASI perimeter vertices to the nearest PSD edge (d_ASI-PSD_, log-scale x-axis), and (**J**) ASI area (log-scale x-axis), at 30 min (left) and 2 h (right). In all plots (B-C, E-F, H-J), data are shown for control (blue) and cLTD (green) synapses (see Figure 6G for synapse counts). Asterisks indicate significant condition and condition-by-ASI perimeter interaction (int) effects for synapse subsets defined by d_ag_ (BH-adjusted *p < 0.05; see Table S6 for statistical details). Cluster legend in B applies to B-C, E-F, I-J.

### Transient decrease in minimum PAP distance to the ASI perimeter during cLTD

On average, astroglial processes apposed OML synapses at a minimum distance of 49 nm. No significant differences in the mean PAP apposition distance were observed between control and cLTD conditions when OML synapses were analyzed collectively (Fig. S2D), and this minimal distance did not correlate with ASI perimeter across synapse clusters (Fig. 8E, F). Nonetheless, for cluster 1 synapses with astroglia within 30 nm of the ASI, the mean minimum PAP distance to the ASI was significantly decreased at 30 minutes during cLTD (Fig. 8E). Significant interactions between condition and ASI perimeter were also observed for cluster 1 synapses with astroglia within 10 nm at both 30 minutes and 2 hours, and within 30 nm at 30 minutes (Fig. 8E, F). Therefore, the transient spine morphology changes associated with cLTD were accompanied by modest but specific increases in astroglial proximity, particularly at distances close to the ASI.

### Transient increase in astroglial access to the PSD during cLTD

At 30 minutes during cLTD, the average distance from the PSD to the region of the ASI perimeter apposed by astroglia was transiently reduced (Fig. 8H). As observed in the MML, this distance correlated positively with the PSD’s offset from the overall ASI perimeter. Additionally, this relationship had a slope close to one under both control and cLTD conditions—indicating that astroglial positioning does not influence the direction of PSD displacement (Fig. 8I). After controlling for ASI area, the effect of cLTD on the PSD-to-astroglia distance was no longer significant (Fig. 8J). These findings suggest that spine morphology changes associated with cLTD temporarily enhanced astroglial access to the PSD.

### Comparable amount of perisynaptic astroglia at MML and OML control synapses

To test whether inherent baseline differences between control MML and OML layers were responsible for the observed differences between effects of LTP and cLTD at dentate gyrus synapses, we repeated all the analyses by comparing MML and OML ASI^ag+^ synapses under control conditions, with the 30 minute and 2 hour time points pooled.

The proportion of synapses with astroglial apposition at the ASI did not differ between the dentate gyrus layers (Fig. 9A). However, the MML had a higher relative percentage of cluster 2 synapses compared to the OML (Fig. 9B). ASI perimeters were comparable between layers (Fig. 9C), whereas PSD areas were larger in the MML than in the OML (Fig. 9D). An interaction between layer and ASI perimeter influenced the mean minimum PAP distance to the ASI for cluster 1 synapses (Fig. 9E). In contrast, this distance for cluster 2 synapses (Fig. 9E), as well as the average fraction of astroglial surround for either cluster (Fig. 9F), did not significantly differ between layers. No significant differences were observed for the distributions of PSD offset (Fig. 9G) and astroglial distance to the PSD (Fig. 9H).

**Figure 9.**
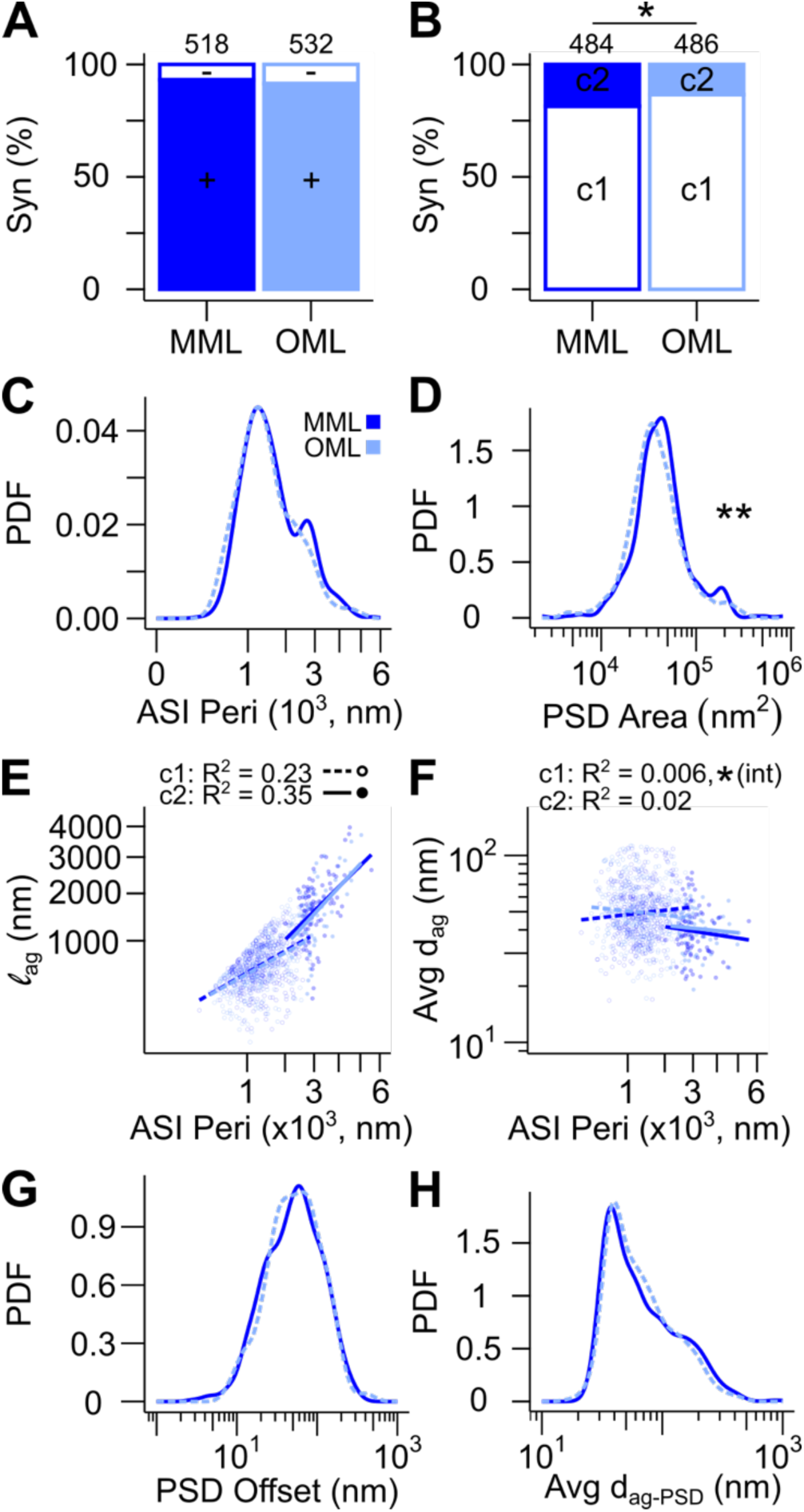
Postsynaptic ultrastructure and astroglial coverage of synapses in the MML are mostly comparable to synapses in the OML under control conditions. **(A)** Stacked bar graphs showing the relative percentage of ASI^ag-^ (-, white, d_ag_ > 120 nm) versus ASI^ag+^ (+, sky blue, d_ag_ ≤ 120 nm) synapses and **(B)** c1 (open) versus c2 (filled) synapses in each layer. Numbers above bars represent total synapse counts. **(C)** Kernel density estimate plots (KDE) showing the probability density function (PDF) for ASI perimeter (square-root axis) and **(D)** PSD area (log-scale axis). **(E)** Regression plots of the length of the ASI perimeter surrounded by astroglia within 120 nm (l_ag_, square-root y-axis) and **(F)** the average astroglia distance to the ASI perimeter (avg d_ag_, log-scale y-axis) versus ASI perimeter (square-root x-axis) for c1 or c2 synapses. Raw data and R² values are displayed. **(G)** KDE showing the PDF for PSD offset from the ASI center (log-scale axis) and **(H)** average distance between the ASI perimeter segment with astroglial apposition within 120 nm to the nearest PSD edge (d_ag-PSD_, log scale axis). In A-H, data are from control MML (dark blue) and OML (light blue) synapses (time points pooled), with only synapses with d_ag_ ≤ 120 nm included in analyses plotted in B-H (see Figures 3G and 6G for synapse counts). Asterisks indicate significant layer and layer-by-ASI perimeter interaction (int) effects (*p < 0.05, **p < 0.01; see Table S7 for statistical details). Cluster legend in E applies to E-F.

Finally, the ASI perimeter (Fig. 10A) and PSD area (Fig. 10B) distributions differed significantly during LTP versus cLTD. However, condition and layer did not interact to influence either the mean length of astroglial surround (Fig. 10C) or PAP distance to the ASI perimeter (Fig. 10D). LTP and cLTD conditions also showed significant differences in PSD offset (Fig. 10E) and average PAP distance to the PSD (Fig. 10F). Together, these results suggest that, while some baseline differences exist between molecular layers, they are unlikely to explain the different LTP- and cLTD-related changes observed in this study.

**Figure 10.**
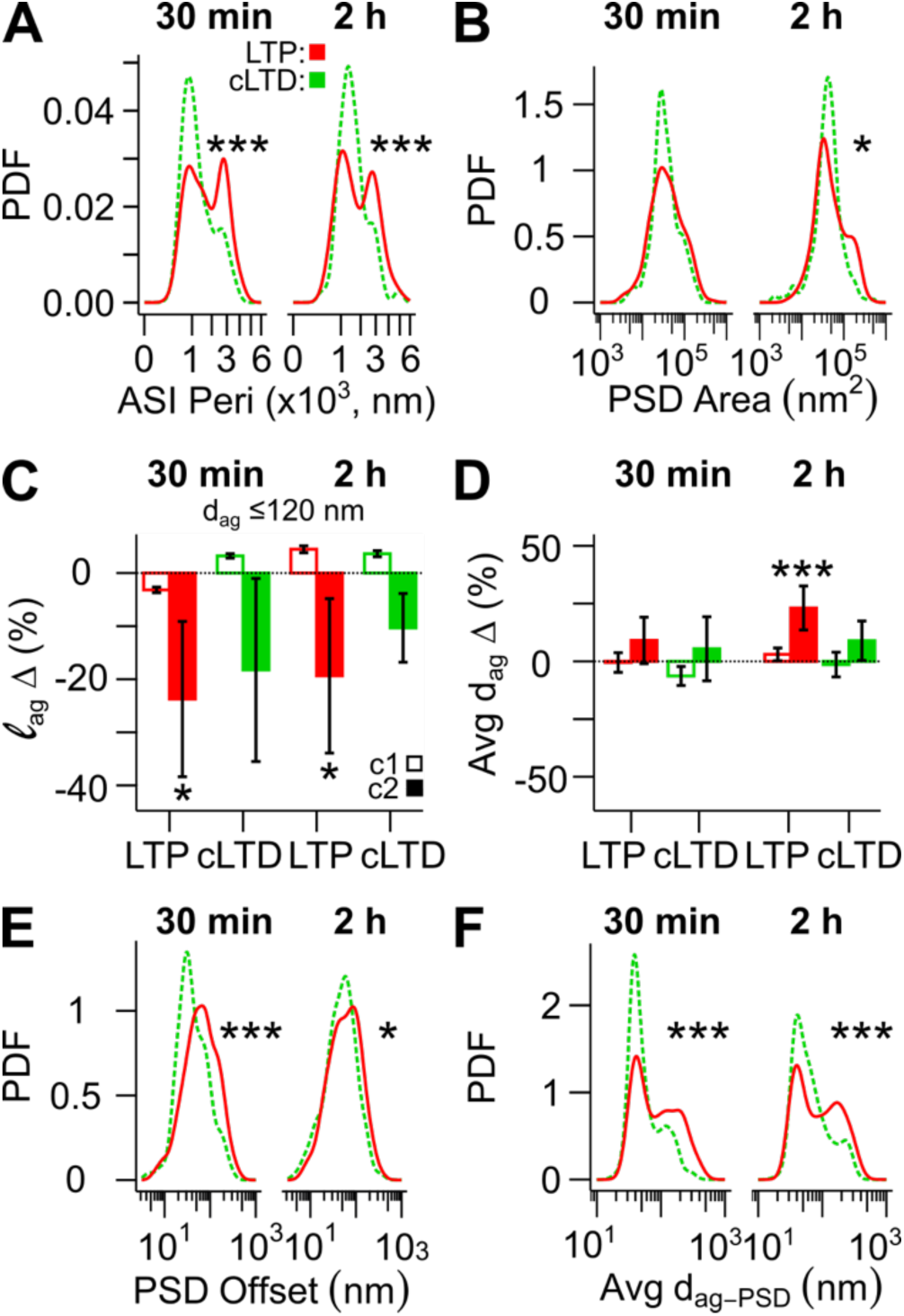
Comparison of relative changes in tripartite synapse ultrastructure associated with LTP and cLTD in the dentate gyrus. **(A)** Kernel density estimate (KDE) plots showing the probability density function (PDF) for ASI perimeter (square-root axis) and **(B)** PSD area (log-scale axis). **(C)** Bar graphs show the percentage change in the length of the ASI perimeter surrounded by astroglia within 120 nm (l_ag_) and **(D)** the average distance between astroglia and the ASI perimeter (d_ag_) during LTP or cLTD relative to control. In C-D, percent changes were calculated after controlling for ASI perimeter, and separately for c1 (open bars) and c2 (filled bars) synapses. Stars above bars indicate significance levels for condition effects based on layer-specific analyses and error bars indicate standard errors. **(E)** KDE plots show the PDF for PSD offset (log-scale axis) and **(F)** the average distance between astroglia and the nearest PSD edge (d_ag-PSD_, log-scale axis). In A-F, data are from MML synapses during LTP (red) and OML synapses during cLTD (green) with d_ag_ ≤ 120 nm, at 30 min (left) and 2 h (right) after LTP or cLTD induction (see Figures 3E, 3G, 6E, and 6G for synapse counts). *p < 0.05, ***p < 0.001 (see Table S8 for statistical details).

## Discussion

We found that a large majority of dendritic spines in the dentate gyrus have perisynaptic astroglia at the perimeter of their ASI. The relative frequencies of these tripartite synapses were not altered during LTP or cLTD; however, proximity of perisynaptic astroglial processes to the synapses was affected by the synaptic plasticity (Figs. 11, 12). The range of spine volumes was expanded during LTP, resulting in a net increase in large spines and PSD areas, the latter of which we noted for the first time to be offset from the center of the ASI (Fig. 11A). Since astroglial proximity to the ASI perimeter of the enlarged synapses was reduced, this decreased access of astroglia to the PSD (Fig. 12A). Meanwhile, cLTD produced only subtle decreases in spine volume, PSD area, and the proximity of astroglia to the ASI perimeter (Figs. 11B, 12B). During cLTD, there was a transient initial decrease in the length of the ASI perimeter, possibly reflecting a reduced complexity in spine shape, which would enhance astroglial access to the PSD (Fig. 12B).

**Figure 11.**
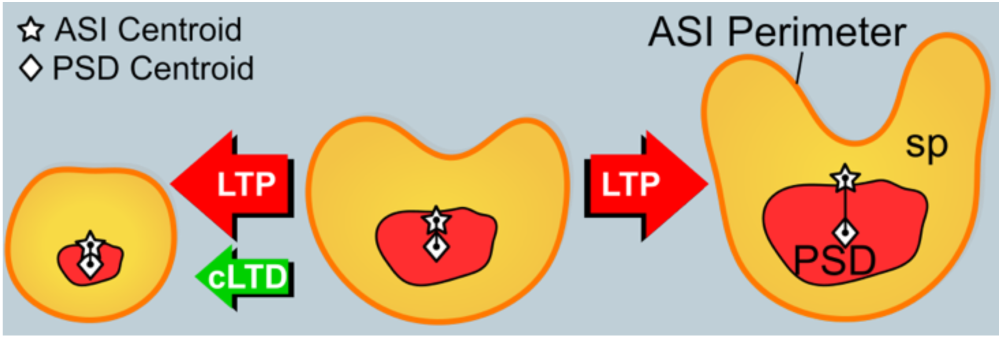
Summary of main effects on spines and PSDs. Under all conditions and spine sizes, the PSD was offset from the center of the ASI perimeter. The distribution of spine volumes and PSD areas expanded, increasing the frequency of both smaller and larger synapses at both 30 min and 2 h after the induction of LTP (red arrows). The ASI perimeter and PSD offset distributions shifted unidirectionally toward larger values at 2 h during LTP. In contrast, a shift towards spines with smaller ASI perimeters and less PSD offset from the ASI center and perimeter occurred transiently at 30 min during cLTD (green arrow).

**Figure 12.**
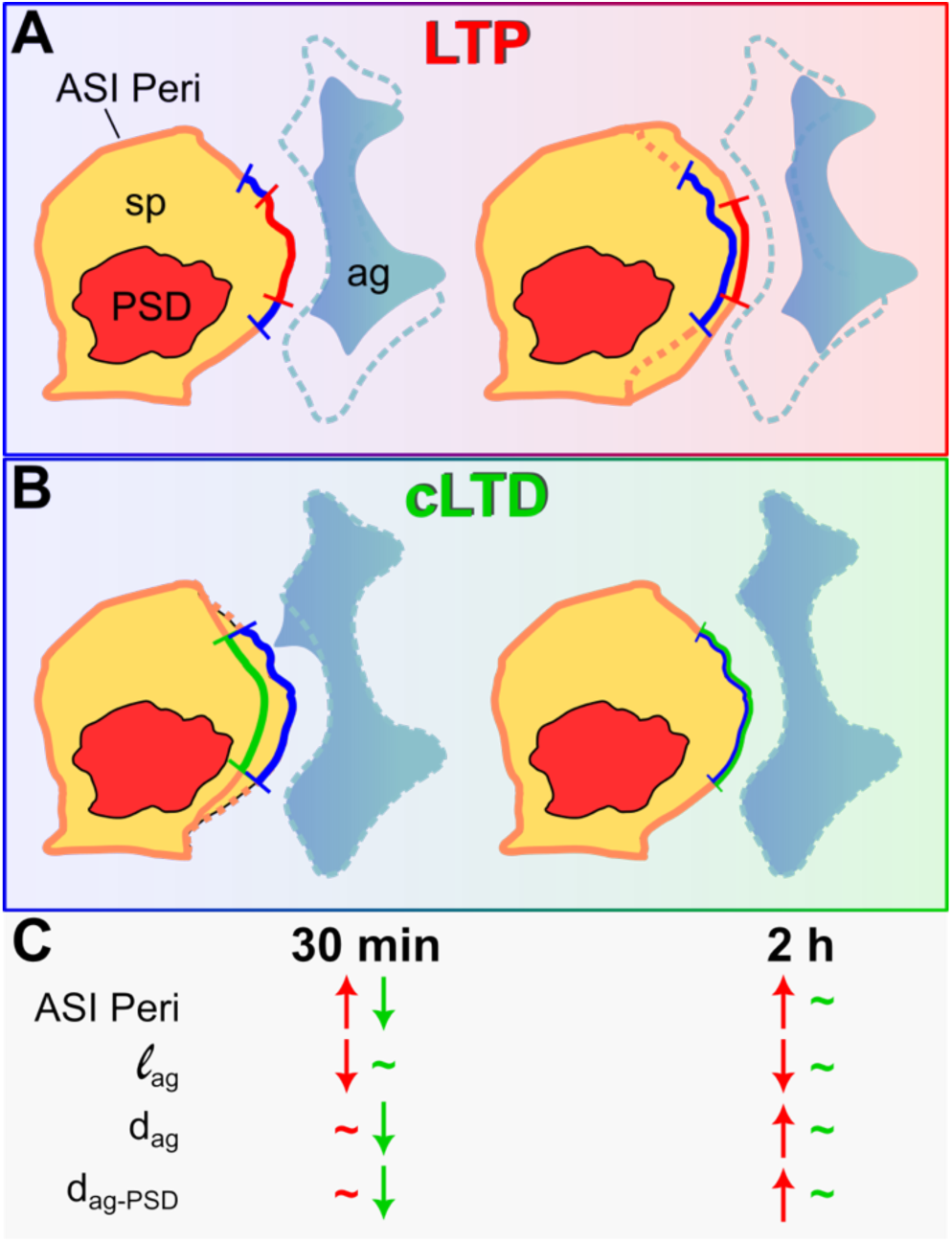
Summary of main effects between perisynaptic astroglia and synapses. **A)** The length of astroglia (l_ag_) apposed to the ASI perimeter was reduced by 30 min and further reduced by 2 h during LTP as spines and the ASI enlarged. The distances to the astroglia (d_ag_ and d_ag-PSD_) increased, resulting in reduced astroglial access to the PSD. **B)** In contrast, the length of the ASI perimeter and distances to the astroglia (d_ag_ and d_ag-PSD_) were decreased somewhat at small synapses by 30 min during cLTD, providing a transient increase in astroglial access to the PSD. **C)** Summary of effects at 30 min and 2 hr for LTP (red) and cLTD (green).

We developed a novel 3D approach to establish reliable criteria to investigate how astroglia relate to synapses on dendritic spines. Application of this or similar approaches will be needed to compare astroglial coverage of synapses across brain regions. For example, we showed that synapses can vary dramatically depending on the criteria used for astroglial apposition in the dentate gyrus, from 60% of synapses having astroglia located within 10 nm of the ASI perimeter to 85% having astroglia located within 120 nm. This latter frequency surpasses previous estimates based on single-section analyses that reported <40% of dentate gyrus synapses having astroglia at their ASI perimeters (Wenzel et al., 1991). Single-section sampling would fail to capture that most synapses are not surrounded by glia, inadvertently misclassifying those with partial astroglial apposition as lacking glia coverage altogether.

To define a functionally relevant threshold for astroglial apposition, we restricted our analyses to synapses with astroglia located no more than 120 nm from the ASI perimeter. Mathematical models indicate that when astroglia appose synapses within 150 nm, the increased glutamate uptake diminishes AMPA receptor currents by 50% (Pannasch et al., 2014). Therefore, this 120 nm cut-off allowed us to focus on synapses with reasonable physiological access to their surrounding PAP.

Our findings in the dentate gyrus contrast with other 3D EM studies in hippocampal area CA1, where approximately 62% of synapses were reported to have perisynaptic astroglia (Ventura and Harris, 1999; Witcher et al., 2007). Less strict criteria for apposition may explain why other reports suggest >80% of CA1 synapses have perisynaptic astroglia, which could include astroglia that are separated from the ASI by intervening structures (Chai et al., 2017; Gavrilov et al., 2018). Additionally, the fraction of neuropil occupied by astroglia can be higher near smaller synapses (Medvedev et al., 2014; Gavrilov et al., 2018). Thus, differences in spine and synapse size might account for the disparity in astroglial coverage between smaller synapses in the dentate gyrus and those spanning a larger size range in area CA1 (Harris and Stevens, 1989).

In the dentate gyrus, both LTP and cLTD were associated with significant morphological changes at tripartite synapses. LTP expanded the range of spine and PSD sizes, consistent with prior findings (Bromer et al., 2018). Meanwhile, cLTD in the OML did not lead to the decrease in spine size that was previously linked to LTD in the CA1 region (Okamoto et al., 2004; Zhou et al., 2004). Instead, we observed an increase in spines with shorter ASI perimeters but not necessarily smaller volumes, suggesting a temporary shift towards less complex spine morphologies during cLTD in the dentate gyrus.

Our findings suggest that astroglia maintain a delicate balance between too little and too much synapse coverage to establish a permissive environment for synaptic plasticity. LTP and cLTD-associated changes occurred primarily at synapses with astroglial apposition. Furthermore, consistent with previous findings (Witcher et al., 2007; Bellesi et al., 2015), mean spine and synapse size were larger for synapses with astroglial coverage. In contrast, no dentate gyrus synapse exhibited 100% PAP coverage. Moreover, cLTD did not impact astroglia-ASI apposition, but LTP led to selective astroglial withdrawal from the perimeter of large spine synapses. Rats reared in a complex environment show increased astroglia-synapse contact (Jones and Greenough, 1996). At the same time, astroglial contact at synapses has been linked to inhibited synapse growth during memory consolidation (Ostroff et al., 2014), whereas astroglial retraction from synapses is associated with enhanced fear memory (Badia-Soteras et al., 2023). Therefore, non-uniform PAP coverage at the ASI perimeter of dentate gyrus synapses may differentially affect synaptic efficiency.

In area CA1, astroglial processes withdraw from synapses following LTP induction, leading to an increase in NMDA receptor-dependent extrasynaptic communication, with no selectivity based on spine size (Henneberger et al., 2020). Astroglia organize synapses into astroglia-defined synaptic clusters (Salmon et al., 2023), and model predictions suggest that processes positioned farther away from synapses are better able to maintain calcium microdomains (Toman et al., 2023). Thin spines are more often transient, while larger mushroom-shaped spines are generally considered to be more permanent, leading to their proposed designations as “learning” and “memory” spines, respectively (Bourne and Harris, 2007). Astroglial withdrawal reduces glutamate uptake (Gavrilov et al., 2018; Henneberger et al., 2020), likely enhancing synaptic transmission (Pannasch et al., 2014). However, it also diminishes the synaptic availability of astroglia-derived D-serine and glycine (Le Bail et al., 2015), potentially impairing further LTP at large spines (Perez-Alvarez et al., 2014), as astrocytic glycine is particularly important for LTP in the dentate MML (Sateesh and Abraham, 2025). Therefore, the LTP-associated retraction of astroglial processes specifically from large dentate gyrus synapses may represent a mechanism tailored to this region’s role in pattern separation, potentially enhancing the network contribution of stable “memory” spines (Aimone et al., 2011; Hassanpoor and Saidi, 2020).

Adding complexity to our understanding of astroglial-synapse coverage, we also found that the PSD center is, on average, located ∼50 nm from the ASI center. LTP was associated with increases in both the degree of PSD offset and the distance between surrounding PAP and the nearest PSD edge. In contrast, cLTD induction resulted in decreases in both these measures. The extent of PSD offset showed a stronger positive correlation with ASI area than PSD area. The PSD offset direction did not occur preferentially towards or away from astroglia-ASI apposition. Hence, alterations in spine rather than PSD size likely contributed to the altered PSD offset during LTP and cLTD. These changes, alongside shifts in PAP proximity, may have shaped astroglial access to the PSD. In addition, these observations highlight that the extent of astroglial coverage at the ASI perimeter alone provides an incomplete understanding of astroglia-synapse dynamics.

Although dentate gyrus astroglia exhibit both functional and morphological diversity (Kosaka and Hama, 1986; Bushong et al., 2003; Karpf et al., 2022; Viana et al., 2023), we observed only subtle baseline differences between synapses in the middle and outer molecular layers. The relative proportion of synapses with versus without astroglia apposition at the ASI did not differ significantly between these regions. Similarly, the extent of astroglial surround of and average astroglial proximity to the ASI perimeter were comparable between the MML and OML. Interestingly, the mean PSD area was significantly greater in the MML compared to the OML, even under control conditions. Astroglia form large, complex networks connected via gap junctions (Giaume et al., 2010; Chever et al., 2016; Mederos et al., 2018; Aten et al., 2022; Karpf et al., 2022). Therefore, selective astroglia coupling may partially account for this uniformity in astroglial-synapse coverage but not synapse size.

In this study, we showed that during LTP and cLTD in the dentate gyrus, astroglia processes in close proximity to synapses undergo morphological changes that both parallel and diverge from ultrastructural plasticity observed in other brain regions. Astroglia variability has been demonstrated across sex and developmental stages (Mouton et al., 2002; Conejo et al., 2003; Johnson et al., 2008; Rurak et al., 2022); therefore, the conclusions we draw here are limited to the dentate gyrus of the adult, male rodent brain. Additionally, given the subtle nature of the structural changes we observed, future studies would benefit from increased sample size, as well as the inclusion of both male and female animals. The efficacy of many astroglial synaptic functions is determined by the proximity of PAP to the synapse (Pannasch et al., 2014; Toman et al., 2023). Thus, future research should also explore how LTP and cLTD uniquely influence astroglial interactions with the PSD’s active versus nascent zones— subregions distinguished by presence or absence of apposing presynaptic vesicles, respectively. Finally, the dentate gyrus is one of the few regions where adult neurogenesis occurs (Aimone et al., 2011; Denoth-Lippuner and Jessberger, 2021). Hence, although the degree of PAP coverage is similar between adult-born and pre-existing synapses (Krzisch et al., 2015), exploring whether this consistency is maintained during synaptic plasticity could provide insights into the nuanced role of dentate gyrus astroglia in learning and memory.

## Author Contributions

AJN: conceptualization, methodology, investigation, formal analysis, writing – original draft, writing – review and editing. MK: investigation, writing – review and editing. PHP – investigation, writing – review and editing. WCA: contributed animal experiments, writing – review and editing. KMH: conceptualization, methodology, resources, funding acquisition, supervision of analyses and figures, writing – review and editing.

## Data Accessibility

The original EM images, PyReconstruct files (.jser format), and compiled quantitative measurements (in .csv format) used for statistical analysis in R will be made available on 3DEM.org via the Texas Data Repository (doi: https://doi.org/10.18738/T8/S8DT5E).

## Acknowledgements

We acknowledge that the tissue used in these studies were prepared as part of the thesis work reported in Bowden et al., 2012. We thank S.E. Mason-Parker for performing the *in vivo* electrophysiology and perfusions, J. Bowden for early work in the initiation of the experiments and early analyses, W. Yin for serial sectioning, tSEM imaging, and image alignment, J.M. Mendenhall for helping with tSEM imaging, B. Smith and L. Perry for initial serial sectioning and TEM imaging, L.F. Lindsey for helping with some of the original TrakEM2 image alignment, Dusten Hubbard for substantial tracing and editing of dendrites and spines, and J. Long for contributing to astroglia tracing. We also thank the team of students, including J. Ahn, S. Anand, P. Das, M. Ginjupalli, E. Haney, P. Mahableshwarkar, L. Robertson, and K. Rubenzer, who contributed to initial synapse and spine tracing efforts. The authors used ChatGPT (OpenAI) to assist with language editing and grammar correction during manuscript preparation. The authors acknowledge the Texas Advanced Computing Center (TACC, 3dem.org) at The University of Texas at Austin for providing computational resources that contributed to the results reported in this paper (URL: www.tacc.utexas.edu). Grant support: **NSF:** 2014862, 2219894; **NIH:** 1R56MH139176, 2R01MH095980

## Supplementary Material

**Table S1.** Summary of statistical output comparing spine volume, ASI perimeter, and astroglial presence at the ASI between control and LTP synapses in the MML. Statistical outcomes for comparisons visualized in. **Figure 3. Table columns (col):** (Col 1) Figure panel. (Col 2) Time following DBS onset: 30 min, 2 h, or Both. (Col 3) Synapse subset defined by d_ag_: ≤ 10 nm, ≤ 30 nm, ≤ 120 nm. (Col 4) Response variable (Var) and covariate (CoVar): PAP present (ag+) or absent (ag-) at ASI, spine volume (spVol), ASI perimeter (ASIperi), cluster 1 (c1), cluster 2 (c2). (Col 5) Synapse cluster: c1, c2, both. (Col 6) Statistical test: Chi-squared test (χ²), Kolmogorov-Smirnov test (KS), Linear mixed model (LMM). (Col 7) (Col 8) p-value for condition (C) effect. (Col 9) p-value for condition effect adjusted using the Benjamini-Hochberg (BH) procedure. (Col 10) Proportion of variance in the dependent variable explained by condition effect (partial eta², η_p_^2^). (Col 11) Coefficient of determination (R^2^). (Col 12) p-value for covariate effect adjusted using the BH procedure.

| Fig. | Time | d<sub>ag</sub><br>(nm) | Var,<br>CoVar | Cluster | Test | Statistic | p (C) | p<br>(C, adj) | $\eta_p^2$ | R<sup>2</sup> | p<br>(CoVar) |
| --- | --- | --- | --- | --- | --- | --- | --- | --- | --- | --- | --- |
| 3D | 30 min | 120 | ag- vs. ag+ | Both | $\chi^2$ | 0.099 | 0.75 | — | — | — | — |
| 3D | 2 h | 120 | ag- vs. ag+ | Both | $\chi^2$ | 2.5 | 0.11 | — | — | — | — |
| 3E | 30 min | 10 | SpVol | Both | KS | 0.2 | 0.0029 | 0.0029 | — | — | — |
| 3E | 30 min | 30 | SpVol | Both | KS | 0.21 | <0.001 | <0.001 | — | — | — |
| 3E | 30 min | 120 | SpVol | Both | KS | 0.19 | <0.001 | <0.001 | — | — | — |
| 3E | 2 h | 10 | SpVol | Both | KS | 0.27 | <0.001 | <0.001 | — | — | — |
| 3E | 2 h | 30 | SpVol | Both | KS | 0.28 | <0.001 | <0.001 | — | — | — |
| 3E | 2 h | 120 | SpVol | Both | KS | 0.25 | <0.001 | <0.001 | — | — | — |
| 3F | 30 min | 10 | ASlperi | Both | KS | 0.12 | 0.15 | 0.22 | — | — | — |
| 3F | 30 min | 30 | ASlperi | Both | KS | 0.12 | 0.093 | 0.22 | — | — | — |
| 3F | 30 min | 120 | ASlperi | Both | KS | 0.094 | 0.22 | 0.22 | — | — | — |
| 3F | 2 h | 10 | ASlperi | Both | KS | 0.3 | <0.001 | <0.001 | — | — | — |
| 3F | 2 h | 30 | ASlperi | Both | KS | 0.29 | <0.001 | <0.001 | — | — | — |
| 3F | 2 h | 120 | ASlperi | Both | KS | 0.25 | <0.001 | <0.001 | — | — | — |
| 3G | Both | 120 | SpVol | 1 vs. 2 | LMM | 41 | <0.001 | — | — | 0.61 | — |
| 3G | Both | 120 | ASlperi | 1 vs. 2 | LMM | 41 | <0.001 | — | — | 0.65 | — |
| 3G | 30 min | 120 | c1 vs. c2 | Both | $\chi^2$ | 8.8 | 0.003 | — | — | — | — |
| 3G | 2 h | 120 | c1 vs. c2 | Both | $\chi^2$ | 26 | <0.001 | — | — | — | — |
| 3G | 30 min | 10 | SpVol,<br>ASlperi | 1 | LMM | 1.8 | 0.093 | 0.18 | — | 0.35 | <0.001 |
| 3G | 30 min | 10 | SpVol,<br>ASlperi | 2 | LMM | 1.6 | 0.11 | 0.11 | — | 0.37 | <0.001 |
| 3G | 30 min | 30 | SpVol,<br>ASlperi | 1 | LMM | 1.2 | 0.27 | 0.27 | — | 0.36 | <0.001 |
| 3G | 30 min | 30 | SpVol,<br>ASlperi | 2 | LMM | 2 | 0.05 | 0.098 | — | 0.36 | <0.001 |
| 3G | 30 min | 120 | SpVol,<br>ASlperi | 1 | LMM | 1.7 | 0.12 | 0.18 | — | 0.32 | <0.001 |
| 3G | 30 min | 120 | SpVol,<br>ASlperi | 2 | LMM | 1.9 | 0.065 | 0.098 | — | 0.34 | <0.001 |
| 3G | 2 h | 10 | SpVol,<br>ASlperi | 1 | LMM | 1.4 | 0.17 | 0.22 | — | 0.28 | <0.001 |
| 3G | 2 h | 10 | SpVol,<br>ASlperi | 2 | LMM | 0.73 | 0.48 | 0.52 | — | 0.61 | <0.001 |
| 3G | 2 h | 30 | SpVol,<br>ASlperi | 1 | LMM | 1.3 | 0.22 | 0.22 | — | 0.27 | <0.001 |
| 3G | 2 h | 30 | SpVol,<br>ASlperi | 2 | LMM | 0.66 | 0.52 | 0.52 | — | 0.6 | <0.001 |
| 3G | 2 h | 120 | SpVol,<br>ASlperi | 1 | LMM | 1.6 | 0.13 | 0.22 | — | 0.27 | <0.001 |
| 3G | 2 h | 120 | SpVol,<br>ASlperi | 2 | LMM | 0.65 | 0.52 | 0.52 | — | 0.6 | <0.001 |

**Table S2.** Summary of statistical output comparing PSD area and PSD offset from the ASI center between control and LTP synapses in the MML. **Statistical outcomes for comparisons visualized in Figure 4. Cols** 1–3, 5, and 10– 12 are as described for Table S1. (Col 4) Response variable and covariate: PSD area (PSDarea), PSD offset from ASI center (PSDoffset), ASI area (ASIarea). If a significant interaction between the covariate and condition was detected, this is indicated by “(int).” (Col 6) Statistical test: Kolmogorov-Smirnov test (KS), Linear mixed model (LMM). (Col 7) Test statistic: D for KS test, t for LMM. (Col 8) p-value for condition effect, or for interaction effect if significant (as indicated in Col 4). (Col 9) p-value for condition effect or interaction effect adjusted using the Benjamini-Hochberg (BH) procedure.

| Fig. | Time | d<sub>ag</sub><br>(nm) | Var,<br>CoVar | Cluster | Test | Statistic | p (C) | P<br>(C, adj) | $\eta_p^2$ | R<sup>2</sup> | P<br>(CoVar) |
| --- | --- | --- | --- | --- | --- | --- | --- | --- | --- | --- | --- |
| 4C | 30 min | 10 | PSDarea | Both | KS | 0.13 | 0.11 | 0.11 | — | — | — |
| 4C | 30 min | 30 | PSDarea | Both | KS | 0.13 | 0.051 | 0.076 | — | — | — |
| 4C | 30 min | 120 | PSDarea | Both | KS | 0.18 | <0.001 | 0.002 | — | — | — |
| 4C | 2 h | 10 | PSDarea | Both | KS | 0.16 | 0.034 | 0.034 | — | — | — |
| 4C | 2 h | 30 | PSDarea | Both | KS | 0.18 | 0.0055 | 0.013 | — | — | — |
| 4C | 2 h | 120 | PSDarea | Both | KS | 0.16 | 0.0087 | 0.013 | — | — | — |
| 4D | 30 min | 10 | PSDoffset | Both | KS | 0.13 | 0.11 | 0.17 | — | — | — |
| 4D | 30 min | 30 | PSDoffset | Both | KS | 0.12 | 0.098 | 0.17 | — | — | — |
| 4D | 30 min | 120 | PSDoffset | Both | KS | 0.094 | 0.23 | 0.23 | — | — | — |
| 4D | 2 h | 10 | PSDoffset | Both | KS | 0.19 | 0.007 | 0.011 | — | — | — |
| 4D | 2 h | 30 | PSDoffset | Both | KS | 0.18 | 0.0066 | 0.011 | — | — | — |
| 4D | 2 h | 120 | PSDoffset | Both | KS | 0.13 | 0.038 | 0.038 | — | — | — |
| 4E | 30 min | 10 | PSDarea,<br>ASlarea | 1 | LMM | -1.4 | 0.16 | 0.16 | — | 0.54 | <0.001 |
| 4E | 30 min | 10 | PSDarea,<br>ASlarea | 2 | LMM | -0.38 | 0.71 | 0.81 | — | 0.28 | <0.001 |
| 4E | 30 min | 30 | PSDarea,<br>ASlarea | 1 | LMM | -1.6 | 0.13 | 0.16 | — | 0.55 | <0.001 |
| 4E | 30 min | 30 | PSDarea,<br>ASlarea | 2 | LMM | -0.31 | 0.76 | 0.81 | — | 0.27 | <0.001 |
| 4E | 30 min | 120 | PSDarea,<br>ASlarea | 1 | LMM | -1.9 | 0.078 | 0.16 | — | 0.52 | <0.001 |
| 4E | 30 min | 120 | PSDarea,<br>ASlarea | 2 | LMM | -0.25 | 0.81 | 0.81 | — | 0.29 | <0.001 |
| 4E | 2 h | 10 | PSDarea,<br>ASlarea | 1 | LMM | -1.6 | 0.14 | 0.14 | — | 0.31 | <0.001 |
| 4E | 2 h | 10 | PSDarea,<br>ASlarea | 2 | LMM | -2.5 | 0.027 | 0.081 | — | 0.73 | <0.001 |
| 4E | 2 h | 30 | PSDarea,<br>ASlarea | 1 | LMM | -1.7 | 0.11 | 0.14 | — | 0.34 | <0.001 |
| 4E | 2 h | 30 | PSDarea,<br>ASlarea | 2 | LMM | -1.9 | 0.075 | 0.083 | — | 0.54 | <0.001 |
| 4E | 2 h | 120 | PSDarea,<br>ASlarea | 1 | LMM | -1.7 | 0.1 | 0.14 | — | 0.33 | <0.001 |
| 4E | 2 h | 120 | PSDarea,<br>ASlarea | 2 | LMM | -1.8 | 0.083 | 0.083 | — | 0.55 | <0.001 |
| 4F | 30 min | 10 | PSDoffset,<br>ASlarea | 1 | LMM | 1.5 | 0.15 | 0.46 | — | 0.2 | <0.001 |
| 4F | 30 min | 10 | PSDoffset,<br>ASlarea | 2 | LMM | 0.46 | 0.65 | 0.65 | — | 0.12 | <0.001 |
| 4F | 30 min | 30 | PSDoffset,<br>ASlarea | 1 | LMM | 0.91 | 0.38 | 0.57 | — | 0.22 | <0.001 |
| 4F | 30 min | 30 | PSDoffset,<br>ASlarea | 2 | LMM | 0.51 | 0.61 | 0.65 | — | 0.11 | <0.001 |
| 4F | 30 min | 120 | PSDoffset,<br>ASlarea | 1 | LMM | 0.55 | 0.59 | 0.59 | — | 0.21 | <0.001 |
| 4F | 30 min | 120 | PSDoffset,<br>ASlarea | 2 | LMM | 0.66 | 0.51 | 0.65 | — | 0.079 | <0.001 |
| 4F | 2 h | 10 | PSDoffset,<br>ASlarea (int) | 1 | LMM | -2.4 | 0.019 | 0.023 | — | 0.26 | — |
| 4F | 2 h | 10 | PSDoffset,<br>ASlarea | 2 | LMM | 0.15 | 0.88 | 0.88 | — | 0.047 | 0.043 |
| 4F | 2 h | 30 | PSDoffset,<br>ASlarea (int) | 1 | LMM | -2.3 | 0.023 | 0.023 | — | 0.29 | — |
| 4F | 2 h | 30 | PSDoffset,<br>ASlarea | 2 | LMM | 0.39 | 0.7 | 0.88 | — | 0.074 | 0.0096 |
| 4F | 2 h | 120 | PSDoffset,<br>ASlarea | 1 | LMM | 0.53 | 0.6 | 0.6 | — | 0.21 | <0.001 |
| 4F | 2 h | 120 | PSDoffset,<br>ASlarea | 2 | LMM | 0.55 | 0.58 | 0.88 | — | 0.087 | 0.0065 |

**Table S3.** Summary of statistical output comparing length of astroglial surround, astroglial distance to the ASI perimeter, and astroglial access to the PSD between control and LTP synapses in the MML. **Statistical outcomes for comparisons visualized in Figure 5**. Cols 1–2, 5, and 10– 12 are as described for Table S1. (Col 3) Synapse subset defined by d_ag_: ≤ 10 nm, ≤ 30 nm, ≤ 120 nm. When the response variable was length of astroglial surround, only synapses with d_ag_ ≤ 120 nm were included, and Col 3 indicates the threshold used. (Col 4) Response variable and covariate: length of astroglia surround (l_ag_), astroglia distance (d_ag_) to the ASI perimeter, average distance from the astroglia-apposed ASI perimeter to the PSD (d_ag-PSD_), average distance from the overall ASI perimeter to the PSD (d_ASI-PSD_), ASI area. If a significant interaction between the covariate and condition was detected, this is indicated by “(int).” (Col 6) Statistical test: Kolmogorov-Smirnov test (KS), Linear mixed model (LMM). (Col 7) Test statistic: D for KS test, t for LMM. (Col 8) p-value for condition effect, or for interaction effect if significant (as indicated in Col 4). (Col 9) p-value for condition effect or interaction effect adjusted using the Benjamini-Hochberg (BH) procedure.

| Fig. | Time | d<sub>ag</sub><br>(nm) | Var,<br>CoVar | Cluster | Test | Statistic | p (C) | p<br>(C, adj) | $\eta_p^2$ | R<sup>2</sup> | p<br>(CoVar) |
| --- | --- | --- | --- | --- | --- | --- | --- | --- | --- | --- | --- |
| 5B | 30 min | 10 | $l_{ag}$,<br>ASlperi | 1 | LMM | 0.42 | 0.68 | 0.71 | — | 0.059 | <0.001 |
| 5B | 30 min | 10 | $l_{ag}$,<br>ASlperi | 2 | LMM | -2.9 | 0.0038 | 0.011 | 0.054 | 0.2 | <0.001 |
| 5B | 30 min | 30 | $l_{ag}$,<br>ASlperi | 1 | LMM | -0.37 | 0.71 | 0.71 | — | 0.059 | <0.001 |
| 5B | 30 min | 30 | $l_{ag}$,<br>ASlperi | 2 | LMM | -2.3 | 0.022 | 0.022 | 0.036 | 0.19 | <0.001 |
| 5B | 30 min | 120 | $l_{ag}$,<br>ASlperi | 1 | LMM | -0.54 | 0.6 | 0.71 | — | 0.27 | <0.001 |
| 5B | 30 min | 120 | $l_{ag}$,<br>ASlperi | 2 | LMM | -2.5 | 0.012 | 0.018 | 0.042 | 0.26 | <0.001 |
| 5C | 2 h | 10 | $l_{ag}$,<br>ASlperi | 1 | LMM | -0.52 | 0.61 | 0.61 | — | 0.056 | <0.001 |
| 5C | 2 h | 10 | $l_{ag}$,<br>ASlperi | 2 | LMM | -2.4 | 0.017 | 0.025 | 0.054 | 0.16 | <0.001 |
| 5C | 2 h | 30 | $l_{ag}$,<br>ASlperi | 1 | LMM | -0.76 | 0.45 | 0.61 | — | 0.07 | <0.001 |
| 5C | 2 h | 30 | $l_{ag}$,<br>ASlperi | 2 | LMM | -2.8 | 0.007 | 0.021 | 0.068 | 0.26 | <0.001 |
| 5C | 2 h | 120 | $l_{ag}$,<br>ASlperi | 1 | LMM | 0.9 | 0.38 | 0.61 | — | 0.24 | <0.001 |
| 5C | 2 h | 120 | $l_{ag}$,<br>ASlperi | 2 | LMM | -2.1 | 0.047 | 0.047 | 0.042 | 0.34 | <0.001 |
| 5E | 30 min | 10 | d<sub>ag</sub>,<br>ASlperi | 1 | LMM | -1.3 | 0.2 | 0.3 | — | 0.042 | 0.055 |
| 5E | 30 min | 10 | d<sub>ag</sub>,<br>ASlperi | 2 | LMM | 0.88 | 0.38 | 0.38 | — | 0.0063 | 0.94 |
| 5E | 30 min | 30 | $d_{ag,}$<br>ASlperi | 1 | LMM | -1.6 | 0.13 | 0.3 | — | 0.026 | 0.11 |
| 5E | 30 min | 30 | $d_{ag,}$<br>ASlperi | 2 | LMM | 1.3 | 0.21 | 0.31 | — | 0.016 | 0.66 |
| 5E | 30 min | 120 | $d_{ag,}$<br>ASlperi | 1 | LMM | -0.096 | 0.92 | 0.92 | — | 1.00E-04 | 0.91 |
| 5E | 30 min | 120 | $d_{ag,}$<br>ASlperi | 2 | LMM | 1.6 | 0.12 | 0.31 | — | 0.05 | 0.07 |
| 5F | 2 h | 10 | $d_{ag,}$<br>ASlperi | 1 | LMM | -0.62 | 0.54 | 0.79 | — | 0.034 | 0.021 |
| 5F | 2 h | 10 | $d_{ag,}$<br>ASlperi | 2 | LMM | 3.7 | <0.001 | <0.001 | 0.14 | 0.14 | 0.88 |
| 5F | 2 h | 30 | $d_{ag,}$<br>ASlperi | 1 | LMM | -0.26 | 0.79 | 0.79 | — | 0.0039 | 0.47 |
| 5F | 2 h | 30 | $d_{ag,}$<br>ASlperi | 2 | LMM | 4 | <0.001 | <0.001 | 0.14 | 0.15 | 0.88 |
| 5F | 2 h | 120 | $d_{ag,}$<br>ASlperi | 1 | LMM | 0.78 | 0.44 | 0.79 | — | 0.002 | 0.77 |
| 5F | 2 h | 120 | $d_{ag,}$<br>ASlperi | 2 | LMM | 3.5 | <0.001 | <0.001 | 0.11 | 0.14 | 0.42 |
| 5H | 30 min | 10 | $d_{ag-psd}$ | Both | KS | 0.14 | 0.089 | 0.089 | — | — | — |
| 5H | 30 min | 30 | $d_{ag-psd}$ | Both | KS | 0.14 | 0.032 | 0.089 | — | — | — |
| 5H | 30 min | 120 | $d_{ag-psd}$ | Both | KS | 0.12 | 0.065 | 0.089 | — | — | — |
| 5H | 2 h | 10 | $d_{ag-psd}$ | Both | KS | 0.29 | <0.001 | <0.001 | — | — | — |
| 5H | 2 h | 30 | $d_{ag-psd}$ | Both | KS | 0.28 | <0.001 | <0.001 | — | — | — |
| 5H | 2 h | 120 | $d_{ag-psd}$ | Both | KS | 0.25 | <0.001 | <0.001 | — | — | — |
| 5I | 30 min | 10 | $d_{ag-psd,}$<br>$d_{ASl-psd}$ | 1 | LMM | 0.28 | 0.78 | 0.9 | — | 0.91 | <0.001 |
| 5I | 30 min | 10 | $d_{ag-psd,}$<br>$d_{ASl-psd}$ | 2 | LMM | -1.4 | 0.17 | 0.2 | — | 0.74 | <0.001 |
| 5I | 30 min | 30 | $d_{ag-psd,}$<br>$d_{ASl-psd}$ | 1 | LMM | 0.85 | 0.42 | 0.9 | — | 0.88 | <0.001 |
| 5I | 30 min | 30 | $d_{ag-psd,}$<br>$d_{ASl-psd}$ | 2 | LMM | -1.3 | 0.2 | 0.2 | — | 0.73 | <0.001 |
| 5I | 30 min | 120 | $d_{ag-psd}$<br>$d_{ASI-psd}$ | 1 | LMM | 0.13 | 0.9 | 0.9 | — | 0.78 | <0.001 |
| 5I | 30 min | 120 | $d_{ag-psd}$<br>$d_{ASI-psd}$ | 2 | LMM | -1.4 | 0.17 | 0.2 | — | 0.73 | <0.001 |
| 5I | 2 h | 10 | $d_{ag-psd}$<br>$d_{ASI-psd}$ | 1 | LMM | 0.57 | 0.57 | 1 | — | 0.89 | <0.001 |
| 5I | 2 h | 10 | $d_{ag-psd}$<br>$d_{ASI-psd}$ | 2 | LMM | -0.3 | 0.77 | 0.99 | — | 0.73 | <0.001 |
| 5I | 2 h | 30 | $d_{ag-psd}$<br>$d_{ASI-psd}$ | 1 | LMM | 0.36 | 0.72 | 1 | — | 0.87 | <0.001 |
| 5I | 2 h | 30 | $d_{ag-psd}$<br>$d_{ASI-psd}$ | 2 | LMM | -0.012 | 0.99 | 0.99 | — | 0.73 | <0.001 |
| 5I | 2 h | 120 | $d_{ag-psd}$<br>$d_{ASI-psd}$ | 1 | LMM | 0.0044 | 1 | 1 | — | 0.82 | <0.001 |
| 5I | 2 h | 120 | $d_{ag-psd}$<br>$d_{ASI-psd}$ | 2 | LMM | 0.23 | 0.82 | 0.99 | — | 0.74 | <0.001 |
| 5J | 30 min | 10 | $d_{ag-psd}$<br>ASlarea | 1 | LMM | 1.3 | 0.21 | 0.3 | — | 0.5 | <0.001 |
| 5J | 30 min | 10 | $d_{ag-psd}$<br>ASlarea (int) | 2 | LMM | -2.1 | 0.035 | 0.035 | — | 0.37 | — |
| 5J | 30 min | 30 | $d_{ag-psd}$<br>ASlarea | 1 | LMM | 1.2 | 0.26 | 0.3 | — | 0.45 | <0.001 |
| 5J | 30 min | 30 | $d_{ag-psd}$<br>ASlarea | 2 | LMM | -0.014 | 0.99 | 0.99 | — | 0.32 | <0.001 |
| 5J | 30 min | 120 | $d_{ag-psd}$<br>ASlarea | 1 | LMM | 1.1 | 0.3 | 0.3 | — | 0.43 | <0.001 |
| 5J | 30 min | 120 | $d_{ag-psd}$<br>ASlarea | 2 | LMM | -0.39 | 0.71 | 0.99 | — | 0.3 | <0.001 |
| 5J | 2 h | 10 | $d_{ag-psd}$<br>ASlarea | 1 | LMM | 2.6 | 0.019 | 0.056 | 0.03 | 0.5 | <0.001 |
| 5J | 2 h | 10 | $d_{ag-psd}$<br>ASlarea | 2 | LMM | 0.9 | 0.37 | 0.37 | — | 0.22 | <0.001 |
| 5J | 2 h | 30 | $d_{ag-psd}$<br>ASlarea | 1 | LMM | 1.9 | 0.071 | 0.093 | — | 0.47 | <0.001 |
| 5J | 2 h | 30 | $d_{ag-psd}$,<br>ASlarea | 2 | LMM | 1.3 | 0.22 | 0.34 | — | 0.24 | <0.001 |
| 5J | 2 h | 120 | $d_{ag-psd}$,<br>ASlarea | 1 | LMM | 1.8 | 0.093 | 0.093 | — | 0.38 | <0.001 |
| 5J | 2 h | 120 | $d_{ag-psd}$,<br>ASlarea | 2 | LMM | 1.5 | 0.13 | 0.34 | — | 0.27 | <0.001 |

**Table S4.** Summary of statistical output comparing spine volume, ASI perimeter, and astroglial presence at the ASI between control and cLTD synapses in the OML. **Statistical outcomes for comparisons visualized in Figure 6**. Cols 1–12 are as described for Table S1.

| Fig. | Time | d<sub>ag</sub><br>(nm) | Var,<br>CoVar | Cluster | Test | Statistic | p (C) | p<br>(C, adj) | $\eta_p^2$ | R<sup>2</sup> | p<br>(CoVar) |
| --- | --- | --- | --- | --- | --- | --- | --- | --- | --- | --- | --- |
| 6D | 30 min | 120 | ag- vs. ag+ | Both | $\chi^2$ | 3.3 | 0.069 | — | — | — | — |
| 6D | 2 h | 120 | ag- vs. ag+ | Both | $\chi^2$ | 3.7 | 0.054 | — | — | — | — |
| 6E | 30 min | 10 | SpVol | Both | KS | 0.14 | 0.11 | 0.11 | — | — | — |
| 6E | 30 min | 30 | SpVol | Both | KS | 0.14 | 0.038 | 0.057 | — | — | — |
| 6E | 30 min | 120 | SpVol | Both | KS | 0.13 | 0.028 | 0.057 | — | — | — |
| 6E | 2 h | 10 | SpVol | Both | KS | 0.1 | 0.38 | 0.38 | — | — | — |
| 6E | 2 h | 30 | SpVol | Both | KS | 0.1 | 0.25 | 0.37 | — | — | — |
| 6E | 2 h | 120 | SpVol | Both | KS | 0.1 | 0.21 | 0.37 | — | — | — |
| 6F | 30 min | 10 | ASlperi | Both | KS | 0.12 | 0.26 | 0.26 | — | — | — |
| 6F | 30 min | 30 | ASlperi | Both | KS | 0.13 | 0.09 | 0.13 | — | — | — |
| 6F | 30 min | 120 | ASlperi | Both | KS | 0.16 | 0.0042 | 0.013 | — | — | — |
| 6F | 2 h | 10 | ASlperi | Both | KS | 0.044 | 1 | 1 | — | — | — |
| 6F | 2 h | 30 | ASlperi | Both | KS | 0.055 | 0.94 | 1 | — | — | — |
| 6F | 2 h | 120 | ASlperi | Both | KS | 0.055 | 0.89 | 1 | — | — | — |
| 6G | Both | 120 | SpVol | 1 vs. 2 | LMM | 32 | <0.001 | — | — | 0.47 | — |
| 6G | Both | 120 | ASlperi | 1 vs. 2 | LMM | 31 | <0.001 | — | — | 0.49 | — |
| 6G | 30 min | 120 | c1 vs. c2 | Both | $\chi^2$ | 1 | 0.32 | — | — | — | — |
| 6G | 2 h | 120 | c1 vs. c2 | Both | $\chi^2$ | 0.088 | 0.77 | — | — | — | — |
| 6G | 30 min | 10 | SpVol,<br>ASlperi | 1 | LMM | -0.23 | 0.82 | 0.92 | — | 0.26 | <0.001 |
| 6G | 30 min | 10 | SpVol,<br>ASlperi | 2 | LMM | 0.26 | 0.8 | 0.91 | — | 0.17 | <0.001 |
| 6G | 30 min | 30 | SpVol,<br>ASlperi | 1 | LMM | -0.1 | 0.92 | 0.92 | — | 0.26 | <0.001 |
| 6G | 30 min | 30 | SpVol,<br>ASlperi | 2 | LMM | 0.11 | 0.91 | 0.91 | — | 0.16 | <0.001 |
| 6G | 30 min | 120 | SpVol,<br>ASlperi | 1 | LMM | -0.23 | 0.82 | 0.92 | — | 0.25 | <0.001 |
| 6G | 30 min | 120 | SpVol,<br>ASlperi | 2 | LMM | 0.11 | 0.91 | 0.91 | — | 0.16 | <0.001 |
| 6G | 2 h | 10 | SpVol,<br>ASlperi | 1 | LMM | 0.093 | 0.93 | 0.93 | — | 0.2 | <0.001 |
| 6G | 2 h | 10 | SpVol,<br>ASlperi | 2 | LMM | 0.63 | 0.54 | 0.54 | — | 0.7 | <0.001 |
| 6G | 2 h | 30 | SpVol,<br>ASlperi | 1 | LMM | 0.22 | 0.83 | 0.93 | — | 0.2 | <0.001 |
| 6G | 2 h | 30 | SpVol,<br>ASlperi | 2 | LMM | 0.7 | 0.5 | 0.54 | — | 0.7 | <0.001 |
| 6G | 2 h | 120 | SpVol,<br>ASlperi | 1 | LMM | 0.21 | 0.84 | 0.93 | — | 0.22 | <0.001 |
| 6G | 2 h | 120 | SpVol,<br>ASlperi | 2 | LMM | 1 | 0.31 | 0.54 | — | 0.69 | <0.001 |

**Table S5.**
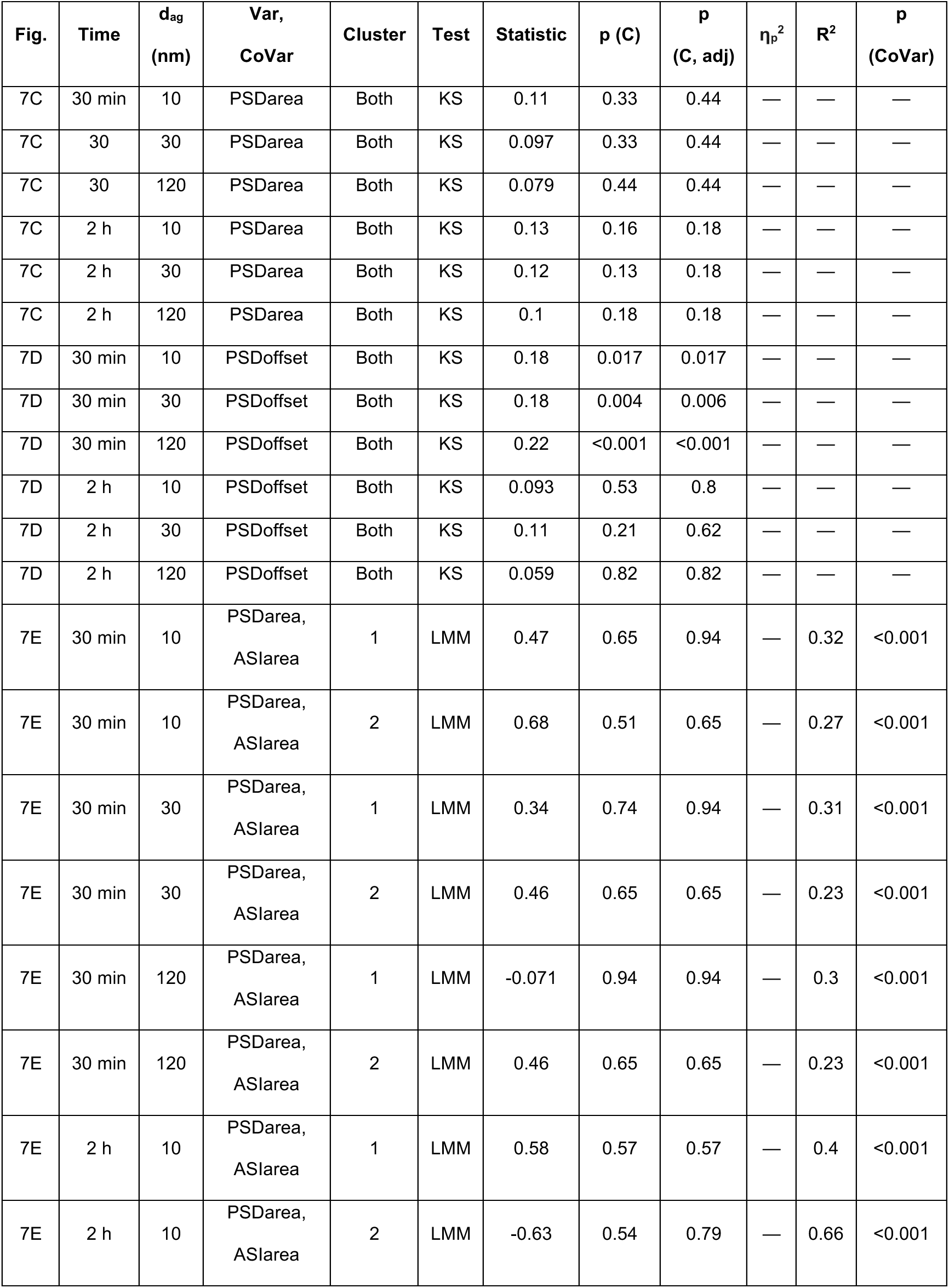
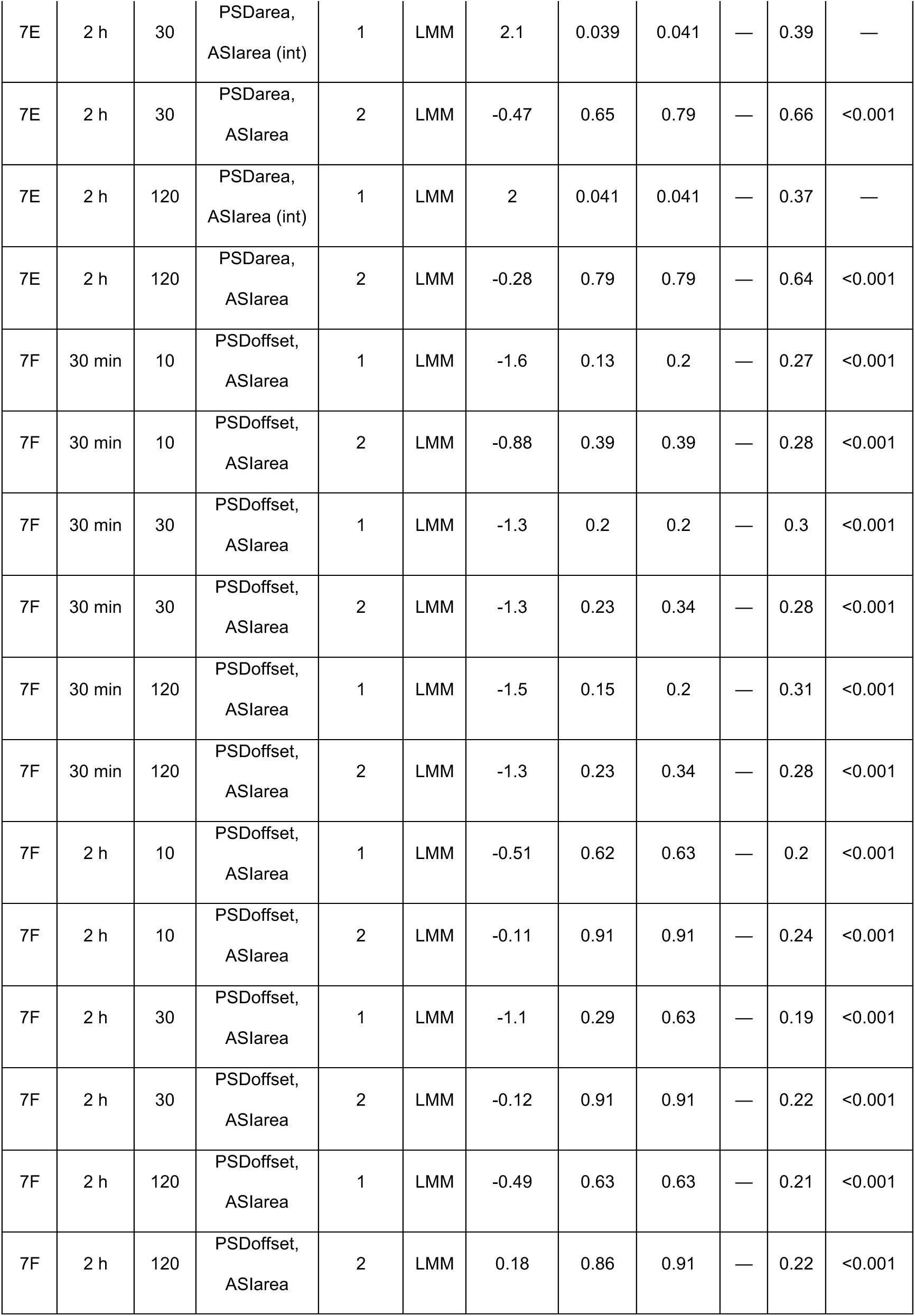
Summary of statistical output comparing PSD area and PSD offset from the ASI center between control and cLTD synapses in the OML. **Statistical outcomes for comparisons visualized in Figure 7**. Cols 1–3, 5, and 10– 12 are as described for Table S1. (Col 4) Response variable and covariate: PSD area, PSD offset from ASI center, ASI area. If a significant interaction between the covariate and condition was detected, this is indicated by “(int).” (Col 6) Statistical test: Kolmogorov-Smirnov test (KS), Linear mixed model (LMM). (Col 7) Test statistic: D for KS test, t for LMM. (Col 8) p-value for condition effect, or for interaction effect if significant (as indicated in Col 4). (Col 9) p-value for condition effect or interaction effect adjusted using the Benjamini-Hochberg (BH) procedure.

**Table S6.** Summary of statistical output comparing length of astroglial surround, astroglial distance to the ASI perimeter, and astroglial access to the PSD between control and cLTD synapses in the OML. **Statistical outcomes for comparisons visualized in Figure 8**. Cols 1–2, 5, and 10– 12 are as described for Table S1. (Col 3) Synapse subset defined by d_ag_: ≤ 10 nm, ≤ 30 nm, ≤ 120 nm. When the response variable was length of astroglial surround, only synapses with d_ag_ ≤ 120 nm were included, and Col 3 indicates the threshold used. (Col 4) Response variable and covariate: l_ag_, d_ag_, ASI perimeter, d_ag-PSD_, d_ASI-PSD_, ASI area. If a significant interaction between the covariate and condition was detected, this is indicated by “(int).” (Col 6) Statistical test: Kolmogorov-Smirnov test (KS), Linear mixed model (LMM). (Col 7) Test statistic: D for KS test, t for LMM. (Col 8) p-value for condition effect, or for interaction effect if significant (as indicated in Col 4). (Col 9) p-value for condition or interaction effect adjusted using the Benjamini-Hochberg (BH) procedure.

| Fig. | Time | d<sub>ag</sub><br>(nm) | Var,<br>CoVar | Cluster | Test | Statistic | p (C) | p<br>(C, adj) | $\eta_p^2$ | R<sup>2</sup> | p<br>(CoVar) |
| --- | --- | --- | --- | --- | --- | --- | --- | --- | --- | --- | --- |
| 8B | 30 min | 10 | $l_{ag}$,<br>ASlperi | 1 | LMM | 0.41 | 0.68 | 0.71 | — | 0.082 | <0.001 |
| 8B | 30 min | 10 | $l_{ag}$,<br>ASlperi | 2 | LMM | -0.31 | 0.76 | 0.76 | — | 0.06 | 0.058 |
| 8B | 30 min | 30 | $l_{ag}$,<br>ASlperi | 1 | LMM | 0.37 | 0.71 | 0.71 | — | 0.11 | <0.001 |
| 8B | 30 min | 30 | $l_{ag}$,<br>ASlperi | 2 | LMM | -1.2 | 0.25 | 0.37 | — | 0.16 | 0.0059 |
| 8B | 30 min | 120 | $l_{ag}$,<br>ASlperi | 1 | LMM | 0.73 | 0.48 | 0.71 | — | 0.34 | <0.001 |
| 8B | 30 min | 120 | $l_{ag}$,<br>ASlperi | 2 | LMM | -1.4 | 0.2 | 0.37 | — | 0.26 | <0.001 |
| 8C | 2 h | 10 | $l_{ag}$,<br>ASlperi | 1 | LMM | 0.58 | 0.57 | 0.57 | — | 0.033 | <0.001 |
| 8C | 2 h | 10 | $l_{ag}$,<br>ASlperi (int) | 2 | LMM | -2.6 | 0.012 | 0.018 | — | 0.19 | — |
| 8C | 2 h | 30 | $l_{ag}$,<br>ASlperi | 1 | LMM | 0.82 | 0.41 | 0.57 | — | 0.032 | <0.001 |
| 8C | 2 h | 30 | $l_{ag}$,<br>ASlperi (int) | 2 | LMM | -2.3 | 0.023 | 0.023 | — | 0.3 | — |
| 8C | 2 h | 120 | $l_{ag}$,<br>ASlperi | 1 | LMM | 0.64 | 0.54 | 0.57 | — | 0.19 | <0.001 |
| 8C | 2 h | 120 | $l_{ag}$,<br>ASlperi (int) | 2 | LMM | -3.1 | 0.0028 | 0.0085 | — | 0.47 | — |
| 8E | 30 min | 10 | d<sub>ag</sub>,<br>ASlperi (int) | 1 | LMM | 2 | 0.048 | 0.048 | — | 0.061 | — |
| 8E | 30 min | 10 | d<sub>ag</sub>,<br>ASlperi | 2 | LMM | -0.44 | 0.67 | 0.67 | — | 0.022 | 0.86 |
| 8E | 30 min | 30 | $d_{ag}$,<br>ASlperi | 1 | LMM | -2.6 | 0.01 | 0.02 | 0.02 | 0.029 | 0.14 |
| 8E | 30 min | 30 | $d_{ag}$,<br>ASlperi | 2 | LMM | 0.48 | 0.64 | 0.67 | — | 0.0076 | 0.86 |
| 8E | 30 min | 120 | $d_{ag}$,<br>ASlperi | 1 | LMM | -1.7 | 0.084 | 0.084 | — | 0.013 | 0.14 |
| 8E | 30 min | 120 | $d_{ag}$,<br>ASlperi | 2 | LMM | 0.48 | 0.64 | 0.67 | — | 0.0076 | 0.86 |
| 8F | 2 h | 10 | $d_{ag}$,<br>ASlperi (int) | 1 | LMM | 2.1 | 0.035 | 0.036 | — | 0.046 | — |
| 8F | 2 h | 10 | $d_{ag}$,<br>ASlperi | 2 | LMM | 1.2 | 0.22 | 0.33 | — | 0.028 | 0.68 |
| 8F | 2 h | 30 | $d_{ag}$,<br>ASlperi (int) | 1 | LMM | 2.1 | 0.036 | 0.036 | — | 0.017 | — |
| 8F | 2 h | 30 | $d_{ag}$,<br>ASlperi | 2 | LMM | 0.73 | 0.47 | 0.47 | — | 0.017 | 0.68 |
| 8F | 2 h | 120 | $d_{ag}$,<br>ASlperi | 1 | LMM | -0.3 | 0.77 | 0.77 | — | 0.0019 | 0.45 |
| 8F | 2 h | 120 | $d_{ag}$,<br>ASlperi | 2 | LMM | 1.3 | 0.19 | 0.33 | — | 0.034 | 0.68 |
| 8H | 30 min | 10 | $d_{ag-psd}$ | Both | KS | 0.14 | 0.097 | 0.097 | — | — | — |
| 8H | 30 min | 30 | $d_{ag-psd}$ | Both | KS | 0.14 | 0.047 | 0.07 | — | — | — |
| 8H | 30 min | 120 | $d_{ag-psd}$ | Both | KS | 0.16 | 0.0041 | 0.012 | — | — | — |
| 8H | 2 h | 10 | $d_{ag-psd}$ | Both | KS | 0.096 | 0.49 | 0.49 | — | — | — |
| 8H | 2 h | 30 | $d_{ag-psd}$ | Both | KS | 0.1 | 0.29 | 0.49 | — | — | — |
| 8H | 2 h | 120 | $d_{ag-psd}$ | Both | KS | 0.085 | 0.4 | 0.49 | — | — | — |
| 8I | 30 min | 10 | $d_{ag-psd}$,<br>$d_{ASl-psd}$ (int) | 1 | LMM | 2.8 | 0.0058 | 0.012 | — | 0.86 | — |
| 8I | 30 min | 10 | $d_{ag-psd}$,<br>$d_{ASl-psd}$ | 2 | LMM | 0.8 | 0.45 | 0.71 | — | 0.91 | <0.001 |
| 8I | 30 min | 30 | $d_{ag-psd}$,<br>$d_{ASl-psd}$ | 1 | LMM | 1.2 | 0.24 | 0.24 | — | 0.82 | <0.001 |
| 8I | 30 min | 30 | $d_{ag-psd}$,<br>$d_{ASl-psd}$ | 2 | LMM | -0.37 | 0.71 | 0.71 | — | 0.88 | <0.001 |
| 8I | 30 min | 120 | $d_{ag-psd}$<br>$d_{ASI-psd} (int)$ | 1 | LMM | 2.2 | 0.028 | 0.028 | — | 0.82 | — |
| 8I | 30 min | 120 | $d_{ag-psd}$<br>$d_{ASI-psd}$ | 2 | LMM | -0.37 | 0.71 | 0.71 | — | 0.88 | <0.001 |
| 8I | 2 h | 10 | $d_{ag-psd}$<br>$d_{ASI-psd}$ | 1 | LMM | -0.78 | 0.44 | 0.53 | — | 0.88 | <0.001 |
| 8I | 2 h | 10 | $d_{ag-psd}$<br>$d_{ASI-psd}$ | 2 | LMM | 2 | 0.049 | 0.073 | — | 0.87 | <0.001 |
| 8I | 2 h | 30 | $d_{ag-psd}$<br>$d_{ASI-psd}$ | 1 | LMM | -0.63 | 0.53 | 0.53 | — | 0.87 | <0.001 |
| 8I | 2 h | 30 | $d_{ag-psd}$<br>$d_{ASI-psd}$ | 2 | LMM | 2.3 | 0.026 | 0.073 | — | 0.86 | <0.001 |
| 8I | 2 h | 120 | $d_{ag-psd}$<br>$d_{ASI-psd} (int)$ | 1 | LMM | -2.2 | 0.029 | 0.029 | — | 0.78 | — |
| 8I | 2 h | 120 | $d_{ag-psd}$<br>$d_{ASI-psd}$ | 2 | LMM | 1.3 | 0.18 | 0.18 | — | 0.82 | <0.001 |
| 8J | 30 min | 10 | $d_{ag-psd}$<br>ASlarea | 1 | LMM | -1.1 | 0.27 | 0.33 | — | 0.32 | <0.001 |
| 8J | 30 min | 10 | $d_{ag-psd}$<br>ASlarea | 2 | LMM | 0.17 | 0.87 | 0.87 | — | 0.48 | <0.001 |
| 8J | 30 min | 30 | $d_{ag-psd}$<br>ASlarea | 1 | LMM | -1 | 0.33 | 0.33 | — | 0.3 | <0.001 |
| 8J | 30 min | 30 | $d_{ag-psd}$<br>ASlarea | 2 | LMM | -0.23 | 0.82 | 0.87 | — | 0.48 | <0.001 |
| 8J | 30 min | 120 | $d_{ag-psd}$<br>ASlarea | 1 | LMM | -1.2 | 0.27 | 0.33 | — | 0.31 | <0.001 |
| 8J | 30 min | 120 | $d_{ag-psd}$<br>ASlarea | 2 | LMM | -0.23 | 0.82 | 0.87 | — | 0.48 | <0.001 |
| 8J | 2 h | 10 | $d_{ag-psd}$<br>ASlarea (int) | 1 | LMM | -3.8 | <0.001 | <0.001 | — | 0.36 | — |
| 8J | 2 h | 10 | $d_{ag-psd}$<br>ASlarea (int) | 2 | LMM | -2 | 0.048 | 0.048 | — | 0.56 | — |
| 8J | 2 h | 30 | $d_{ag-psd}$<br>ASlarea (int) | 1 | LMM | -3.7 | <0.001 | <0.001 | — | 0.36 | — |
| 8J | 2 h | 30 | $d_{ag-psd}$,<br>ASlarea (int) | 2 | LMM | -2.3 | 0.023 | 0.045 | — | 0.56 | — |
| 8J | 2 h | 120 | $d_{ag-psd}$,<br>ASlarea (int) | 1 | LMM | -3.5 | <0.001 | <0.001 | — | 0.36 | — |
| 8J | 2 h | 120 | $d_{ag-psd}$,<br>ASlarea | 2 | LMM | 1.8 | 0.091 | 0.091 | — | 0.5 | <0.001 |

**Table S7.** Summary of statistical output comparing ultrastructure of MML and OML tripartite synapses under control conditions. **Statistical outcomes for comparisons visualized in Figure 9**. Cols 1–3, 5–7, and 11–12 are as described for Table S1. (Col 4) Response variable and covariate: PAP present or absent at ASI, c1, c2, ASI perimeter, PSD area, l_ag_, d_ag_, PSD offset from the ASI center, d_ag-PSD_. If a significant interaction between the covariate and layer was detected, this is indicated by “(int).” (Col 8) p-value for layer (L) effect, or for interaction effect if significant (as indicated in Col 4). (Col 9) p-value for layer or interaction effect adjusted using the Benjamini-Hochberg (BH) procedure. (Col 10) Proportion of variance in the dependent variable explained by layer effect (partial eta², η_p_^2^).

| Fig. | Time | d<sub>ag</sub><br>(nm) | Var,<br>CoVar | Cluster | Test | Statistic | p (L) | p<br>(L, adj) | $\eta_p^2$ | R<sup>2</sup> | p<br>(CoVar) |
| --- | --- | --- | --- | --- | --- | --- | --- | --- | --- | --- | --- |
| 9A | Both | 120 | ag- vs. ag+ | Both | $\chi^2$ | 1.3 | 0.25 | — | — | — | — |
| 9B | Both | 120 | c1 vs. c2 | Both | $\chi^2$ | 5.3 | 0.022 | — | — | — | — |
| 9C | Both | 120 | ASlperi | Both | KS | 0.084 | 0.068 | — | — | — | — |
| 9D | Both | 120 | PSDarea | Both | KS | 0.084 | 0.068 | — | — | — | — |
| 9E | Both | 120 | <sup>lag</sup>,<br>ASlPeri | 1 | LMM | -1.1 | 0.29 | — | — | 0.23 | <0.001 |
| 9E | Both | 120 | <sup>lag</sup>,<br>ASlPeri | 2 | LMM | -0.25 | 0.81 | — | — | 0.35 | <0.001 |
| 9F | Both | 120 | <sup>dag</sup>,<br>ASlPeri (int) | 1 | LMM | -2.1 | 0.038 | — | — | 0.0056 | — |
| 9F | Both | 120 | <sup>dag</sup>,<br>ASlPeri | 2 | LMM | 0.59 | 0.56 | — | — | 0.016 | 0.14 |
| 9G | Both | 120 | PSDoffset | Both | KS | 0.055 | 0.47 | — | — | — | — |
| 9H | Both | 120 | <sup>dag-psd</sup> | Both | KS | 0.069 | 0.2 | — | — | — | — |

**Table S8.** Summary of statistical output comparing ultrastructure of tripartite synapses in the MML during LTP and in the OML during cLTD. **Statistical outcomes for comparisons visualized in Figure 10**. Cols 1-3, 5, 10-12 are as described for Table S1. (Col 4) Response variable and covariate(s): ASI perimeter, PSD area, l_ag_, d_ag_, PSD offset from the ASI center, d_ag-PSD_. For l_ag_ and d_ag_, “cond × layer (int)” indicates the tested interaction between condition and layer, controlling for ASI perimeter. (Col 6) Statistical test: Kolmogorov-Smirnov test (KS), Linear mixed model (LMM). (Col 7) Test statistic: D for KS test, t for LMM. (Col 8) p-value for condition effect or for interaction effect (if indicated in Col 4). (Col 9) p-value for condition effect or interaction effect adjusted using the Benjamini-Hochberg (BH) procedure.

| Fig. | Time | d<sub>ag</sub><br>(nm) | Var,<br>CoVar | Cluster | Test | Statistic | p (C) | p<br>(C, adj) | $\eta_p^2$ | R<sup>2</sup> | p<br>(CoVar) |
| --- | --- | --- | --- | --- | --- | --- | --- | --- | --- | --- | --- |
| 10A | 30 min | 120 | ASlperi | Both | KS | 0.25 | <0.001 | — | — | — | — |
| 10A | 2 h | 120 | ASlperi | Both | KS | 0.2 | <0.001 | — | — | — | — |
| 10B | 30 min | 120 | PSDarea | Both | KS | 0.1 | 0.16 | — | — | — | — |
| 10B | 2 h | 120 | PSDarea | Both | KS | 0.14 | 0.034 | — | — | — | — |
| 10C | 30 min | 120 | l<sub>ag</sub>, ASlperi,<br>cond x layer (int) | 1 | LMM | 0.65 | 0.52 | — | — | 0.31 | — |
| 10C | 30 min | 120 | l<sub>ag</sub>, ASlperi,<br>cond x layer (int) | 2 | LMM | 0.52 | 0.6 | — | — | 0.26 | — |
| 10C | 2 h | 120 | l<sub>ag</sub>, ASlperi,<br>cond x layer (int) | 1 | LMM | -0.013 | 0.99 | — | — | 0.21 | — |
| 10C | 2 h | 120 | l<sub>ag</sub>, ASlperi,<br>cond x layer (int) | 2 | LMM | 0.52 | 0.61 | — | — | 0.36 | — |
| 10D | 30 min | 120 | d<sub>ag</sub>, ASlperi,<br>cond x layer (int) | 1 | LMM | -0.85 | 0.4 | — | — | 0.0046 | — |
| 10D | 30 min | 120 | d<sub>ag</sub>, ASlperi,<br>cond x layer (int) | 2 | LMM | -1.2 | 0.24 | — | — | 0.049 | — |
| 10D | 2 h | 120 | d<sub>ag</sub>, ASlperi,<br>cond x layer (int) | 1 | LMM | -0.62 | 0.54 | — | — | 0.003 | — |
| 10D | 2 h | 120 | d<sub>ag</sub>, ASlperi,<br>cond x layer (int) | 2 | LMM | -1.5 | 0.14 | — | — | 0.097 | — |
| 10E | 30 min | 120 | PSDoffset | Both | KS | 0.28 | <0.001 | — | — | — | — |
| 10E | 2 h | 120 | PSDoffset | Both | KS | 0.15 | 0.014 | — | — | — | — |
| 10F | 30 min | 120 | d<sub>ag-psd</sub> | Both | KS | 0.26 | <0.001 | — | — | — | — |
| 10F | 2 h | 120 | d<sub>ag-psd</sub> | Both | KS | 0.21 | <0.001 | — | — | — | — |

**Table S9.** Summary of statistical outputs comparing the mean spine volume, ASI perimeter, length of astroglial surround, and astroglial distance to the ASI perimeter between control and LTP conditions in the MML. **Statistical outcomes for comparisons visualized in Figure S1. Table columns (col):** (Col 1) Figure panel. (Col 2) Time following DBS onset: 30 min, 2 h. (Col 3) Synapse subset defined by d_ag_ ≤ 120 nm. (Col 4) Response variable (Var) and covariate (CoVar): Spine volume (spVol), PAP present or absent from the ASI within 120 nm (ASIcat), fraction of astroglia surround at the ASI perimeter with 120 nm (l_ag_Frac), astroglial distance to the ASI perimeter (d_ag_). (Col 5) Synapse cluster: cluster 1 and 2 combined (both). (Col 6) Statistical test: Linear mixed model (LMM). (Col 7) Test statistic: t-value. (Col 8) p-value for condition (C) effect. (Col 9) Proportion of variance in the dependent variable explained by condition effect (partial eta², η_p_^2^). (Col 10) Coefficient of determination (R^2^). (Col 11) p-value for covariate effect.

| Fig. | Time | $d_{ag}$<br>(nm) | Var,<br>CoVar | Cluster | Test | Statistic | p (C) | $\eta_p^2$ (C) | R<sup>2</sup> | p (CoVar) |
| --- | --- | --- | --- | --- | --- | --- | --- | --- | --- | --- |
| S1A | 30 min | 120 | SpVol,<br>ASlcat | Both | LMM | 0.35 | 0.73 | — | 0.035 | <0.001 |
| S1A | 2 h | 120 | SpVol,<br>ASlcat | Both | LMM | 4.1 | <0.001 | 0.033 | 0.054 | 0.0099 |
| S1B | 30 min | 120 | ASlperi,<br>ASlcat | Both | LMM | -0.48 | 0.64 | — | 0.038 | <0.001 |
| S1B | 2 h | 120 | ASlperi,<br>ASlcat | Both | LMM | 3 | 0.0026 | 0.018 | 0.031 | 0.0057 |
| S1C | 30 min | 120 | $l_{ag}Frac$ | Both | LMM | -0.62 | 0.55 | — | 0.0015 | — |
| S1C | 2 h | 120 | $l_{ag}Frac$ | Both | LMM | 0.28 | 0.78 | — | <0.001 | — |
| S1D | 30 min | 120 | $d_{ag}$ | Both | LMM | 0.11 | 0.91 | — | <0.001 | — |
| S1D | 2 h | 120 | $d_{ag}$ | Both | LMM | 1.3 | 0.21 | — | 0.0035 | — |

**Table S10.** Summary of statistical output comparing the mean spine volume, ASI perimeter, length of astroglial surround, and astroglial distance to the ASI perimeter between control and cLTD conditions in the OML. **Statistical outcomes for comparisons visualized in Figure S2.** Cols 1-11 are as described for Table S9.

| Fig. | Time | $d_{ag}$<br>(nm) | Var, CoVar | Cluster | Test | Statistic | p (C) | $\eta_p^2$ (C) | R<sup>2</sup> | p (CoVar) |
| --- | --- | --- | --- | --- | --- | --- | --- | --- | --- | --- |
| S2A | 30 min | 120 | SpVol,<br>ASlcat | Both | LMM | -0.27 | 0.79 | — | 0.011 | 0.011 |
| S2A | 2 h | 120 | SpVol,<br>ASlcat | Both | LMM | -0.34 | 0.74 | — | 0.034 | <0.001 |
| S2B | 30 min | 120 | ASlperi,<br>ASlcat | Both | LMM | -0.54 | 0.6 | — | 0.022 | <0.001 |
| S2B | 2 h | 120 | ASlperi,<br>ASlcat | Both | LMM | -0.77 | 0.45 | — | 0.027 | <0.001 |
| S2C | 30 min | 120 | $l_{ag}Frac$ | Both | LMM | 0.95 | 0.34 | — | 0.0019 | — |
| S2C | 2 h | 120 | $l_{ag}Frac$ | Both | LMM | 0.64 | 0.54 | — | 0.0015 | — |
| S2D | 30 min | 120 | $d_{ag}$ | Both | LMM | -1.7 | 0.095 | — | 0.0057 | — |
| S2D | 2 h | 120 | $d_{ag}$ | Both | LMM | 0.14 | 0.89 | — | <0.001 | — |

**Table S11.** Summary of statistical output comparing the mean PSD area and PSD offset, and the relationship between these variables and astroglial apposition at the synapse, between control and LTP conditions in the MML. **Statistical outcomes for comparisons visualized in Figure S3.** Cols 1-3, 5-7, 9-11 are as described for Table S9. (Col 4) Response variable (Var) and covariate (CoVar): PSD area (PSDarea), PSD offset from the ASI center (PSDoffset), PAP present or absent from the ASI within 120 nm (ASIcat), length of astrogial surround at the ASI perimeter within 120 nm (l_ag_), astroglial distance to the ASI perimeter (d_ag_). If a significant interaction between the covariate and condition was detected, this is indicated by “(int).” (Col 8) p-value for condition effect, or for interaction effect if significant (as indicated in Col 4).

| Fig. | Time | $d_{ag}$<br>(nm) | Var, CoVar | Cluster | Test | Statistic | p (C) | $\eta_p^2$ (C) | R<sup>2</sup> | p (CoVar) |
| --- | --- | --- | --- | --- | --- | --- | --- | --- | --- | --- |
| S3A | 30 min | 120 | PSDarea,<br>ASlcat | Both | LMM | -1.4 | 0.19 | — | 0.052 | <0.001 |
| S3A | 2 h | 120 | PSDarea,<br>ASlcat | Both | LMM | 0.68 | 0.51 | — | 0.01 | 0.033 |
| S3B | 30 min | 120 | PSDoffset,<br>ASlcat | Both | LMM | -0.0065 | 0.99 | — | 0.0026 | 0.21 |
| S3B | 2 h | 120 | PSDoffset,<br>ASlcat | Both | LMM | 2.1 | 0.052 | — | 0.022 | 0.0087 |
| S3C | 30 min | 120 | PSDarea,<br>$l_{ag}$ | 1 | LMM | -2.9 | 0.012 | 0.025 | 0.21 | <0.001 |
| S3C | 30 min | 120 | PSDarea,<br>$l_{ag}$ | 2 | LMM | 0.66 | 0.52 | — | 0.078 | <0.001 |
| S3C | 2 h | 120 | PSDarea,<br>$l_{ag}$ (int) | 1 | LMM | 2.4 | 0.018 | — | 0.11 | — |
| S3C | 2 h | 120 | PSDarea,<br>$l_{ag}$ | 2 | LMM | -0.46 | 0.65 | — | 0.21 | <0.001 |
| S3D | 30 min | 120 | PSDarea,<br>$d_{ag}$ | 1 | LMM | -3.6 | 0.0033 | 0.037 | 0.072 | 0.6 |
| S3D | 30 min | 120 | PSDarea,<br>$d_{ag}$ | 2 | LMM | 0.41 | 0.69 | — | 0.047 | 0.0049 |
| S3D | 2 h | 120 | PSDarea,<br>$d_{ag}$ | 1 | LMM | -1.8 | 0.09 | — | 0.036 | 0.92 |
| S3D | 2 h | 120 | PSDarea,<br>$d_{ag}$ | 2 | LMM | -0.85 | 0.41 | — | 0.057 | 0.052 |
| S3E | 30 min | 120 | PSDoffset,<br>l<sub>ag</sub> | 1 | LMM | -0.27 | 0.79 | — | 0.024 | 0.0033 |
| S3E | 30 min | 120 | PSDoffset,<br>l<sub>ag</sub> | 2 | LMM | 1.1 | 0.27 | — | 0.013 | 0.31 |
| S3E | 2 h | 120 | PSDoffset,<br>l<sub>ag</sub> | 1 | LMM | -0.34 | 0.74 | — | 0.11 | <0.001 |
| S3E | 2 h | 120 | PSDoffset,<br>l<sub>ag</sub> (int) | 2 | LMM | -2.1 | 0.039 | — | 0.042 | — |
| S3F | 30 min | 120 | PSDoffset,<br>d<sub>ag</sub> (int) | 1 | LMM | -2.5 | 0.014 | — | 0.022 | — |
| S3F | 30 min | 120 | PSDoffset,<br>d<sub>ag</sub> | 2 | LMM | 0.71 | 0.48 | — | 0.019 | 0.16 |
| S3F | 2 h | 120 | PSDoffset,<br>d<sub>ag</sub> | 1 | LMM | -0.16 | 0.88 | — | 0.00092 | 0.6 |
| S3F | 2 h | 120 | PSDoffset,<br>d<sub>ag</sub> | 2 | LMM | 0.72 | 0.47 | — | 0.0076 | 0.45 |
| S3G | 30 min | 120 | PSDoffset,<br>PSDarea | 1 | LMM | 0.13 | 0.9 | — | 0.052 | <0.001 |
| S3G | 30 min | 120 | PSDoffset,<br>PSDarea | 2 | LMM | 0.92 | 0.36 | — | 0.035 | 0.033 |
| S3G | 2 h | 120 | PSDoffset,<br>PSDarea | 1 | LMM | 0.42 | 0.68 | — | 0.035 | <0.001 |
| S3G | 2 h | 120 | PSDoffset,<br>PSDarea | 2 | LMM | 0.46 | 0.65 | — | 0.0025 | 0.88 |

**Table S12.** Summary of statistical output comparing the ultrastructure of ag-proximal (d_ag-PSD_ < d_ASI-PSD_) versus ag-distal (d_ag-PSD_ ≥ d_ASI-PSD_) synapses between control and LTP conditions in the MML. **Statistical outcomes for comparisons visualized in Figure S4.** Cols 1-3, 5, 8-11 are as described for Table S9. (Col 4) Response variable (Var) and covariate (CoVar): ag-proximal versus ag-distal synapses (offsetCat), spine volume (spVol), PSD area (PSDarea). (Col 6) Statistical test: Chi-squared test (Χ^2^), Linear mixed model (LMM). (Col 7) Test statistic: χ² for χ² test, t for LMM.

| Fig. | Time | $d_{ag}$<br>(nm) | Var, CoVar | Cluster | Test | Statistic | p (C) | $\eta_p^2$ (C) | R<sup>2</sup> | p (CoVar) |
| --- | --- | --- | --- | --- | --- | --- | --- | --- | --- | --- |
| S4A | 30 min | 120 | OffsetCat | Both | $\chi^2$ | 0.1 | 0.75 | — | — | — |
| S4A | 2 h | 120 | OffsetCat | Both | $\chi^2$ | 0.74 | 0.39 | — | — | — |
| S4B | 30 min | 120 | SpVol,<br>OffsetCat | Both | LMM | 0.39 | 0.7 | — | 0.0084 | 0.031 |
| S4B | 2 h | 120 | SpVol,<br>OffsetCat | Both | LMM | 4.5 | <0.001 | 0.044 | 0.061 | 0.022 |
| S4C | 30 min | 120 | ASlperi,<br>OffsetCat | Both | LMM | -0.48 | 0.63 | — | 0.0078 | 0.074 |
| S4C | 2 h | 120 | ASlperi,<br>OffsetCat | Both | LMM | 3.3 | <0.001 | 0.024 | 0.042 | 0.0019 |
| S4D | 30 min | 120 | PSDarea,<br>OffsetCat | Both | LMM | -1.3 | 0.23 | — | 0.0084 | 0.22 |
| S4D | 2 h | 120 | PSDarea,<br>OffsetCat | Both | LMM | 1.1 | 0.3 | — | 0.019 | 0.006 |

**Table S13:** Summary of statistical output comparing the mean PSD area and PSD offset, and the relationship between these variables and astroglial apposition at the synapse, between control and cLTD conditions in the OML. **Statistical outcomes for comparisons visualized in Figure S5.** Cols 1-3, 5-7, 9-11 are as described for Table S9. (Col 4) Response variable (Var) and covariate (CoVar): PSD area (PSDarea), PSD offset from the ASI center (PSDoffset), PAP present or absent from the ASI within 120 nm (ASIcat), length of astrogial surround at the ASI perimeter within 120 nm (l_ag_), astroglial distance to the ASI perimeter (d_ag_). If a significant interaction between the covariate and condition was detected, this is indicated by “(int).” (Col 8) p-value for condition effect, or for interaction effect if significant (as indicated in Col 4).

| Fig. | Time | $d_{ag}$<br>(nm) | Var, CoVar | Cluster | Test | Statistic | p (C) | $\eta_p^2$ (C) | R<sup>2</sup> | p (CoVar) |
| --- | --- | --- | --- | --- | --- | --- | --- | --- | --- | --- |
| S5A | 30 min | 120 | PSDarea,<br>ASlcat | Both | LMM | -0.011 | 0.99 | — | 0.016 | 0.0035 |
| S5A | 2 h | 120 | PSDarea,<br>ASlcat | Both | LMM | -0.89 | 0.37 | — | 0.026 | <0.001 |
| S5B | 30 min | 120 | PSDoffset,<br>ASlcat | Both | LMM | -2 | 0.07 | — | 0.028 | 0.096 |
| S5B | 2 h | 120 | PSDoffset,<br>ASlcat | Both | LMM | -0.42 | 0.68 | — | 0.0052 | 0.12 |
| S5C | 30 min | 120 | PSDarea,<br>$l_{ag}$ | 1 | LMM | -0.73 | 0.48 | — | 0.09 | <0.001 |
| S5C | 30 min | 120 | PSDarea,<br>$l_{ag}$ | 2 | LMM | 0.56 | 0.59 | — | 0.1 | 0.0099 |
| S5C | 2 h | 120 | PSDarea,<br>$l_{ag}$ | 1 | LMM | -0.97 | 0.35 | — | 0.061 | <0.001 |
| S5C | 2 h | 120 | PSDarea,<br>$l_{ag}$ | 2 | LMM | -0.45 | 0.66 | — | 0.25 | <0.001 |
| S5D | 30 min | 120 | PSDarea,<br>$d_{ag}$ | 1 | LMM | -0.91 | 0.38 | — | 0.0036 | 0.71 |
| S5D | 30 min | 120 | PSDarea,<br>$d_{ag}$ | 2 | LMM | 0.15 | 0.88 | — | 0.045 | 0.11 |
| S5D | 2 h | 120 | PSDarea,<br>$d_{ag}$ | 1 | LMM | -0.82 | 0.42 | — | 0.0018 | 0.87 |
| S5D | 2 h | 120 | PSDarea,<br>$d_{ag}$ | 2 | LMM | -0.91 | 0.38 | — | 0.041 | 0.2 |
| S5E | 30 min | 120 | PSDoffset,<br>$l_{ag}$ | 1 | LMM | -1.9 | 0.082 | — | 0.092 | <0.001 |
| S5E | 30 min | 120 | PSDoffset,<br>$l_{ag}$ | 2 | LMM | -1.3 | 0.19 | — | 0.054 | 0.35 |
| S5E | 2 h | 120 | PSDoffset,<br>l<sub>ag</sub> | 1 | LMM | -0.86 | 0.4 | — | 0.052 | <0.001 |
| S5E | 2 h | 120 | PSDoffset,<br>l<sub>ag</sub> (int) | 2 | LMM | -2.4 | 0.021 | — | 0.11 | — |
| S5F | 30 min | 120 | PSDoffset,<br>d<sub>ag</sub> | 1 | LMM | -2 | 0.064 | — | 0.034 | 0.17 |
| S5F | 30 min | 120 | PSDoffset,<br>d<sub>ag</sub> | 2 | LMM | -1.5 | 0.14 | — | 0.052 | 0.38 |
| S5F | 2 h | 120 | PSDoffset,<br>d<sub>ag</sub> | 1 | LMM | -0.8 | 0.43 | — | 0.0024 | 0.69 |
| S5F | 2 h | 120 | PSDoffset,<br>d<sub>ag</sub> (int) | 2 | LMM | 3 | 0.0041 | — | 0.11 | — |
| S5G | 30 min | 120 | PSDoffset,<br>PSDarea | 1 | LMM | -1.7 | 0.11 | — | 0.047 | 0.0016 |
| S5G | 30 min | 120 | PSDoffset,<br>PSDarea | 2 | LMM | -1.5 | 0.13 | — | 0.04 | 0.83 |
| S5G | 2 h | 120 | PSDoffset,<br>PSDarea (int) | 1 | LMM | -4 | <0.001 | — | 0.049 | — |
| S5G | 2 h | 120 | PSDoffset,<br>PSDarea | 2 | LMM | -0.11 | 0.92 | — | 0.04 | 0.084 |

**Table S14.** Summary of statistical output comparing the ultrastructure of ag-proximal (d_ag-PSD_ < d_ASI-PSD_) versus ag-distal (d_ag-PSD_ ≥ d_ASI-PSD_) synapses between control and cLTD conditions in the OML. **Statistical outcomes for comparisons visualized in Figure S6.** Cols 1-3, 5, 8-11 are as described for Table S9. (Col 4) Response variable (Var) and covariate (CoVar): ag-proximal versus ag-distal synapses (offsetCat), spine volume (spVol), PSD area (PSDarea). (Col 6) Statistical test: Chi-squared test (Χ^2^), Linear mixed model (LMM). (Col 7) Test statistic: χ² for χ² test, t for LMM.

| Fig. | Time | $d_{ag}$<br>(nm) | Var, CoVar | Cluster | Test | Statistic | p (C) | $\eta_p^2$ (C) | R<sup>2</sup> | p (CoVar) |
| --- | --- | --- | --- | --- | --- | --- | --- | --- | --- | --- |
| S6A | 30 min | 120 | prox vs. dist | Both | $\chi^2$ | 0.16 | 0.69 | — | — | — |
| S6A | 2 h | 120 | prox vs. dist | Both | $\chi^2$ | 0.87 | 0.35 | — | — | — |
| S6B | 30 min | 120 | SpVol,<br>OffsetCat | Both | LMM | -0.2 | 0.85 | — | <0.001 | 0.69 |
| S6B | 2 h | 120 | SpVol,<br>OffsetCat | Both | LMM | -0.033 | 0.97 | — | <0.001 | 0.55 |
| S6C | 30 min | 120 | ASlperi,<br>OffsetCat | Both | LMM | -0.62 | 0.54 | — | 0.0021 | 0.9 |
| S6C | 2 h | 120 | ASlperi,<br>OffsetCat | Both | LMM | -0.31 | 0.76 | — | 0.0019 | 0.41 |
| S6D | 30 min | 120 | PSDarea,<br>OffsetCat | Both | LMM | -0.17 | 0.87 | — | <0.001 | 0.79 |
| S6D | 2 h | 120 | PSDarea,<br>OffsetCat | Both | LMM | -0.57 | 0.57 | — | <0.001 | 0.95 |

---

**Figure S1.**
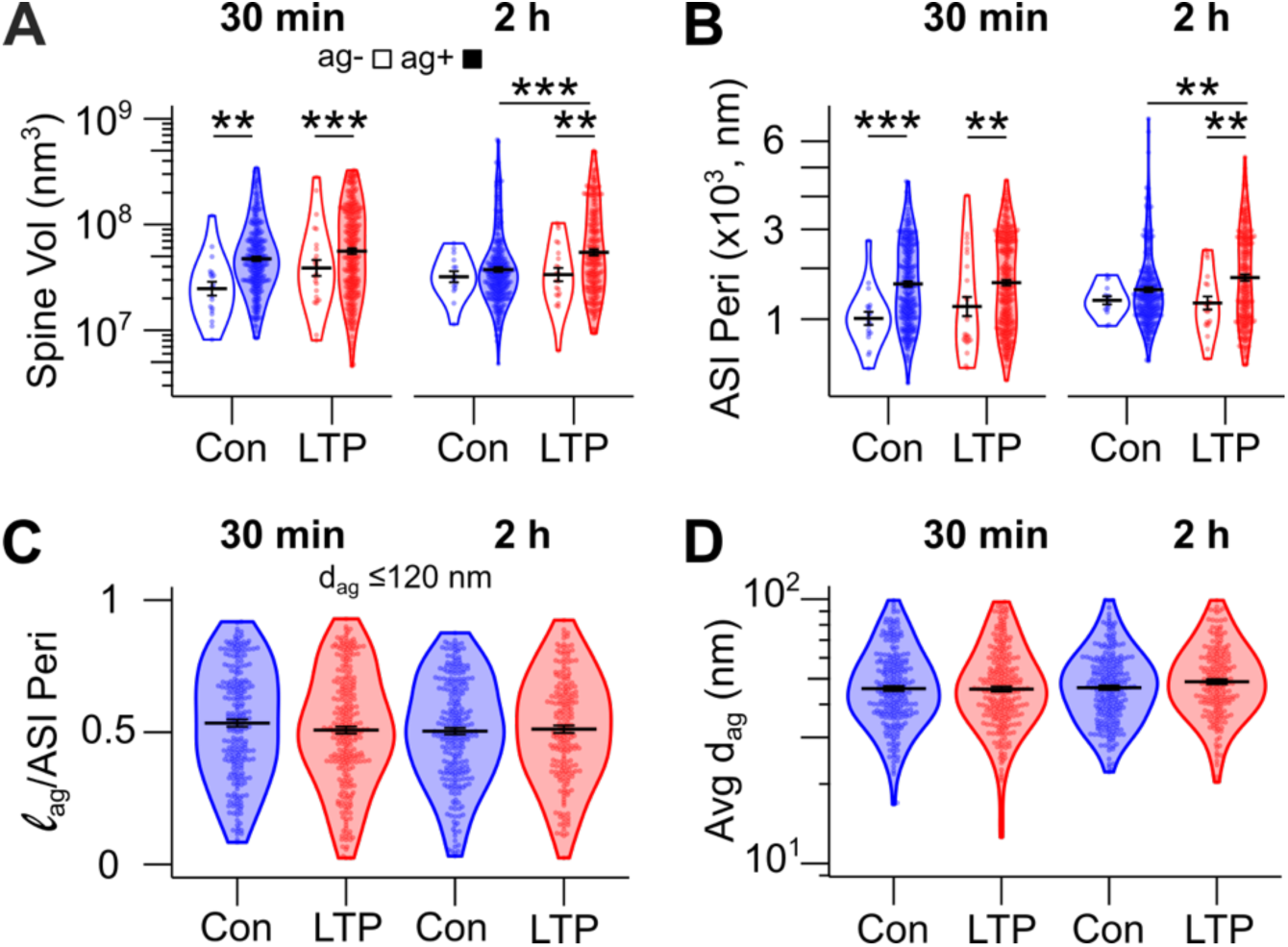
Mean spine size and astroglia apposition at the ASI perimeter during LTP. **(A)** Spine volume (log-scale y-axis) and **(B)** ASI perimeter (square-root scale on y-axis) for ASI^ag-^ (white, d_ag_ > 120 nm) and ASI^ag+^ (sky blue, d_ag_ ≤ 120 nm) synapses. **(C)** Fraction of ASI perimeter surrounded by astroglia within 120 nm (l_ag_/ASI perimeter) and **(D)** the average (avg) distance from the ASI perimeter to the nearest astroglia (d_ag_, log-scale axis) for synapses with astroglia within 120 nm of the ASI perimeter. In all panels, violin plots show the data distribution, overlaid with individual data points (beeswarm plots), group means (horizontal black lines), and standard errors (black error bars). Data from control (blue) and LTP (red) MML synapses at 30 minutes (left) and 2 hours (right) after LTP induction are shown. *p < 0.05, **p < 0.01, ***p < 0.001. (See Table S9 for statistical details.)

**Figure S2.**
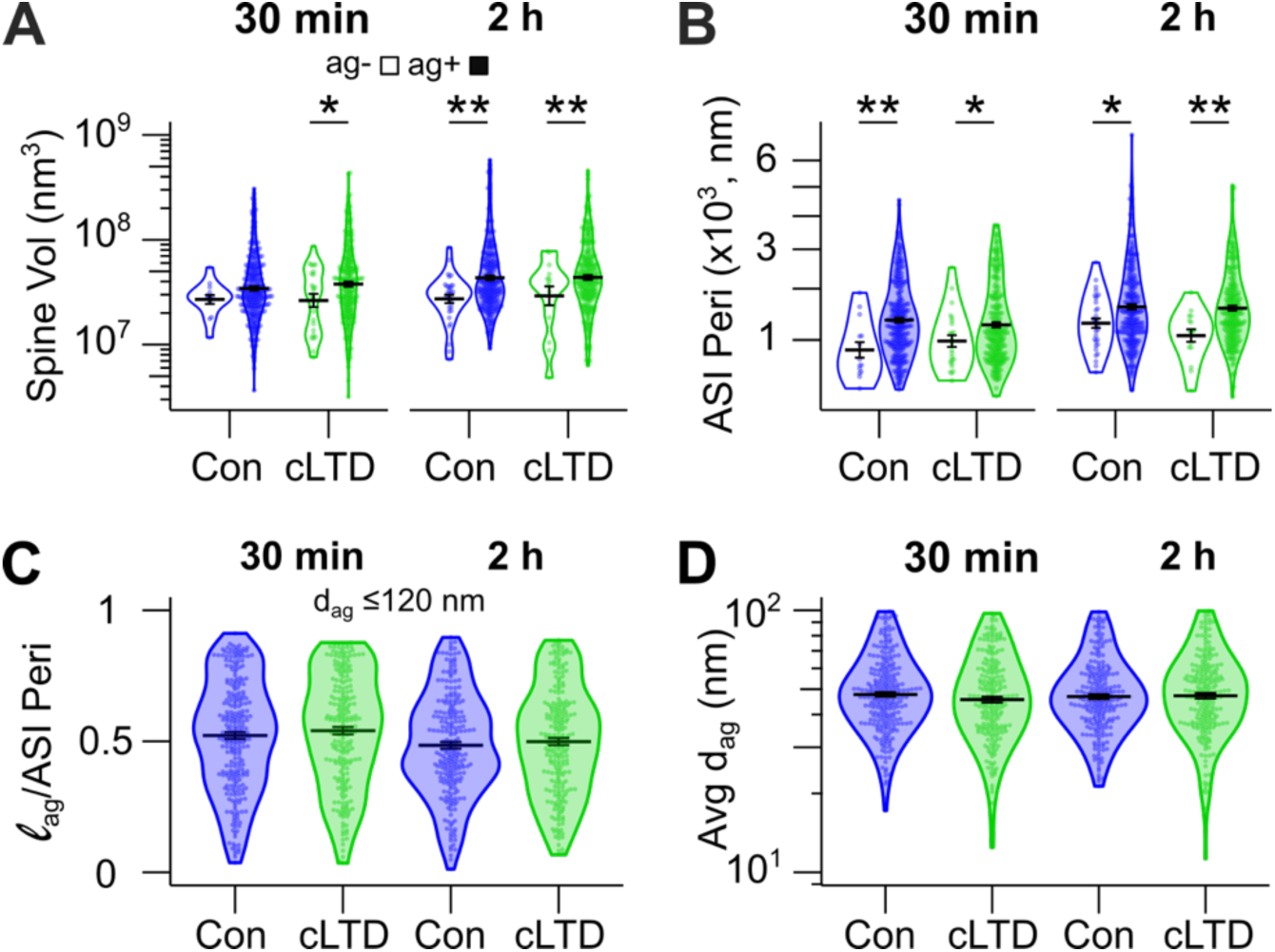
Mean spine size and astroglia apposition at the ASI perimeter during cLTD. **(A)** Spine volume (log-scale y-axis) and **(B)** ASI perimeter (square-root scale y-axis) for ASI^ag-^ (white, d_ag_ > 120 nm) and ASI^ag+^ (sky blue, d_ag_ ≤ 120 nm) synapses. **(C)** Fraction of ASI perimeter surrounded by astroglia within 120 nm (l_ag_/ASI perimeter) and **(D)** the average (avg) distance from the ASI perimeter to the nearest astroglia (d_ag_, log-scale axis) for synapses with astroglia within 120 nm of the ASI perimeter. In all panels, violin plots show the data distribution, overlaid with individual data points (beeswarm plots), group means (horizontal black lines), and standard errors (black error bars). Data from control (blue) and cLTD (green) OML synapses at 30 minutes (left) and 2 hours (right) after cLTD induction are shown. *p < 0.05, **p < 0.01. (See Table S10 for statistical details.)

**Figure S3.**
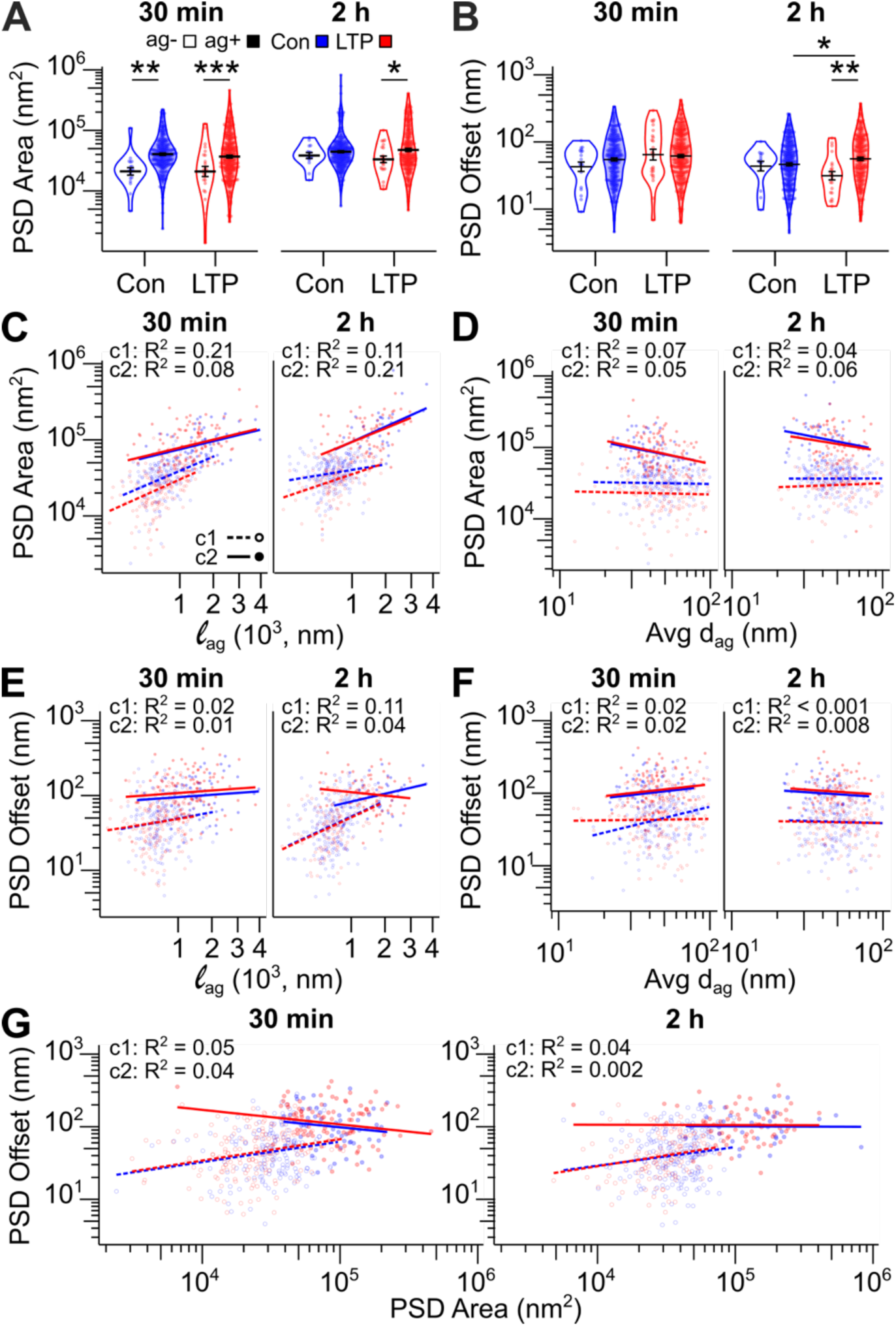
Impact of astroglia apposition at the ASI perimeter on mean PSD area and PSD offset during control and LTP. **(A)** Violin plots of PSD area (log-scale axis) and **(B)** PSD offset (log-scale axis) for ASI^ag-^ (white, d_ag_ > 120 nm) and ASI^ag+^ (sky blue, d_ag_ ≤ 120 nm) synapses, overlaid with individual data points (beeswarm plots), group means (horizontal black lines), and standard errors (black error bars). **(C)** Regression plot of PSD area (log-scale y-axis) versus the length of the ASI perimeter surrounded by astroglia within 120 nm (l_ag_ square-root x-axis) and **(D)** the average (avg) distance between astroglia processes and the ASI perimeter (d_ag_ log-scale x-axis) for c1 and c2 synapses. **(E)** Regression plot of PSD offset (log-scale y-axis) versus l_ag_ based on d_ag_ ≤ 120 nm (square-root x-axis), **(F)** average d_ag_ (log-scale x-axis), and **(G)** PSD area (log-scale x-axis) for c1 and c2 synapses. In A-G, data from control (blue) and LTP (red) MML synapses with astroglia within 120 nm of the ASI perimeter at 30 minutes (left) and 2 hours (right) after LTP induction are shown, with only synapses with d_ag_ ≤ 120 nm analyzed in C-G. *p < 0.05, **p < 0.01, ***p < 0.001. (See Table S11 for statistical details.)

**Figure S4.**
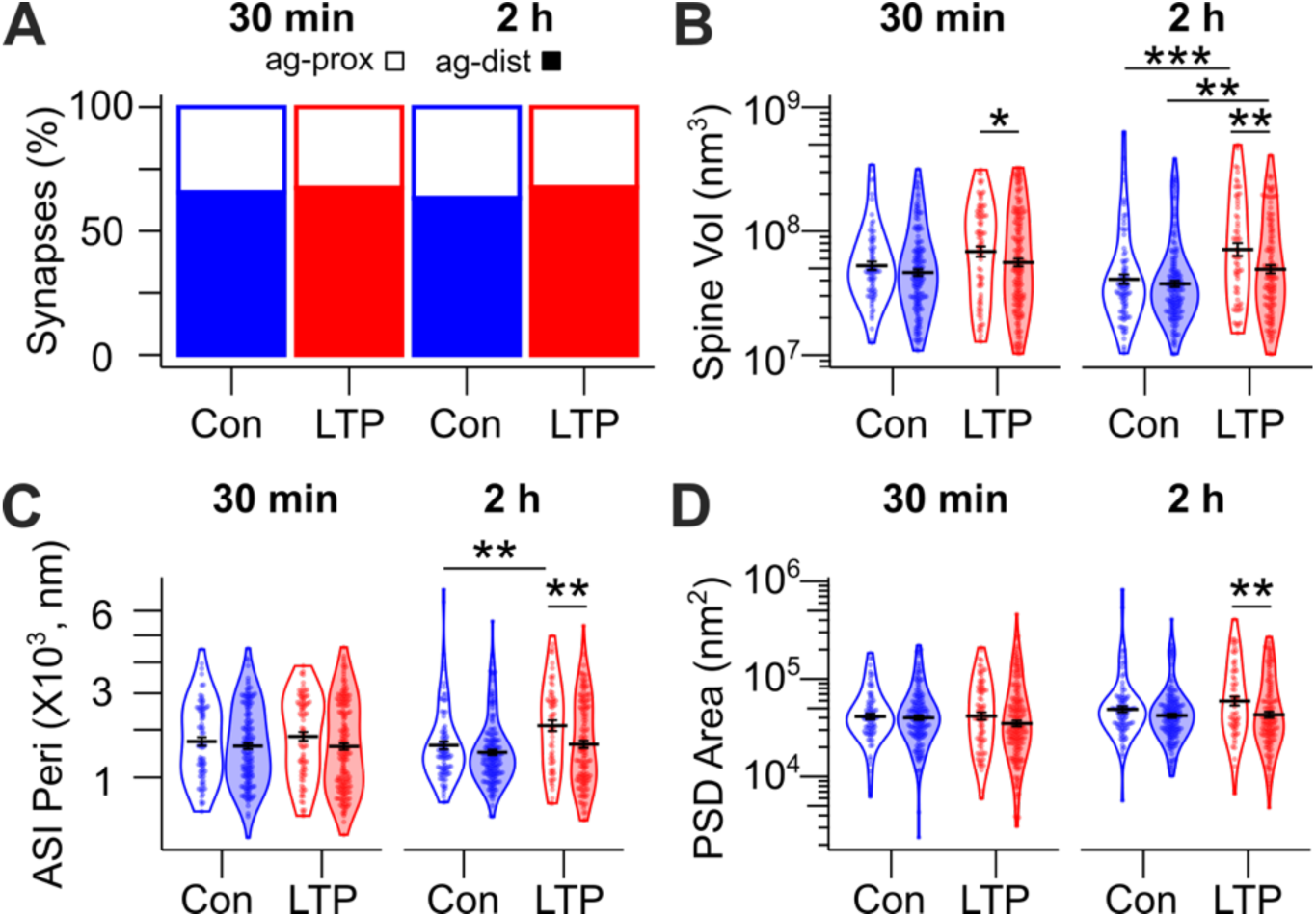
Comparison of ag-proximal (d_ag-PSD<_d_ASI-PSD_) and ag-distal (d_ag-PSD_ ≥ _dASI-PSD_) synapses during control and LTP. MML synapses with astroglia within 120 nm of the synapse were categorized based on PSD offset: PSD positioned relatively closer to (ag-proximal) or farther from (ag-distal) astroglia apposition at the ASI perimeter. **(A)** Stacked bar graph of the relative percentage of ag-proximal versus ag-distal synapses. Numbers above bars indicate total synapse counts. **(B)** Violin plots of spine volume (log-scale axis), **(C)** ASI perimeter (square-root axis), and **(C)** PSD area (log-scale axis) for ag-proximal and ag-distal MML synapses, overlaid with individual data points (beeswarm plots), group means (horizontal black lines), and standard errors (black error bars). In A-D, only data from control (blue) and LTP (red) MML synapses with d_ag_ ≤ 120 nm at 30 minutes (left) and 2 hours (right) after LTP induction are shown. *p < 0.05, **p < 0.01, ***p < 0.001. (See Table S12 for statistical details.)

**Figure S5.**
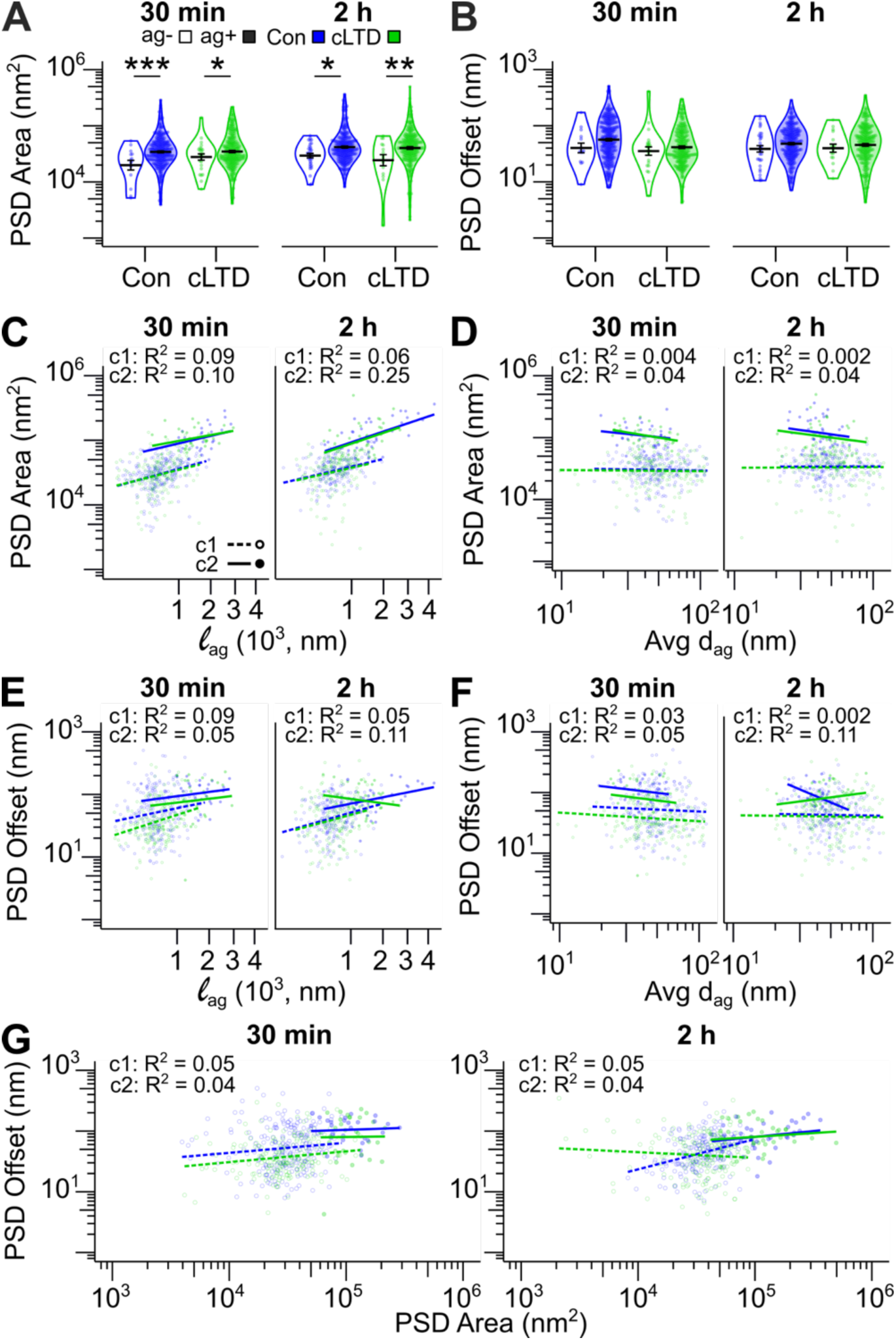
Impact of astroglia apposition at the ASI perimeter on mean PSD area and PSD offset during control and cLTD. **(A)** Violin plots of PSD area (log-scale axis) and **(B)** PSD offset (log-scale axis) for ASI^ag-^ (white, d_ag>_120 nm) and ASI^ag+^ (sky blue, d_ag_ ≤ 120 nm) synapses, overlaid with individual data points (beeswarm plots), group means (horizontal black lines), and standard errors (black error bars). **(C)** Regression plot of PSD area (log-scale y-axis) versus the length of the ASI perimeter surrounded by astroglia within 120 nm (l_ag_ plotted on square-root x-axis) and **(D)** the average (avg) distance between astroglia processes and the ASI perimeter (d_ag_, log-scale x-axis) for c1 and c2 synapses. **(E)** Regression plot of PSD offset (log-scale y-axis) versus l_ag_ based on d_ag_ ≤ 120 nm (square-root x-axis), **(F)** the average d_ag_ (log-scale x-axis), and **(G)** PSD area (log-scale x-axis) for c1 and c2 synapses. In A-G, data from control (blue) and cLTD (green) OML synapses with astroglia within 120 nm of the ASI perimeter at 30 minutes (left) and 2 hours (right) after cLTD induction are shown, with only synapses with d_ag_ ≤ 120 nm analyzed in C-G. *p < 0.05, **p < 0.01, ***p < 0.001. (See Table S13 for statistical details.)

**Figure S6.**
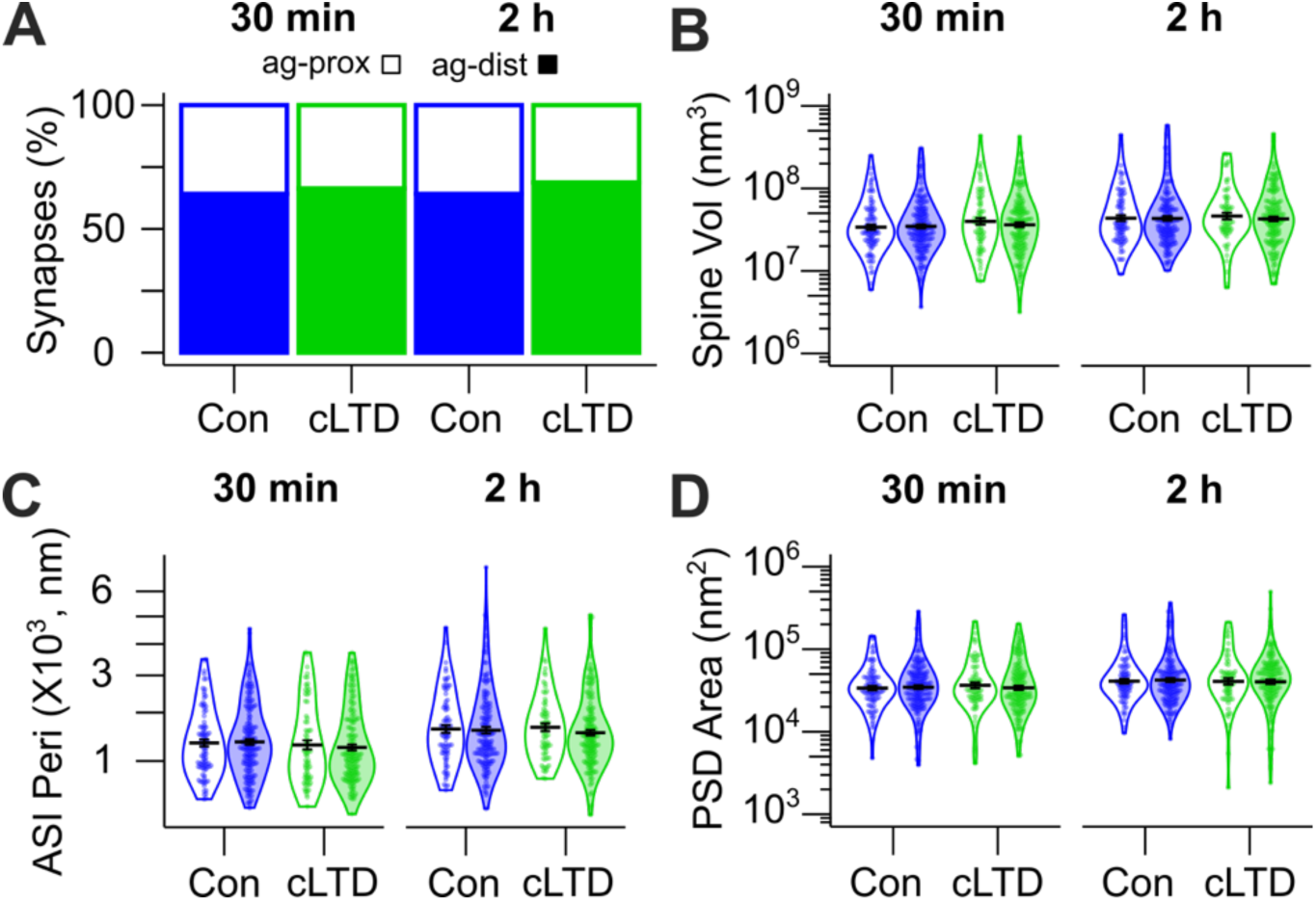
Comparison of ag-proximal (d_ag-PSD<_d_ASI-PSD_) and ag-distal (d_ag-PSD_ ≥_dASI-PSD_) synapses during control and cLTD in the OML. OML synapses with astroglia within 120 nm of the synapse were categorized based on PSD offset: PSD positioned relatively closer to (ag-proximal) or farther from (ag-distal) astroglia apposition at the ASI perimeter. **(A)** Stacked bar graph of the relative percentage of ag-proximal versus ag-distal synapses. Numbers above bars indicate total synapse counts. **(B)** Violin plots of spine volume (log-scale axis), **(C)** ASI perimeter (square-root axis), and **(D)** PSD area (log-scale axis) for ag-proximal and ag-distal OML synapses, overlaid with individual data points (beeswarm plots), group means (horizontal black lines), and standard errors (black error bars). In A-D, only data from control (blue) and cLTD (green) OML synapses with d_ag_ ≤ 120 nm at 30 minutes (left) and 2 hours (right) after cLTD induction are shown. (See Table S14 for statistical details.)

## Notes

https://doi.org/10.18738/T8/S8DT5E

### Competing Interest Statement

The authors have declared no competing interest.

